# A computational study to assess the pathogenicity of single or combinations of missense variants on respiratory Complex I

**DOI:** 10.1101/2023.12.11.571090

**Authors:** Laura Rigobello, Francesca Lugli, Leonardo Caporali, Alessio Bartocci, Jacopo Fadanni, Francesco Zerbetto, Luisa Iommarini, Valerio Carelli, Anna Maria Ghelli, Francesco Musiani

**Affiliations:** Department of Pharmacy and Biotechnology, University of Bologna, Italy; Department of Chemistry “Giacomo Ciamician”, University of Bologna, Italy; IRCCS Istituto delle Scienze Neurologiche di Bologna, Programma di Neurogenetica, Bologna, Italy; Department of Biomedical and Neuromotor Sciences (DIBINEM), University of Bologna, Italy; Department of Physics, University of Trento, Italy; INFN-TIFPA, Trento Institute for Fundamental Physics and Applications, Trento Italy

## Abstract

Variants found in the respiratory complex I (CI) subunit genes encoded by mitochondrial DNA can cause severe genetic diseases. However, it is difficult to establish *a priori* whether a single or a combination of CI variants may impact oxidative phosphorylation. Here we propose a computational approach based on coarse-grained molecular dynamics simulations. One of the primary CI variants (m.14484T>C/*MT-ND6*) associated with the Leber hereditary optic neuropathy was used as a test case. This variant was investigated alone or in combination with two additional rare CI variants whose role remains uncertain. We found that the primary variant stiffens CI dynamics in the crucial E-channel region and that one of the other variants, located in the vicinity of the primary one, further worsens the stiffening. This approach may be extended to other variants candidate to exert a pathogenic impact on CI function, or to investigate the interaction of multiple variants.

**Teaser:** Molecular dynamics is able to predict the functional impact of variants hitting respiratory complex I mitochondrial genes.

## INTRODUCTION

Respiratory complex I (CI, EC 1.6.5.3) is the largest enzyme of the mitochondrial respiratory chain (*1, 2*). In mammals, CI is under a dual genetic control as out of its 44 subunits, 7 (namely ND1-6 and ND4L) are encoded by the multicopy mitochondrial genome (mtDNA), whereas the remaining ones and all the assembly factors needed to its biogenesis are encoded by the nuclear genome (nDNA) (*3–5*). Pathogenic variants affecting CI subunits have been described in both nDNA and mtDNA-encoded components, leading to variable phenotypic expression. The latter ranges from severe and lethal infantile encephalopathy known as Leigh syndrome to adult-onset milder phenotypes such as Leber’s hereditary optic neuropathy (LHON). These diseases all characterized by CI dysfunction, which is the most frequent amongst metabolic disorders in humans (*6*). LHON was the first human disease to be associated with mtDNA pathogenic variants (*7*) and is characterized by subacute loss of central vision in one eye, followed, usually after 3-6 months, by the involvement of the fellow eye due to retinal ganglion cells degeneration (*8*). LHON prevalence is currently estimated one in 25,000-50,000 people in European population, thus resulting the most frequent mitochondrial disease (*6*). About 90% of LHON cases are induced by the three primary mtDNA point mutations, namely m.3460G>A/*MT-ND1*, m.11778G>A/*MT-ND4*, and m.14484T>C/*MT-ND6* (*9*). The remaining 10% of patients harbors one or more “rare mutations” in mtDNA (*10*) or unique combination of missense variants (*11*). More recently, recessive mutations, yet affecting in most cases CI, have been also found to underlie LHON (*6*).

The pathogenic variants established as risk factor for LHON are characterized by incomplete penetrance and may need either the combination with further genetic factors, or specific environmental exposures, to express the disease. Amongst the supplementary genetic factors, the most studied is the mtDNA itself, characterized by an intrinsic variability, which has been selected in population-specific haplotypes by evolutionary pressure (*12*). For example, the missense m.T14484C/*MT-ND6* LHON variant needs to arise in the context of haplogroup J, since other missense mtDNA variants defining this haplogroup increase its pathogenic potential (*13*). In fact, the m.T14484C/*MT-ND6* variant may be recognized in population genetics surveys, reaching a frequency as high as 1 in 800 individuals in absence of any pathology (*14, 15*) when preferentially associated with H or U haplogroups, or may be found as diagnostic marker of LHON in pedigrees with one or more affected individuals along the maternal line. On the opposite end, specific combinations in single pedigrees have been shown to increase greatly the disease penetrance up to almost complete penetrance as in the case of a Chinese family presenting the co-existence of the primary m.14484T>C/*MT-ND6* pathogenic variant with the rare m.10680G>A/*MT-ND4L* and a third m.13942A>G/*MT-ND5* variants (*16*). Hence, the m.14484T>C/*MT-ND6* variant is a good example of the challenges encountered in attributing a specific pathogenic role to mtDNA variants, as since its initial identification the role played in LHON pathogenesis was ambiguous (*17, 18*). This ambiguity is mainly generated by its frequency in the general population and its weak evidence of functional impact on CI function.

Overall, the LHON pathogenic mutations affecting CI induce a decrease of oxidative phosphorylation driven by CI-linked substrates and is associated with an increase of reactive oxygen species production that, in turn, enhance the sensitivity of the cells to apoptosis (*19*). Despite the general biochemical defect induced by LHON mutations is quite known, the molecular changes triggered by mutated amino acids on CI mechanism are still unknown. However, thanks to the greatly improved understanding of the mechanistic function of CI, gained mainly after the release of cryo-EM and X-ray crystal structures (*1, 2, 20–25*), we can now attempt to better define how mtDNA variants may impinge on CI structure and function, in particular when occurring in specific combinations. The CI structure is commonly divided into three functional modules (Fig. 1): *i*) the NADH oxidation module (N module) that catalyzes the electron transfer through a chain of iron-sulfur clusters; *ii*) the ubiquinone reduction module (Q module); and *iii*) the proton translocation module (P module) that induces proton translocation (*26*). In particular, the P-module is formed by the mtDNA-encoded subunits and is also involved in the formation of the binding site of the ubiquinone (called Q-cavity), located at the interface between the Q and P module (Fig. 1) (*1, 2, 21–25*). Further, subunits ND1, ND3, ND6 and ND4L form the proximal part of the P module at the interface with the Q module (*22–24, 27*) and create together the E-channel, a region containing several conserved glutamates (Fig. 1) (*28, 29*). Subunits ND2, ND4 and ND5 form the remaining functional core of the P module. Importantly, in the recently released ovine and *Escherichia coli* structures obtained under a range of redox conditions, including catalytic turnover, CI appears in two distinguishable conformations (open and close as proposed by Sazanov and coworkers, see ref. (*30*) for a discussion regarding the different nomenclatures of CI conformations) (*20, 23, 24, 30*). Compared to the close counterpart, in the open conformation the angle formed by the matrix arm (Q plus N modules) with the P module is wider by approximately 7 degrees. Moreover, the open conformation shows some partially disordered regions that are folded in the close conformation. Finally, some regions change their structure. Specifically, in the open conformation the third trans-membrane helix (TMH3) of ND6 forms a π-bulge (Fig. 1), which breaks the hydrophilic axis in the E-channel region and is proposed to be crucial for CI mechanism, preventing a futile proton leaking towards the Q-cavity (*20, 27*). Despite the structural and biochemical advances, the mechanism of the proton pumping coupled to the reduction of ubiquinone is still debated (*27, 31, 32*). The complexity of CI structure implies that many amino acids have a crucial role in the conformational changes required for its activity. Therefore, it is conceivable that the combination of amino acid variants, which alone have negligible effect, can affect the activity of CI inducing LHON disease in some pedigrees. Here, we applied the molecular dynamics (MD) technique with coarse-grained (CG) approach for the study of the effect of mtDNA variants on CI structure. In particular, we investigated the effect of the puzzling primary pathogenic m.14484T>C *MT-ND6* (p.M64V/ND6) variant alone or in combination with the rare m.10680G>A/*MT-ND4L* (p.A71T/ND4L) and m.13942A>G/*MT-ND5* (p.T536A/ND5) variants, providing a “proof of principle” that this computational method may be useful for predicting the possible pathogenic effect of specific variant combination. From now on, the notation reporting the mutation on the protein translate will be used.

**Fig. 1.**
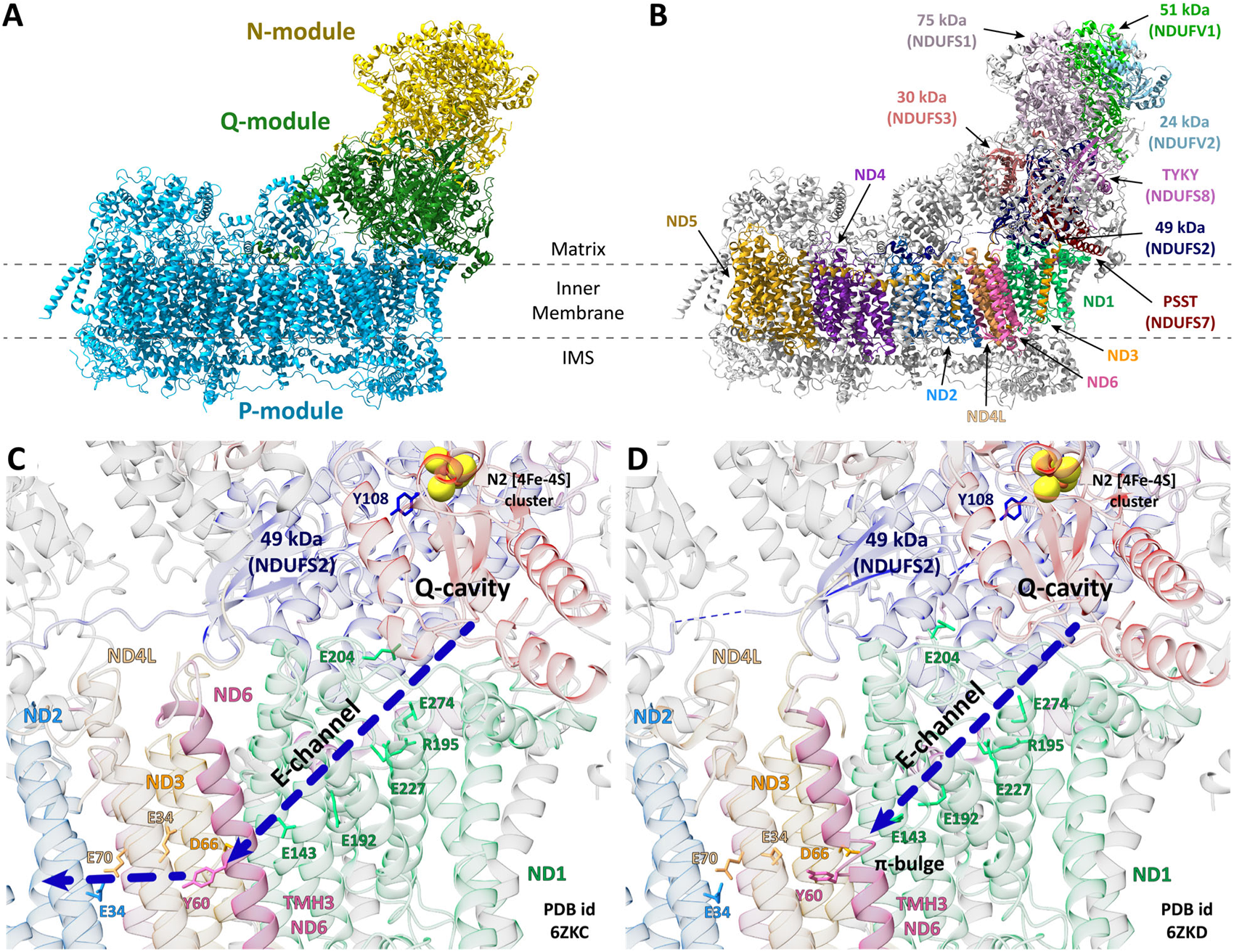
CI general structure and subunit organization. (**A** and **B**) Cartoon representation of the ovine CI structure (PDB id: 6ZKD) across the inner membrane. In panel **A** the N, Q and P module subdivision is reported, while in panel **B** the core subunits are highlighted with different colors (the human gene names are provided in brackets), while the supernumerary subunits are in grey. (**C** and **D**) Detail of the Q-cavity and E-channel regions in the close (**C**, PDB id 6ZKC) and open conformation (**D**, PDB id 6ZKD). The cartoons are colored according to panel **B**. The residues forming the E-channel or with an important role for the CI function are in sticks, while the N2 [4Fe-4S] cluster is reported in spheres colored according to the atom type. ND6 TMH3 and the π-bulge in the open conformation are highlighted, as well as the position of the Q-cavity and of the E-channel. The direction of the proton wire is schematized by dashed blue arrows.

### Rationale of the computational approach

Mammalian CI is a large protein complex, then a fully atomistic MD simulation including a physiological membrane and explicit treatment of the solvent implies the use of ca. 900,000 atoms and a tremendous computational effort (*33, 34*). The aims of our approach are *i*) to reduce the computational cost with respect to atomistic simulations and *ii*) to find the minimal CI structural elements required for the observation of dynamic effects attributable to pathogenic mutations. Regarding the first aim, the use of classical MD simulations with CG models preserving chemical specificity, such as the Martini force-field (*35, 36*), has emerged as powerful strategy to tackle temporal evolution of biomolecular systems (*37–39*). The use of a CG model reduces the number of particles in the simulated systems with a consequent decrease in the computational cost by 2-3 orders of magnitude compared to the atomistic models (*40*). Moreover, the Martini force field has been already successfully used on CI from *Thermus thermophilus* (*41*). Regarding the second goal, the MD simulations were accomplished on the structures comprising the ovine P module and the NDUFS2 subunit (49 kDa subunit in the ovine gene nomenclature, see Table S1, PDB IDs 6ZKA and 6ZKB) (*20*). NDUFS2 structure was partially reconstructed through homology modelling, together with other not solved regions of the P module (see Table S2 and Fig. S1). The simulated structures will be called P+ module hereafter to highlight the presence of the NDUFS2 subunit bound to the P module. The MD simulations were initially conducted on the structures of the P+ module from *Ovis aries* in both open and closed conformation. In addition to having a smaller number of atoms with respect to the complete CI structure, the P+ module has fewer technical difficulties because it does not bind the cofactors, that are in the N and Q modules.

The simulations performed on *wild type* ovine structures were used as control sample, while the simulations of the mutated ovine systems were not performed because some of the residues involved in the studied mutations are not conserved in *O. aries*. We then performed a larger set of simulations on homology models of the human *wild type* and mutated P+ modules in both open and close conformation. Homology modelling was used to obtain good quality human models because the available experimental human CI structures have a lower resolution than the ovine structures and human and sheep sequences have high sequence identity (Table S3). Moreover, the experimental human CI structures were obtained in only one conformation (namely, the open one). Each system was inserted in a double layer membrane resembling the human inner mitochondrial membrane and solvated in a solution at physiological ionic strength (see Materials and Methods section for details). The simulated systems will be labeled hereafter with the letter “O” for ovine and “H” for human structures. The endings “op” and “cl” will identify the open and close conformation, respectively.

The simulations consist of: *i*) three 8 µs-long benchmark simulations on the open and close ovine P+ module structures (wtO^op^ and wtO^cl^ hereafter, respectively); ii) three 16 µs-long simulations performed on the open and close model structures of the human P+ module (wtH^op^ and wtH^cl^); and *iii*) three 16 µs-long simulations performed on the single p.M64V/ND6 mutant (m1) and on the triple p.M64V/ND6 – p.A71T/ND4L – p.T536A/ND5 mutant (m3). Again, all the simulations were performed on the model structures of the open and close human P+ module (namely m1H^op^, m1H^cl^, m3H^op^ and m3H^cl^). The use of three replicas for each system is motivated by the observation that several simulations are more efficient for conformational space sampling than a single long trajectory (*42*). A total of 336 µs of plain MD simulations were carried out.

## RESULTS

### The P+ module is stable in the microsecond time scale

Visual inspection revealed that there are no global unfolding processes throughout the simulation time. To analyze the overall behavior, the root mean square deviation (RMSD) of the backbone beads (BB) was calculated for each subunit in the P+ module and plotted against simulation time (Figs. S2-S9). After the first four microseconds of MD simulation, all the RMSDs settle around a specific value (see Table S5 in the Supplementary Materials and discussion). The ovine and the human replicas display a similar behavior throughout the simulation time, but the overall RMSD values are slightly higher in the second ones, reasonably because the human models have been derived from the ovine structures. Consequently, all the following analysis were conducted by discarding the first 4 μs of simulation time. The NDs subunits have generally the lowest average RMSD values and, consequently, are suitable for deeper analysis in terms of root mean square fluctuations (RMSF) (Figs. S10-S17). As expected, loops, N- and C-terminals have higher RMSF values than the regions involved in secondary structure elements. The average RMSF were also calculated for each subunit of all systems (see Table S5 and discussion in the Supplementary Materials). Again, the NDs subunits have the lowest average RMSF values. As in the case of the deviations from the starting structures, the ovine replicas show slightly lower average RMSF values than the human replicas. These analyses show that the P+ module and its subunits are stable in the simulations time scale and mutations do not produce easily detectable macroscopic effects on the protein complex. Moreover, the modelled human structures show a similar behavior with respect to the ovine experimental structures, supporting the reliability of the computational setup.

### LHON-associated variants do not affect the P+ module interactions with the membrane

Recent MD simulations (*41, 43*) and high-resolution cryo-EM structures from different organisms (*20, 22, 43–46*) suggest that CI interacts with different lipids of the inner mitochondrial membrane. Among these, cardiolipin (CDL) seems to be involved in the activity and stabilization of CI (*20, 25, 46, 47*). Hence, an analysis of the distribution of the lipids around the protein has been performed and a density map of each lipid was calculated for all the simulated systems (Figs. S18-S25). This analysis confirmed that CDL mostly interacts at the interface between ND2-ND4 and ND4-ND5 and with the ND1 subunit (*20, 25, 41, 43*). Moreover, the maps show a higher density around the distal end of CI, suggesting that CDL may also bind ND5 subunit. Palmitoyl-oleoyl phosphatidylcholine (POPC) and palmitoyl-oleoyl phosphatidylethanolamine (POPE) both form nonspecific or transient interactions with P+ module. Conversely, palmitoyl-oleoyl-phosphatidylinositol (POPI) is mostly distributed around the P+ module and probably interacts in a stronger but nonspecific way with all the subunits. Moreover, the lipid distributions are identical in the *wild type* and in the mutated systems. Consequently, the examined mutations do not cause changes in the interaction between the P+ module and the membrane.

### The p.T536A/ND5 variant does not affect the P+ module dynamics

A detailed analysis of the RMSF per residue calculated on the BB beads was conducted on all the NDx subunits, with also a close-up of the regions around each of the three mutations (Fig. 2 and Figs. S26-S32). Specifically, the average RMSF values and the standard deviation were calculated on the three replicas of each system. Considering the spatial distance between the p.T536A/ND5 mutation and the others, this variant was firstly analyzed. The ND5 RMSF behavior of the human open and close conformations is similar in the wild type and mutated systems (Fig. S29) and the comparison between the ovine ND5 fluctuations and those observed in the human models are superimposable (Fig. S32). Visual inspection and a contact analysis of all the residues found within a cut-off of 0.6 nm from each of the mutated residues show that the interaction network of p.T536A/ND5 with other residues in the ND5 subunit does not change significantly in both *wild type* and mutated systems. Indeed, p.T536A/ND5 interacts with residues that are not involved in CI activity (Fig. S33). Consequently, from the calculations performed according to this approach the p.T536A/ND5 mutation does not affect the P+ module dynamics and was excluded from further analysis.

**Fig. 2.**
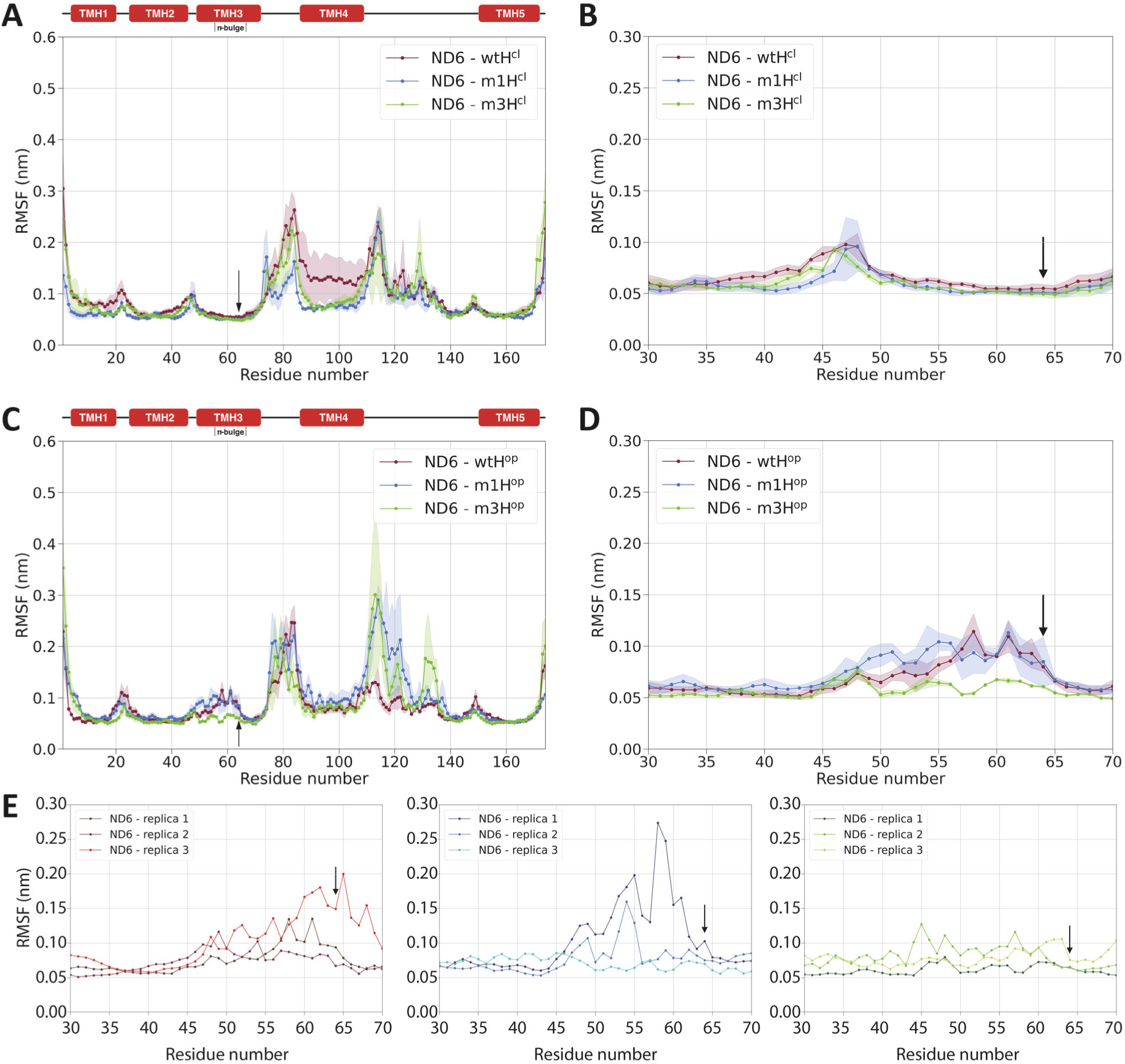
RMSF of the human ND6 subunit in the open and close conformation. (**A** and **C**) Average root mean square fluctuations (RMSF) values as a function of the residue number for ND6 in the human close and open conformation, respectively. The *wild type* ND6 is in red, while the single and the triple mutants are in blue and green, respectively. The width of the shading shows the standard deviation calculated on the three replicas of each simulation. The position of TMHs is reported above the plots as well as the position of the π-bulge in ND6 TMH3. (**B** and **D**) Details of the average RMSF in the p.M64V/ND6 region, which includes the C-terminal part of TMH2, TMH3 and the loop between them, for the human close and open systems, respectively. (**E**) Detail of RMSF as a function of the residue number for each replica in the *wild type* (left panel), single mutant (central panel) and triple mutant (right panel) in the same ND6 region depicted in panels **B** and **D**. The *wild type* RMSF values are in different shades of red, while the single and triple mutants are in different shades of blue and green, respectively. In all panels the mutation position is indicated by a black arrow.

### P+ module fluctuations are affected by the p.M64V/ND6 variant and worsened by the coexistence of the p.A71T/ND4L variant

Considering the p.A71T/ND4L mutation, the RMSF analysis shows that the fluctuations of the region around this mutation in the *wild type* and mutated systems in both conformations are similar (Fig. S28). In the close conformation, the RMSF of p.M64V/ND6 in the *wild type* and mutated systems does not show any effects either (Fig. 2A,B). However, in the open conformation the *wild type* (wtH^op^) and the single mutant (m1H^op^) trajectories show a detectable increase to ca. 0.12 nm of the average fluctuations in the region between residues 45-65 of the ND6 subunit. The latter behavior is present in both the ovine and human *wild type* systems. On the contrary, m3H^op^ behaves similarly to the human systems in close conformation (Fig. 2C,D). Indeed, the average RMSF of the m3H^op^ system is between 0.05 and 0.07 nm with a small standard deviation. Even two replicas of the m1H^op^ system show very small RMSF in the region around the p.M64V/ND6 variant, while the third replica shows significantly larger values (Fig. 2E). In contrast, the RMSF values of the wtH^op^ system range between 0.08 and 0.20 nm for all three replicas. Consequently, it seems that the p.M64V/ND6 mutation alone affects the fluctuations of ND6 TMH3, while the presence of also the p.A71T/ND4L mutation induces a more consistent change in the flexibility of the ND6 π-bulge (Fig. 2E).

### A new proposal for ND6 TMH3 conformational transition during the CI catalytic cycle and the effects of the p.M64V/ND6 and p.A71T/ND4L mutations

Remarkably, visual inspection of the ND6 45-65 region revealed a striking different behavior between the simulations in open and close conformation. In the latter, ND6 TMH3 is tightly folded and does not show significant conformational changes. On the other hand, in the open conformation this region at least partially loses its secondary structure within the first 4 μs of simulation time in all replicas. In the ND6 45-65 region, low RMSF values conceal two different behaviors: *i*) in the close systems the RMSF trend is attributable to small fluctuations with respect to the starting structure; while *ii*) in the open conformation, two out of three replicas in m1H^op^ and all the three m3H^op^ replicas fluctuate less than all the wtH^op^ replicas. Interestingly, one of the m1H^op^ replicas shows an RMSF behavior similar to the wtH^op^ replicas. A possible explanation of this behavior can be found in the way RMSF is calculated. Indeed, RMSF is a measure of the average displacement of each residue compared to the average structure calculated on the last 12 μs of each replica. This means that the replicas with lower RMSF values in the ND6 TMH3 region have smaller fluctuations compared to the average unfolded structure, which in turn is different from the folded starting structure. This unfolding phenomenon has never been reported before for CI, but is not completely unexpected (*48*). According to the catalytic mechanism proposed by Sazanov and co-workers (*20*), CI must undergo a conformational transition from open to close conformations and *vice versa*. This involves a rearrangement of hydrogen bonds that stabilize the secondary structure of the protein in several regions, including also ND6 TMH3. Specifically, it seems that in the open to close transition half of ND6 TMH3 rotates by about 180°. During such a conformational change, several hydrogen bonds are simultaneously broken, while in a helix unfolding event the hydrogen bonds are broken one by one. Consequently, the unfolding of ND6 TMH3 is more energetically favored than the ND6 TMH3 twisting. In other words, the present simulations indicate that the π-bulge can convert into an α-helix, and *vice versa*, through an unfolding/refolding event (*48*). Interestingly, another π-bulge located in ND1 TMH4 remains stable in all the replicas of every system. Moreover, other simulations have shown that π-helix fragments are stable in the time scale explored by our simulations (*40*). For these reasons, the unfolding event could not be ascribed to the CG representation. Finally, visual inspection of the trajectories shows that the unfolding event of the ND6 TMH3 always starts from the loop connecting ND6 TMH3 and TMH4, which is in direct contact with the ubiquinone cavity of CI (*20*) and, in turn, with the Q module in the complete protein. Additionally, an analysis of the water beads in the ND6 TMH3 region (see Supplementary Materials) showed that unfolding is not caused by water insertion between the helices. Consequently, the observed conformational transition is not a direct consequence of solvent exposure of regions that would normally be distant from it.

### Estimation of p.M64V/ND6 and p.A71T/ND4L mutations effect

In order to provide a quantitative description of the effect caused by the primary p.M64V/ND6 mutation alone or in combination with the rare p.A71T/ND4L variant, the origin of the different pattern of fluctuations in the *wild type* protein and the two mutants was explored. In particular, we focused on the π-bulge region by performing a set of analyses including: *i*) the study of the distances between residue pairs considering; *ii*) the backbone dihedral angles in ND6 TMH3; and *iii*) the interaction network formed by each of the mutated residues.

#### Study of the distances between residue pairs

The time-evolving matrix of the distances between residue pairs within 0.7 nm from the point mutations p.M64V/ND6 was calculated for *wild type*, the single and triple mutant systems (see Supplementary Materials). This set is composed of 41 amino acids (“BB set” hereafter, Fig. 3A). An essential dynamics analysis performed on the time-distance matrix (see Supplementary Materials) (*49, 50*) showed that the projections on the first three principal components (PC) suffice to describe the overall dynamics of the region of interest (Fig. S33). The *wild type* system explores a wider range of conformations, while the triple mutant appears to be more rigid in both conformations (Fig. S33). Although it is not possible to distinguish between different conformational states, the fluid dynamics by which each conformation evolves into others within a single macro-state can be analyzed. Specifically, the network elasticity as a function of the lag-time can be estimated (see Supplementary Materials) from the variance, which is roughly inversely proportional to the elasticity of the network. Fig. 3B shows the effect of point mutations on the variance of the network calculated as a function of the lag-time (τ), for open and close systems. In general, the close conformations are stiffer than the open ones. In particular, the m3H^cl^ BB set is stiffer by 11% than wtH^cl^ and m1H^cl^ in a time scale comprised between pico- and nanoseconds.

**Fig. 3.**
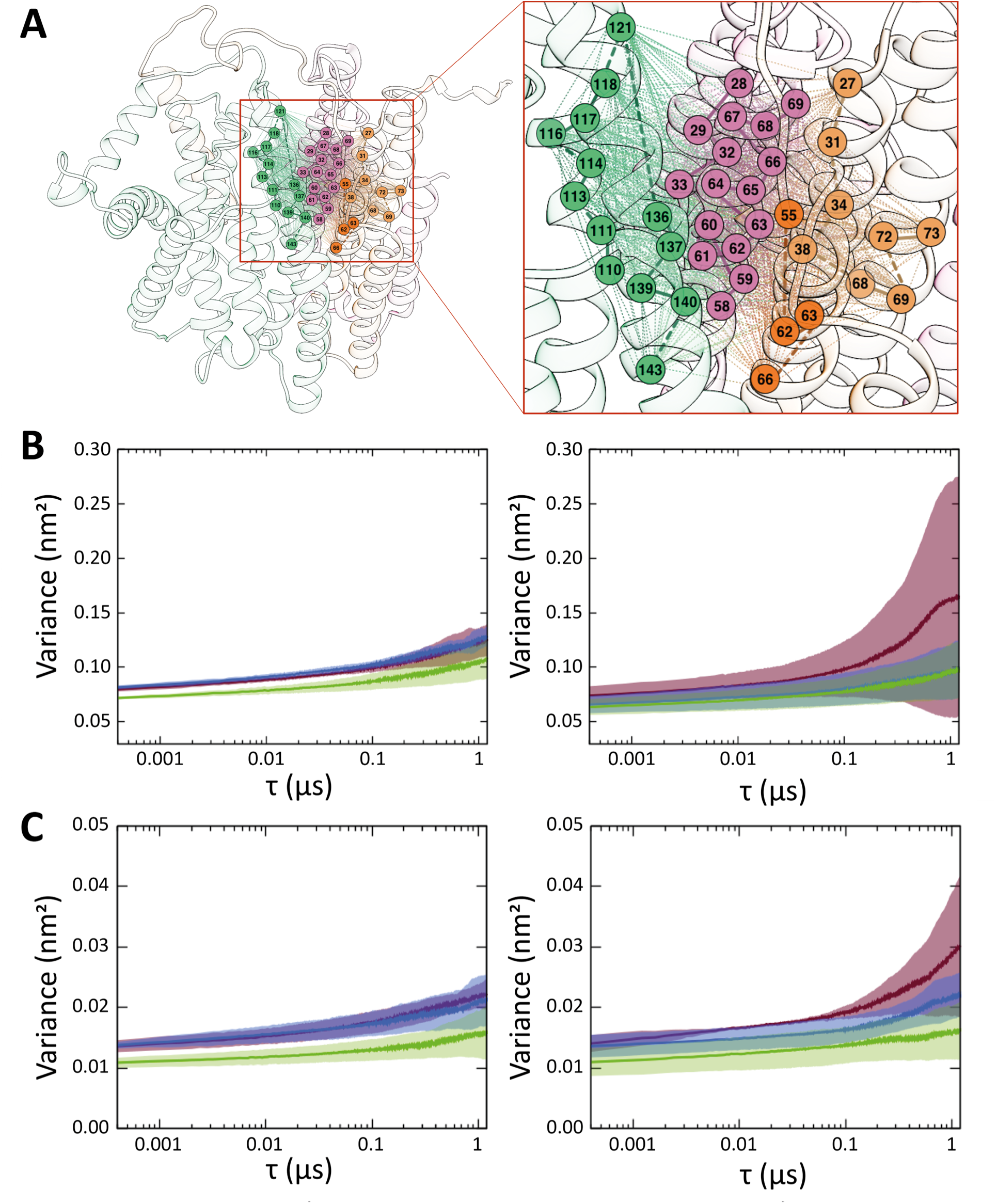
Analysis of the time evolving distance matrices in the p.M64V/ND6 region in the open and close conformation. (**A**) P+ module region within 0.7 nm from p.M64V/ND6. The ND1, ND3, ND4L and ND6 subunits’ transparent ribbons are colored as in Fig. 1. (**B, C**) Variance of the network calculated as a function of the lag-time (τ) in the close (right panel) and open (left) system, for the BB set (**B**) and for the CM set (**C**). In panels **B** and **C**, the *wild type* is in red, while the single and triple mutants are in blue and green. The standard deviation of the variance obtained by averaging over the three replica is represented as a shaded area.

Considering that this effect is observed in the triple mutant only, it can then be ascribed to the presence of the p.A71T/ND4L variant together with p.M64V/ND6. On the other hand, the wtH^op^ system is more flexible and explores more states than m1H^op^ and m3H^op^ in a time scale longer than 0.1 µs. Indeed, m1H^op^ and m3H^op^ BB are similar and both ca. 20% stiffer than the *wild type* system. Therefore, the effect observed on the flexibility of the open conformation is ascribed to the p.M64V/ND6 mutation with only a small additional effect of the p.A71T/ND4L change. To validate the previous result, a similar analysis was conducted using a slightly different set of residues and including the effect of side chains. To do so, the centers of mass (CM) of all the residues within 0.6 nm from p.M64V/ND6 together with the CM of the amino acids likely involved in the proton transfer (*20, 24*) (namely: Y142/ND1, E192/ND1, D66/ND3, E34/ND4L, E70/ND4L and A71T/ND4L) were used to create the “CM set”. The variance calculated as a function of the lag-time of the CM set network (Fig. 3C) agrees with the previous results (Fig. 3B). In the close conformation, m3H^cl^ is 38% stiffer than wtH^cl^, while in the open conformation both m1H^op^ and m3H^op^ are stiffer than wtH^op^ (20% and 38% respectively). In the open conformation m3H^op^ is more rigid than the m1H^op^, which, in turn, is significantly more rigid than the other systems.

#### Analysis of backbone dihedral angles in ND6 TMH3

To understand whether the different elasticity behavior of the ND6 π-bulge region can be ascribed to intra-helical or inter-helical interactions; two further analyses were performed. First, the intra-helical contribution was investigated by calculating the distribution of the dihedral angles formed by the ND6 TMH3 BB beads (residues 56-66), which probably plays a key role in the proton transfer (*20*). In the close conformation, the dihedral angles show a sharp distribution centered around 60 degrees, a typical value of α-helices in the CG representation (Fig. S35). Consequently, the p.M64V/ND6 and p.A71T/ND4L variants cannot induce any evident intra-helical conformational change. On the other hand, the dihedral distributions of the open systems show several conformations that are different from the typical ones of both α- or π-helices (Fig. 4). This is clearly a consequence of the unfolding event observed by visual inspection of the trajectories. The PC analysis conducted on the dihedral angles values (dihedral PCA) (*51*) can provide a measure of the conformational space explored by the π-bulge region in all systems. The total variance of the protein backbone, which is inversely proportional to the stiffness of the system, is reported in Table 1. Since the analysis of the RMSF showed that the individual replicas of the three open systems behaved differently (Fig. 2E), the dihedral PCA was performed on each replica. Two wtH^op^ replicas (namely replicas 1 and 3) as well as one m1H^op^ replica (replica 1) has a larger variance than the other systems. In agreement with the RMSF observations, replica 1 of m1H^op^ is the most flexible. However, the remaining m1H^op^ and m3H^op^ replicas are stiffer than the others, again in agreement with the RMSF observations. Looking at the average values of the cumulative variance in the three systems, p.M64V/ND6 variant induces a partial stiffening of the π-bulge region, which is amplified by the presence of the p.A71T/ND4L change. Considering that the total variance gives information on the ability of the whole π-bulge region to switch from one conformational state to another, it is possible to explain why the three replicas of the m3H^op^ system show small fluctuations compared to an average conformation that remains roughly stable throughout the simulation time (Fig. 2E). To confirm the latter result with a different approach, the variance of each separate conformational state was estimated by fitting with an apt number of Gaussian functions each peak observed in the dihedral angle distributions (Figs. S35-S44). The mean variance has been measured by averaging the half-height widths [σ(φ); i.e., the standard deviation] of all the Gaussian functions used in the fitting of each replica (Table 1). The trend of the average σ(φ) agrees with the results of the dihedral PCA analysis. This correlation suggests that: *i*) all the systems can explore a limited number of specific states and *ii*) a low mean variance of the states is associated with a low total variance and indicates that these systems cannot easily switch from one state to another in the time scale explored.

**Fig. 4.**
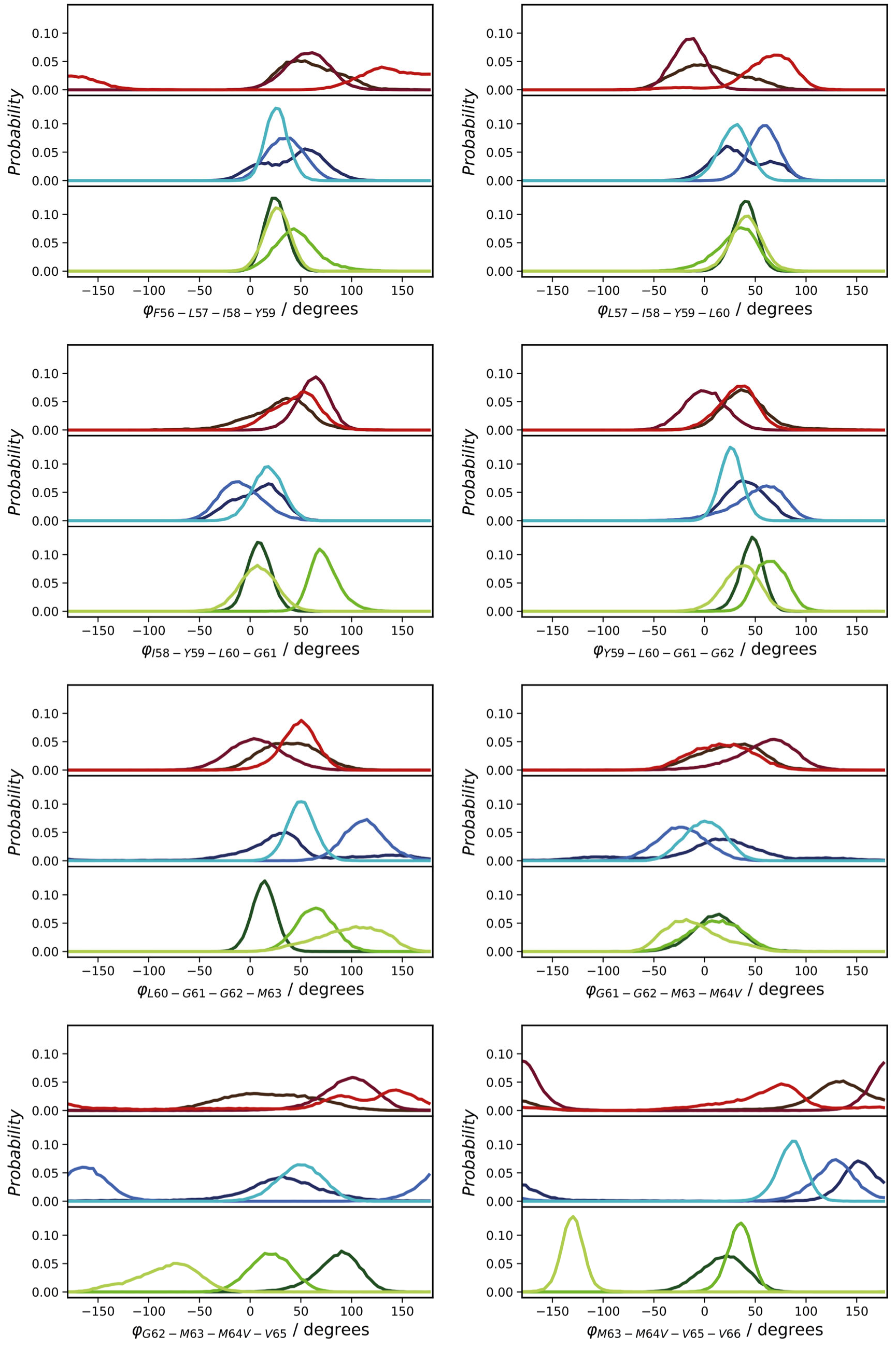
Analysis of the BB dihedrals in the ND6 TMH3 π-bulge region in the open systems. Distribution of the backbone dihedral angles observed throughout the simulations in the open systems. The *wild type* dihedrals are in different shades of red, while the single and triple mutants are in different shades of blue and green, respectively.

**Table 1.** Total variance and average amplitude of ND6 TMH3 BB dihedral angle distribution in the open systems. Total variance calculated as the sum of the eigenvalues derived from the dihedral PCA analysis (*51*) and average half-height widths [σ(φ), degrees] of the Gaussian curves used to fit the dihedral angles distributions in Figs. S28-S36). The last column reports the average values over the three replicas for each system.

| System | Replica 1 |  | Replica 2 |  | Replica 3 |  | Average |  |
| --- | --- | --- | --- | --- | --- | --- | --- | --- |
| | Total variance | Avg. $\sigma(\varphi)$ (degrees) | Total variance | Avg. $\sigma(\varphi)$ (degrees) | Total variance | Avg. $\sigma(\varphi)$ (degrees) | Total variance | Avg. $\sigma(\varphi)$ (degrees) |
| wtH<sup>op</sup> | 2.06 | 49 | 1.16 | 44 | 2.36 | 48 | 1.8 ± 0.6 | 47 ± 3 |
| m1H<sup>op</sup> | 2.59 | 46 | 1.10 | 39 | 0.58 | 31 | 1.4 ± 1.0 | 40 ± 8 |
| m3H<sup>op</sup> | 0.64 | 31 | 0.83 | 34 | 1.07 | 40 | 0.9 ± 0.2 | 35 ± 5 |

#### Interactions formed by each of the mutated residues

The inter-helical interactions were studied by considering all the residues found within a cut-off of 0.6 nm from each of the mutated residues. Figs. 5 and 6 show the fraction of simulation time of each residue found within the cut-off distance from amino acids 64 of ND6 and 71 of ND4L subunit, respectively. In the open conformation, the number of residues within the cut-off distance from amino acid 64 of ND6 subunit is larger than in the close one (Fig. 5). This can be caused by the unfolding of ND6 TMH3. In fact, in the open conformation p.M64V/ND6 moves away from its initial position and comes closer to residues that otherwise would be too far in the folded structure. Therefore, the smaller number of residues within the cut-off distance in the close conformation is probably caused by the stability of the secondary structure of the ND6 TMH3. Interestingly, many of the residues with greater residence time in the open conformation (i.e., ca. >30%, Fig. 5) are found in all three systems studied (wtH^op^, m1H^op^ and m3H^op^). Conversely, the residues with lower residence time (i.e., ca. <30%) are found in only one of the three systems. This means that some of the residues within the cut-off in the wtH^op^ system are not observed in the two mutated systems and some residues are only found for m1H^op^ and m3H^op^. Due to the larger apolar character of valine compared to the *wild type* methionine (see Supplementary Materials) (*35, 40*), it appears that p.M64V/ND6 variant is able to form unspecific interactions with other hydrophobic residues and to settle in meta-stable conformational states upon unfolding. The specific conformation and the contacts established by p.M64V/ND6 in each meta-stable state are peculiar for each replica. The distance analysis on the residues close to amino acid 71 of ND4L subunit shows that the number of residues is roughly maintained in both conformations (Fig. 6), coherently with the conformational stability of ND4L TMH3. Furthermore, the residence times involving this mutated residue are longer in both the triple mutant systems. Indeed, the sum of the residence times in the m3H^op^ and m3H^cl^ systems is significantly higher than in both the *wild type* and single mutant systems. This result is reasonably expected because threonine has a larger steric hinderance than alanine. Moreover, threonine contains a polar group in the side chain, that increases the polar character (see Supplementary Materials) (*35, 40*). Therefore, the mutated residue can form a higher number of interactions with other residues than alanine and, again, this observation can explain the stiffening in the distance network of both BB and CM sets in the triple mutant.

**Fig. 5.**
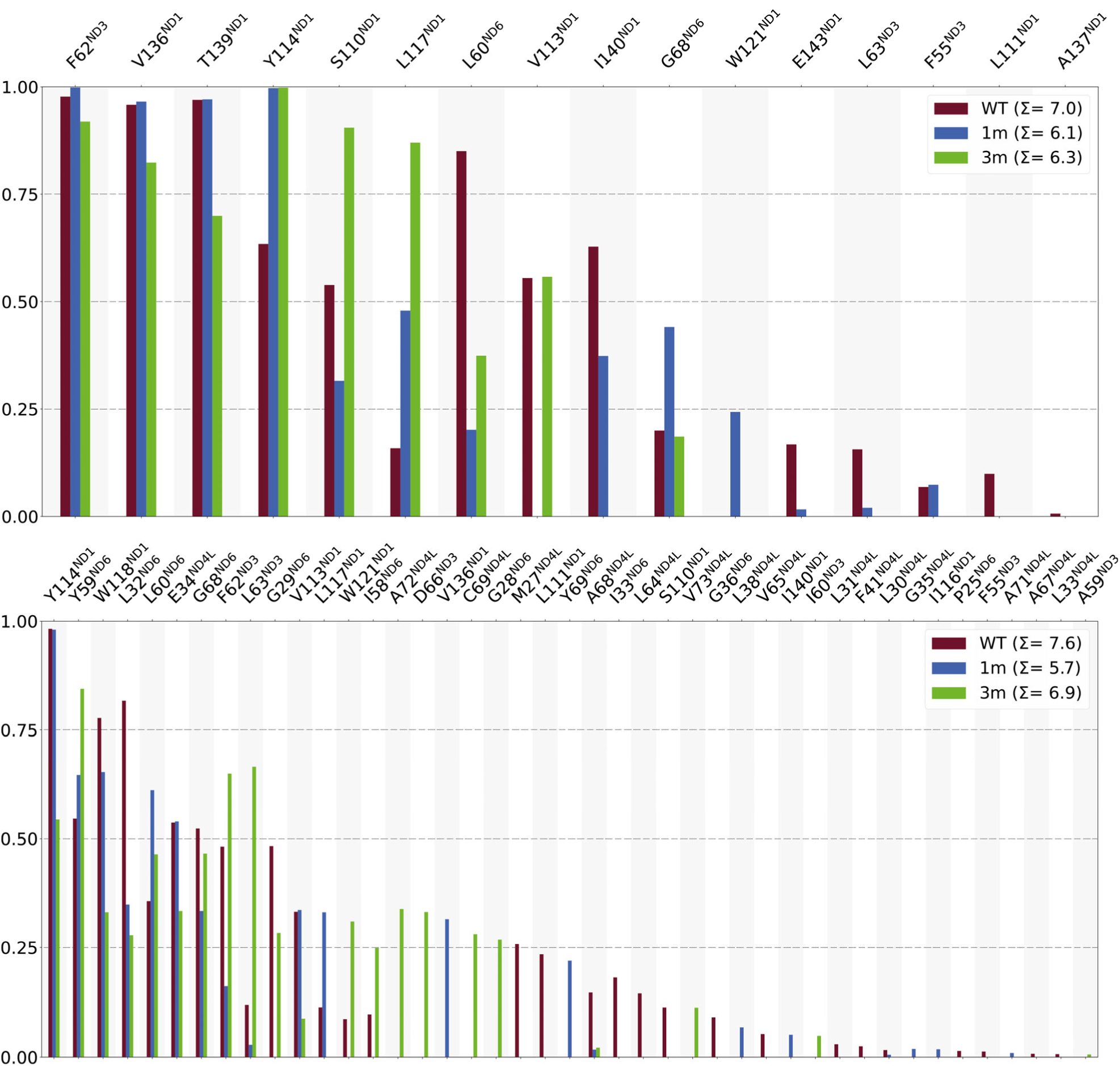
Residues within 0.6 nm of p.M64V/ND6 and residence time within cut-off. The bars are colored in red, blue and green for the *wild type*, single mutant, and triple mutant simulations, respectively. The height of the bars is proportional to the fraction of simulation time. The upper and bottom panels refer to the simulations conducted starting from the close and the open conformation, respectively. The three residues on both sides of p.M64V/ND6 in the ND6 sequence were excluded from the analysis.

**Fig. 6.**
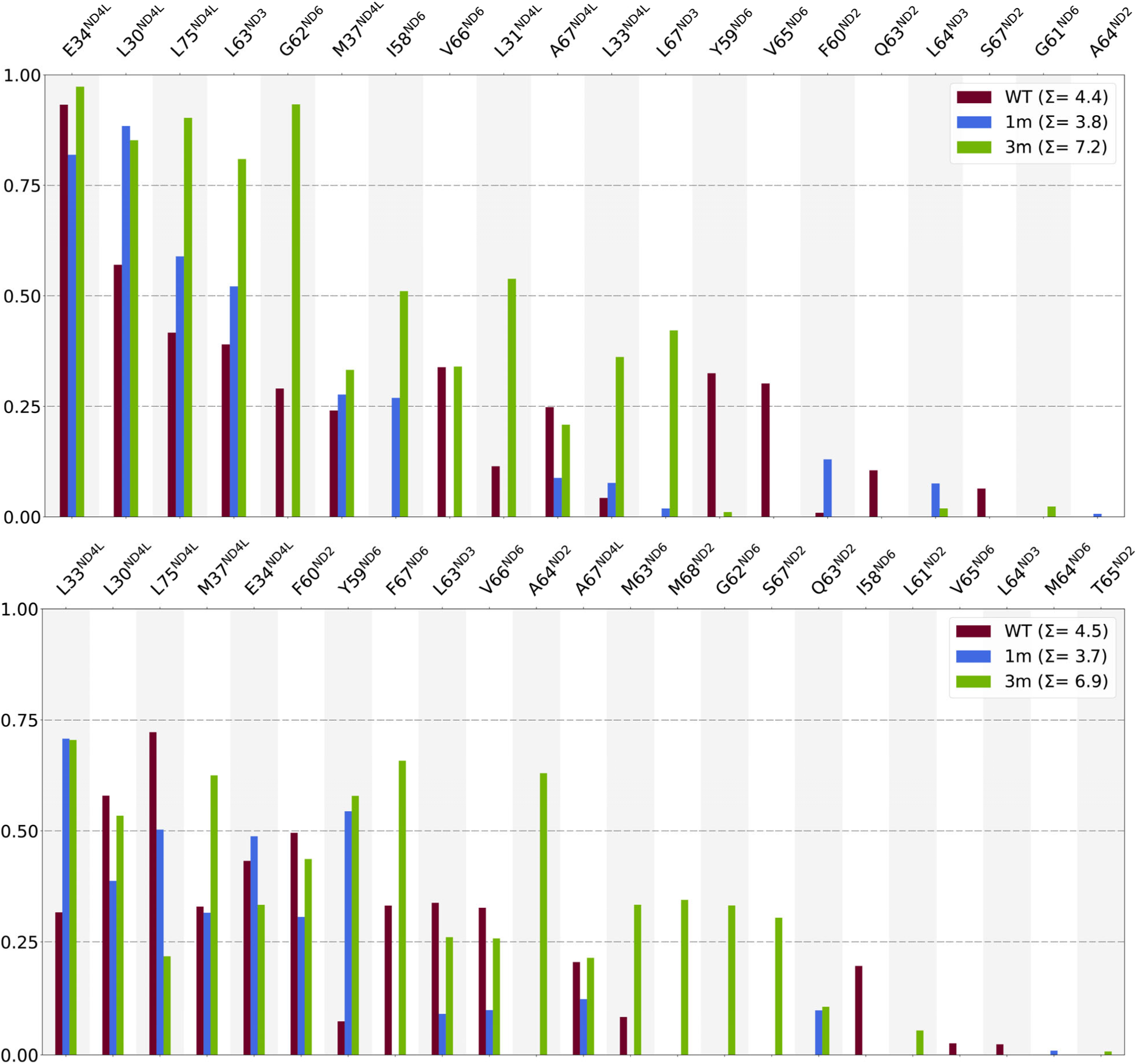
Residues within 0.6 nm of p.A71T/ND4L and residence time within cut-off. The bars are colored in red, blue and green for the *wild type*, single mutant, and triple mutant simulations, respectively. The height of the bars is proportional to the fraction of simulation time. The upper and bottom panels refer to the simulations conducted starting from the close and the open conformation, respectively. The three residues on both sides of p.A71T/ND4L in the ND4L sequence were excluded from the analysis.

## DISCUSSION

This computational study investigated the structural and the dynamical effects of three CI variants associated with LHON, namely m.14484T>C/*MT-ND6* (p.M64V/ND6), m.10680G>A/*MT-ND4L* (p.A71T/ND4L) and 13942A>G/*MT-ND5* (p.T536A/ND5), providing the proof of concept that a CG MD approach can be exploited to predict the impact of putatively pathogenic mtDNA when found alone or in combinations. Indeed, we here show that two of the three mutations, close to each other in the putative E-channel, cooperate in altering the conformational changes, which were previously reported to characterize the open/close states of CI. This phenomenon affects CI efficiency in energy conservation by slowing down proton transport, that is compatible with the currently available biochemical investigations (*9, 10*).

Tens of microsecond-long CG molecular dynamics simulations were carried out starting from the experimental ovine cryo-EM structure as well as on the human models of the CI P module with the addition of the NDUFS2/49 kDa (ovine/human nomenclature). For all the systems, open and close conformations, which correspond to two different stages of the proposed CI catalytic cycle, were considered. The human *wild type* models behave like the experimentally obtained ovine structures, confirming the reliability of the modelling procedure. The analysis of the simulations suggested that the p.T536A/ND5 variant does not affect the proton transfer and the protein function. This change was never observed in LHON patients without the concomitant presence of other pathogenic variants. The p.T536A/ND5 mutation affects the ND5 subunit, which is in contact with subunit ND4. Visual inspection of the trajectories and both the RMSD and RMSF analyses on the dynamics of these two subunits show that there are no significant changes in their secondary structure between closed and open system, as observed in previous experimental studies (*20, 23, 27*). Then, the focus was placed on the remaining two variants, p.M64V/ND6 and p.A71T/ND4L, which are close to each other in the putative E-channel. This proton channel, mainly constituted by glutamate residues, connects the quinone site to the three antiporter-like subunits (*20, 24*) and is composed of ND1, ND3, ND6 and ND4L subunits. According to one of the CI catalytic cycle current hypotheses (*20, 24*), the conformational transitions occurring in the E-channel region are crucial for CI function. The activity of the enzyme is probably associated also with a large conformational transition involving the Q and N modules together with other movements inside the P module (Fig. 1). Specifically, the open to close conformational transition comprises the rearrangement of the ND6 TMH3 and the conversion of a π-bulge fragment into an α-helix. Indeed, the p.M64V/ND6 and p.A71T/ND4L changes are at the hinge between the P module and the remaining CI modules, specifically the former on ND6 TMH3 and the latter on ND4L TMH3. Additionally, to understand the possible cooperative effect of the p.A71T/ND4L mutation on the puzzling pathogenic p.M64V/ND6 mutation, a first set of simulations was conducted on a system carrying this mutation alone. Then a second set of calculations comprising also the p.A71T/ND4L change was considered. The structural and multivariate analyses carried out on such simulations suggested that:

*i)* the two mutations induce a stiffening of the ND6 TMH3 π-bulge region with different effects on the open and close conformations. In particular, p.M64V/ND6 affects the open conformation, while the additional presence of p.A71T/ND4L has a detectable effect on both conformations.
*ii)* in the open conformation, the ND6 TMH3 unfolds in all the simulated systems, but the presence of one or both changes causes the persistence of specific meta-stable conformational states that are not present, on average, in the *wild type* system.

Hence, we conclude that these two phenomena may slow down and/or partially prevent the conformational transition from open to closed and *vice versa*, affecting the coupling between ubiquinone reduction and proton pumping in CI. These results may explain the puzzling pathogenic or non-pathogenic effect of m.14484T>C/*MT-ND6* (p.M64V/ND6) mutation in different mtDNA backgrounds, giving some clues about the role of rare variants in mitochondrially encoded subunits of CI. The combination of these rare aminoacidic changes may have and additive effect on the slight defect induced by m.14484T>C/*MT-ND6* (p.M64V/ND6), significantly decreasing OXPHOS efficiency with the consequent overall bioenergetic defect behind the disease.

### Perspectives

The computational setup used here can be used for the systematic characterization and classification of CI mutations and population-specific non-synonymous variants, alone or in combinations, at a limited computational cost. While the functional effects of single variants can be relatively easily evaluated through biochemical investigations in cell models such as fibroblasts and cybrids, the outcomes of combinations of putatively pathogenic variants are more difficult to define, due to the lack of proper models. In this frame, this computational approach may overcome such experimental limitations, permitting an evaluation of all the possible combinations of mtDNA missense variants in CI genes. Moreover, these findings could be supported by the determination of the high-resolution experimental structures of both *wild type* and mutated human CI. This would allow experimental validation of the observed changes in the network of interactions involving the ND6 TMH3 region. Also, further use of cybrid cells can be instrumental to study and functionally validate the effect of the mutations studied here as well as other and new mutations or population-specific non-synonymous variants.

### Limitations

By grouping atoms together, a CG model reduces the degrees of freedom and, consequently, the computational cost of the simulations and their analysis (*52*). Moreover, the dynamics of a CG system are enhanced through the elimination of additional friction and noise that are present in an atomistic representation. These features make CG representation ideal for dealing with a protein complex of the size of the CI or even just its P+ module. The reduction in the number of degrees of freedom causes a change in the inter-residue interaction scheme as this occurs between beads and no longer between atomistic functional groups.

The absence of the N and Q modules might introduce some artifacts and structural reorganization or accelerate/make impossible certain events that take place during the catalytic cycle. For example, the use of only the P+ module has probably made faster the unfolding of the ND6 TMH3 π-bulge in the open conformation and may have prevented its refolding. On the other hand, the time scale of the open/close transition – and *vice versa* – is likely to be greater than the one considered in the simulations reported here. However, the aim of the present investigation was to study the overall structural and dynamical effects of three specific mutations and not to draw conclusions on mechanistic aspects, for which much longer time scales and atomistic details are required.

Finally, when the simulations reported here were already in progress, a new version of Martini force field (Martini 3) (*53*) was released. The new force field includes improved interaction parameters, additional bead types and the inclusion of additional interactions such as hydrogen bonds and electronic polarizability. It would possibly be interesting to repeat the simulations reported here using Martini 3.

## MATERIALS AND METHODS

### Reconstruction of unsolved regions in ovine P+ module structures

The solved cryo-EM structures of the CI P+ module from *O. aries* in both open (PDB id: 6ZKA) and close conformation (PDB id: 6ZKB) were utilized as main template for the wtO^op^ and wtO^cl^ systems, respectively. The unsolved regions in the selected structures were reconstructed by using Modeller 10.0 (*54*) and generating 100 models for each system. The ovine CI structures in both open (PDB id: 6ZKE) and close conformation (PDB id: 6ZKC) were chosen as template proteins for the missing regions. Table S2 offers a list of the modelled regions in both systems. For each system, the best model was selected on the basis of the DOPE potential (*55*) included in Modeller and was subsequently analyzed using ProCheck (*56*). A loop optimization procedure comprising the generation of 500 models for each system was then applied using Modeller to correct structurally problematic regions. The best models were selected and analyzed as above. Unlike the other P+ module subunits, the 49 kDa subunit is largely modelled and is not in its physiological environment. Consequently, the behavior of this subunit is not considered in this study.

### Homology modelling of human P+ module structures

The *wild type* human structural models in both open and close conformation were generated by using the optimized ovine structures as template. Specifically, the target human aminoacidic sequences (Table S3) were modelled with Modeller 10.0 by using the same procedure described above. The human and ovine sequences have high sequence identity (Table S3) ranging from 60% to 95%, with an average of about 80%. The selected variants (p.M64V/ND6, p.A71T/ND4L and p.T536A/ND5) were created by using the *swapaa* tool included in UCSF Chimera 1.16 (*57*).

### Phosphopantetheine CG mapping and parametrization

The acyl-carrier supernumerary subunits located in the Q and P modules covalently bind a molecule of phosphopantetheine (S-[2-({N-[(2S)-2-hydroxy-3,3-dimethyl-4-(phosphonooxy)butanoyl]-beta-alanyl}amino)ethyl] tetradecanethioate, ZMP) each, which in turn stabilizes through its acyl chains the two supernumerary subunits NDUFAB1. ZMP prosthetic moiety is covalently linked via phosphodiester bond to Ser44 of the acyl carrier protein subunit in the P module. The ZMP molecule was parameterized according to Martini 2.2 procedure based on similarities with phosphatidylserine. First, the atomistic representation of ZMP was mapped into an 8-bead coarse-grained model to split the molecule into reasonable existing building blocks (see Fig. S45). Then, the Martini bead types were assigned based on the chemical building block they are taken to represent. See Supplementary Materials for further details.

### Systems setup

The coarse-grained representation of the two ovine and the six P+ modules was built by using the *Martini Maker* tool (*58, 59*) included in the CHARMM-GUI web server (*60*). Each protein system was converted to the CG representation by using the Martini 2.2 force field (*35, 36*) and then was embedded in a 300 x 300 Å bilayer membrane. The lipid composition was chosen to mimic the human inner mitochondrial membrane, with 40% POPC, 30% POPE, 16% POPI and 14% CDL (*61*). The protein-membrane systems were, then, enclosed in a rectangular box with water and 0.15 M of Na^+^ and Cl^-^ ions, plus counterions to ensure electroneutrality. The use of an elastic network (Elnedyn) was not implemented here to let the system evolve from the initial structure into different states. In CG MD simulations the elastic network prevents deformations of the protein and maintains the tertiary structure and its drift. However, the main goal of this investigation was to track and measure such conformational changes and detect the regions in which they mainly occur. The obtained CG systems have approximately 144,000 beads each. A phosphopantetheine and an AMP molecule (*62*) was added *a posteriori* to each system removing overlapping solvent beads.

### System minimization, equilibration, and production runs

Each system underwent two energy minimization and five restrained equilibration stages using GROMACS 2020.1 (*63–65*). The two energy minimization stages consisted of 5,000 steps of steepest descent without position restraints. The number of steps, the time steps and the force constants of the positional restraints used in the equilibration stages are resumed in Table S7. In all the equilibration stages the leap-frog algorithm was used. In the energy minimization and equilibration stages, periodic boundaries were applied to the systems and the electrostatic interactions were calculated through the Particles Mesh Ewald method (*66*). The cut-off values for the real part of the electrostatic interactions and for the van der Waals interactions were set to 1.1 nm. The temperature and the pressure were regulated with the velocity-rescale thermostat (*67*) and the Berendsen barostat (*68*).

Production runs were performed by using the leap-frog algorithm with a time step of 20 fs and a temperature of 303.15 K. The temperature and the pressure of the systems were set by the velocity-rescale thermostat and the Parrinello-Rahman barostat (*69*). CG models yield a speed-up of 2-3 orders of magnitude relative to all-atom MD (*70*), which opens to the exploration of big size scales to microsecond time lengths. This is probably caused by the larger particle sizes, which generates a smoother energy landscape. The effective time sampled with MARTINI models is averagely 4-fold larger than the all-atoms models. Consequently, MARTINI simulated times are usually multiplied by the standard conversion factor of 4 (*35*) and all the analysis reported here have been already scaled. *Trajectory analysis.* The simulated trajectories were analyzed with GROMACS tools and the Python packages MDAnalysis (*71*), MDTraj 1.9.8 (*72*) and mdciao 0.0.5 (*73*), using in-house Jupyter notebooks (*74*). In the RMSD analyses, the initial cryo-EM structures in the CG representation were considered as reference. Lipids density maps (see Fig. S18-S25) were carried out using the GROMACS *densmap* tool, and the plots were generated using *Gnuplot* (http://www.gnuplot.info). The upper and lower membrane leaflets, named Matrix and IMS, respectively, were defined as the volume slices (along the axis perpendicular to the membrane) between the center of mass of the membrane and the POPC, CDL2, POPI and POPE heads. Other plots were generated with matplotlib library (*75*). UCSF Chimera 1.16 (*57*) and UCSF ChimeraX (*76*) were used for the molecular figures. The dihedral angles distributions were analyzed by using the dihedral PCA procedure (*51*).

## Acknowledgments

The authors thank Giorgio Turtù for technical help and insightful discussions.

## Author contributions

Conceptualization: LR, FL, LC, LI, AMG, FM

Methodology: LR, FL, AB, JF, FM

Investigation: LR, FL, AB, JF, FM

Visualization: LR, FL, AB, JF, FM

Supervision: FL, FM

Writing—original draft: LR, FL, LC, AB, JF, FZ, LI, VC, AMG, FM

Writing—review & editing: LR, FL, LC, AB, JF, FZ, LI, VC, AMG, FM

## Data and materials availability

All data needed to evaluate the conclusions of the paper are present in the paper and/or the Supplementary Materials. The trajectories are available upon request.

