## Supplementary Information for "A computational study to assess the pathogenicity of single or combinations of missense variants on respiratory Complex I"

Supplementary Materials for

### Computational study of mutations associated with Leber's hereditary optic neuropathy in the transmembrane domain of Complex I

Laura Rigobello, Francesca Lugli, Leonardo Caporali, Alessio Bartocci, Jacopo Fadanni, Francesco Zerbetto, Luisa Iommarini, Valerio Carelli, Anna Maria Ghelli, Francesco Musiani

**This PDF file includes:**

Supplementary Text
Figs. S1 to S45
Tables S1 to S7

**Table S1. List of human and ovine CI subunits.**

| *H. sapiens* | | *O. aries* | |
|---|---|---|---|
| **Subunit** | **Abbreviation** | **Subunit** | **Abbreviation** |
| NADH dehydrogenase [ubiquinone] flavoprotein 1, mitochondrial | NDUFV1 | NADH dehydrogenase [ubiquinone] flavoprotein 1, mitochondrial | 51 kDa |
| NADH dehydrogenase [ubiquinone] flavoprotein 2, mitochondrial | NDUFV2 | Mitochondrial complex I, 24 kDa subunit | 24 kDa |
| NADH-ubiquinone oxidoreductase 75 kDa subunit | NDUFS1 | NADH:ubiquinone oxidoreductase core subunit S1 | 75 kDa |
| NADH dehydrogenase [ubiquinone] iron-sulfur protein 2, mitochondrial | NDUFS2 | Mitochondrial complex I, 49 kDa subunit | 49 kDa |
| NADH dehydrogenase [ubiquinone] iron-sulfur protein 3, mitochondrial | NDUFS3 | NADH:ubiquinone oxidoreductase core subunit S3 | 30 kDa |
| NADH dehydrogenase [ubiquinone] iron-sulfur protein 7, mitochondrial | NDUFS7 | Mitochondrial complex I, PSST subunit | PSST |
| NADH dehydrogenase [ubiquinone] iron-sulfur protein 8, mitochondrial | NDUFS8 | Mitochondrial complex I, TYKY subunit | TYKY |
| NADH dehydrogenase [ubiquinone] flavoprotein 3, mitochondrial | NDUFV3 | Mitochondrial complex I, 10 kDa subunit | 10 kDa |
| NADH dehydrogenase [ubiquinone] iron-sulfur protein 6, mitochondrial | NDUFS6 | Mitochondrial complex I, 13 kDa subunit | 13 kDa |
| NADH dehydrogenase [ubiquinone] iron-sulfur protein 4, mitochondrial | NDUFS4 | NADH dehydrogenase [ubiquinone] iron-sulfur protein 4, mitochondrial | NDUFS4 |
| NADH dehydrogenase [ubiquinone] 1 alpha subcomplex subunit 9, mitochondrial | NDUFA9 | NADH:ubiquinone oxidoreductase subunit A9 | NDUFA9 |
| NADH dehydrogenase [ubiquinone] 1 alpha subcomplex subunit 2 | NDUFA2 | NADH dehydrogenase [ubiquinone] 1 alpha subcomplex subunit 2 | NDUFA2 |
| NADH dehydrogenase [ubiquinone] 1 alpha subcomplex subunit 5 | NDUFA5 | Mitochondrial complex I, B13 subunit | NDUFA5 |
| NADH dehydrogenase [ubiquinone] 1 alpha subcomplex subunit 6 | NDUFA6 | NADH:ubiquinone oxidoreductase subunit A6 | NDUFA6 |
| NADH dehydrogenase [ubiquinone] 1 alpha subcomplex subunit 7 | NDUFA7 | Mitochondrial complex I, B14.5a subunit | NDUFA7 |
| NADH dehydrogenase [ubiquinone] 1 alpha subcomplex subunit 12 | NDUFA12 | NADH dehydrogenase [ubiquinone] 1 alpha subcomplex subunit 12 | NDUFA12 |
| NADH-ubiquinone oxidoreductase chain 3 | ND3 | NADH-ubiquinone oxidoreductase chain 3 | ND3 |
| NADH-ubiquinone oxidoreductase chain 1 | ND1 | NADH-ubiquinone oxidoreductase chain 1 | ND1 |
| NADH-ubiquinone oxidoreductase chain 6 | ND6 | NADH-ubiquinone oxidoreductase chain 6 | ND6 |
| NADH-ubiquinone oxidoreductase chain 4L | ND4L | NADH-ubiquinone oxidoreductase chain 4L | ND4L |
| NADH-ubiquinone oxidoreductase chain 5 | ND5 | NADH-ubiquinone oxidoreductase chain 5 | ND5 |
| NADH-ubiquinone oxidoreductase chain 4 | ND4 | NADH-ubiquinone oxidoreductase chain 4 | ND4 |
| NADH-ubiquinone oxidoreductase chain 2 | ND2 | NADH-ubiquinone oxidoreductase chain 2 | ND2 |
| NADH dehydrogenase [ubiquinone] 1 alpha subcomplex subunit 11 | NDUFA11 | Mitochondrial complex I, B14.7 subunit | NDUFA11 |
| NADH dehydrogenase [ubiquinone] 1 beta subcomplex subunit 5, mitochondrial | NDUFB5 | NADH:ubiquinone oxidoreductase subunit B5 | NDUFB5 |
| Acyl carrier protein, mitochondrial | NDUFAB1 | Acyl carrier protein | NDUFAB1 |
| NADH dehydrogenase [ubiquinone] 1 alpha subcomplex subunit 8 | NDUFA8 | NADH dehydrogenase [ubiquinone] 1 alpha subcomplex subunit 8 | NDUFA8 |
| NADH dehydrogenase [ubiquinone] 1 beta subcomplex subunit 10 | NDUFB10 | Mitochondrial complex I, PDSW subunit | PDSW |
| NADH dehydrogenase [ubiquinone] 1 alpha subcomplex subunit 10, mitochondrial | NDUFA10 | NADH dehydrogenase [ubiquinone] 1 alpha subcomplex subunit 10, mitochondrial | NDUFA10 |
| NADH dehydrogenase [ubiquinone] iron-sulfur protein 5 | NDUFS5 | NADH:ubiquinone oxidoreductase subunit S5 | NDUFS5 |

| | | | |
|---|---|---|---|
| NADH dehydrogenase [ubiquinone] 1 alpha subcomplex subunit 3 | NDUFA3 | NADH:ubiquinone oxidoreductase subunit A3 | NDUFA3 |
| NADH dehydrogenase [ubiquinone] 1 beta subcomplex subunit 3 | NDUFB3 | NADH:ubiquinone oxidoreductase subunit B3 | NDUFB3 |
| NADH dehydrogenase [ubiquinone] 1 subunit C2 | NDUFC2 | NADH dehydrogenase [ubiquinone] 1 subunit C2 | NDUFC2 |
| NADH dehydrogenase [ubiquinone] 1 beta subcomplex subunit 4 | NDUFB4 | NADH:ubiquinone oxidoreductase subunit B4 | NDUFB4 |
| NADH dehydrogenase [ubiquinone] 1 alpha subcomplex subunit 13 | NDUFA13 | Mitochondrial complex I, B16.6 subunit | NDUFA13 |
| NADH dehydrogenase [ubiquinone] 1 beta subcomplex subunit 6 | NDUFB6 | Mitochondrial complex I, B17 subunit | NDUFB6 |
| NADH dehydrogenase [ubiquinone] 1 beta subcomplex subunit 7 | NDUFB7 | NADH:ubiquinone oxidoreductase subunit B7 | NDUFB7 |
| NADH dehydrogenase [ubiquinone] 1 beta subcomplex subunit 9 | NDUFB9 | NADH:ubiquinone oxidoreductase subunit B9 | NDUFB9 |
| NADH dehydrogenase [ubiquinone] 1 beta subcomplex subunit 2, mitochondrial | NDUFB2 | NADH:ubiquinone oxidoreductase subunit B2 | NDUFB2 |
| NADH dehydrogenase [ubiquinone] 1 beta subcomplex subunit 8, mitochondrial | NDUFB8 | NADH dehydrogenase [ubiquinone] 1 beta subcomplex subunit 8, mitochondrial | NDUFB8 |
| NADH dehydrogenase [ubiquinone] 1 beta subcomplex subunit 11, mitochondrial | NDUFB11 | Mitochondrial complex I, ESSS subunit | ESSS |
| NADH dehydrogenase [ubiquinone] 1 subunit C1, mitochondrial | NDUFC1 | Mitochondrial complex I, KFYI subunit | KFYI |
| NADH dehydrogenase [ubiquinone] 1 beta subcomplex subunit 1 | NDUFB1 | Mitochondrial complex I, MNLL subunit | MNLL |
| NADH dehydrogenase [ubiquinone] 1 alpha subcomplex subunit 1 | NDUFA1 | Mitochondrial complex I, MWFE subunit | MWFE |

**Table S2. Modelled residues in the ovine P+ module structures.**

| Ovine P+ module subunit | Open | Close |
|---|---|---|
| 49 kDa | 41 - 430 | 41 - 424 |
| ND3 | 31 - 48 | - |
| ND1 | 205 - 210 | - |
| ND6 | 78 - 86 | - |
| NDUFB5 | 1 - 2 | 1 - 2 |
| NDUFAB1 | 1 | 1 |
| PDSW | 1 - 2<br>174 - 175 | 1 - 2<br>174 - 175 |
| NDUFA3 | 1 - 2 | 1 - 2 |
| NDUFB3 | 1 - 10<br>90 - 97 | 1 - 10<br>90 - 97 |
| NDUFA13 | 1 - 4 | 1 - 17 |
| B17 subunit | 38 - 64 | 38 - 64 |
| NDUFB7 | 123 - 136 | 123 - 136 |
| NDUFB9 | 178 | 178 |
| NDUFB2 | 1 - 4<br>70 - 72 | 1 - 4<br>70 - 72 |
| NDUFB8 | 1 - 2 | 1 - 2 |
| ESSS | 1 - 22<br>124 - 125 | 1 - 22<br>124 - 125 |
| MNLL | 1 - 6 | 1 - 6 |

**Table S3. UniProt entries of the human aminoacidic sequences in the modelled systems and sequence identity with respect to the corresponding ovine CI subunit.**

| man subunit | UniProt entry | Sequence identity (%) |
|---|---|---|
| NADH dehydrogenase [ubiquinone] iron-sulfur protein 2 (or 49 kDa subunit) | O75306 | 95.35 |
| NADH-ubiquinone oxidoreductase chain 3 (or ND3) | P03897 | 73.04 |
| NADH-ubiquinone oxidoreductase chain 1 (or ND1) | P03886 | 77.99 |
| NADH-ubiquinone oxidoreductase chain 6 (or ND6) | P03923 | 60.92 |
| NADH-ubiquinone oxidoreductase chain 4L (or ND4L) | P03901 | 76.53 |
| NADH-ubiquinone oxidoreductase chain 5 (or ND5) | P03915 | 71.60 |
| NADH-ubiquinone oxidoreductase chain 4 (or ND4) | P03905 | 75.66 |
| NADH-ubiquinone oxidoreductase chain 2 (or ND2) | P03891 | 63.85 |
| NADH dehydrogenase [ubiquinone] 1 α subcomplex subunit 11 | Q86Y39 | 70.80 |
| NADH dehydrogenase [ubiquinone] 1 β subcomplex subunit 5, mitochondrial | O43674 | 82.52 |
| Acyl carrier protein, mitochondrial | O14561 | 97.73 |
| NADH dehydrogenase [ubiquinone] 1 α subcomplex subunit 8 | P51970 | 87.65 |
| NADH dehydrogenase [ubiquinone] 1 β subcomplex subunit 10 | O96000 | 78.86 |
| NADH dehydrogenase [ubiquinone] 1 α subcomplex subunit 10, mitochondrial | O95299 | 79.69 |
| NADH dehydrogenase [ubiquinone] iron-sulfur protein 5 | O43920 | 70.48 |
| NADH dehydrogenase [ubiquinone] 1 α subcomplex subunit 3 | O95167 | 83.13 |
| NADH dehydrogenase [ubiquinone] 1 β subcomplex subunit 3 | O43676 | 83.51 |
| NADH dehydrogenase [ubiquinone] 1 subunit C2 | O95298 | 75.63 |
| NADH dehydrogenase [ubiquinone] 1 β subcomplex subunit 4 | O95168 | 75.00 |
| NADH dehydrogenase [ubiquinone] 1 α subcomplex subunit 13 | Q9P0J0 | 83.22 |
| NADH dehydrogenase [ubiquinone] 1 β subcomplex subunit 6 | O95139 | 77.95 |
| NADH dehydrogenase [ubiquinone] 1 β subcomplex subunit 7 | P17568 | 85.29 |
| NADH dehydrogenase [ubiquinone] 1 β subcomplex subunit 9 | Q9Y6M9 | 89.33 |
| NADH dehydrogenase [ubiquinone] 1 β subcomplex subunit 2, mitochondrial | O95178 | 90.28 |
| NADH dehydrogenase [ubiquinone] 1 β subcomplex subunit 8, mitochondrial | O95169 | 86.08 |
| NADH dehydrogenase [ubiquinone] 1 β subcomplex subunit 11, mitochondrial | Q9NX14 | 86.29 |
| NADH dehydrogenase [ubiquinone] 1 subunit C1, mitochondrial | O43677 | 79.59 |
| NADH dehydrogenase [ubiquinone] 1 β subcomplex subunit 1 | O75438 | 77.19 |
| NADH dehydrogenase [ubiquinone] 1 α subcomplex subunit 1 | O15239 | 80.00 |

**Fig. S1. Modelled regions in the P+ modules.**

Ribbon diagram of the structures of the ovine P+ module in the open (top panel) and close (bottom panel) conformation with the addition of the modelled regions corresponding to the disordered regions not solved in the structures. The experimentally solved regions are in white, while the modelled ones are in red.

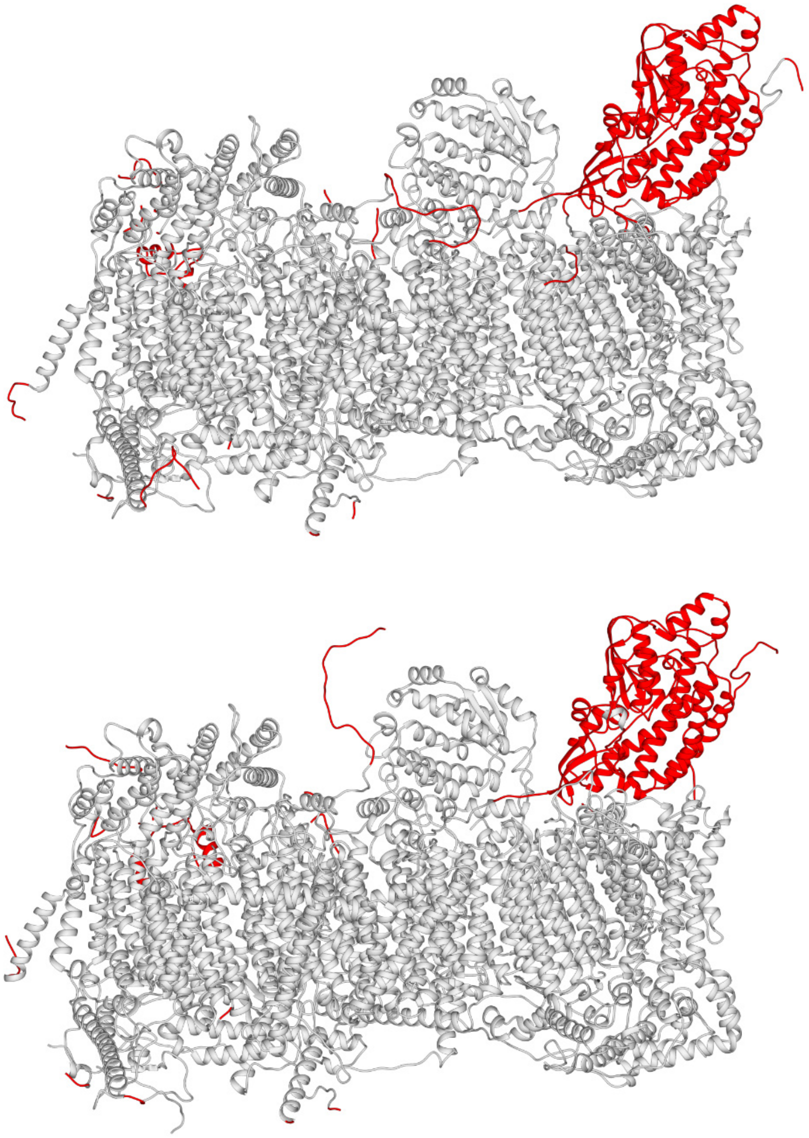

**Fig. S2. Backbone beads RMSD *vs.* simulation time plots of the P+ module subunits in the wtO$^{op}$ structures.**
Each panel label corresponds to the equivalent subunit in Table S1. The RMSD values corresponding to the three replicas are in brown, maroon, and crimson, respectively.

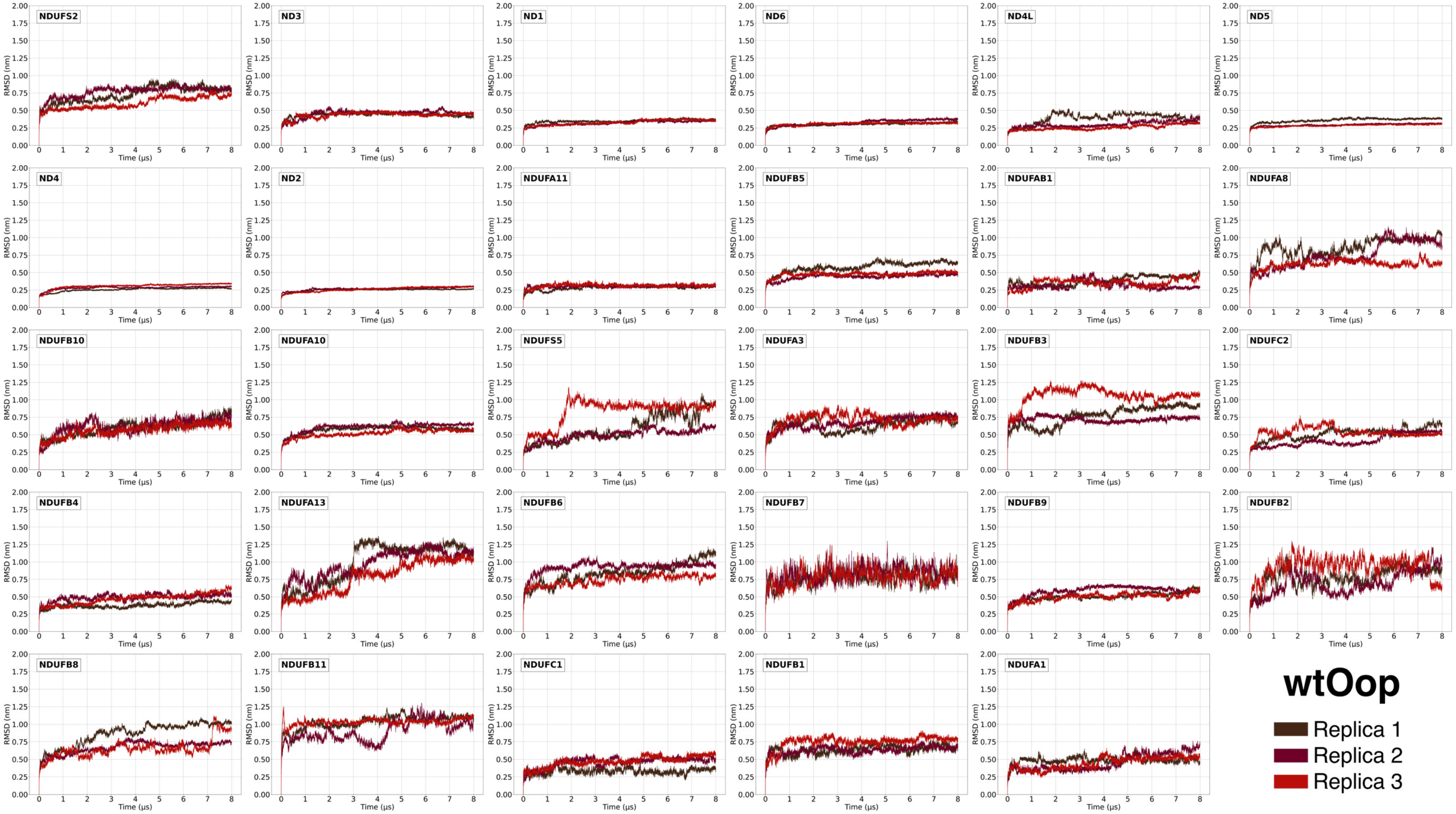

**Fig. S3. Backbone beads RMSD *vs.* simulation time plots of the P+ module subunits in the wtO$^{cl}$ structures.**
Each panel label corresponds to the equivalent subunit in Table S1. The RMSD values corresponding to the three replicas are in brown, maroon, and crimson, respectively.

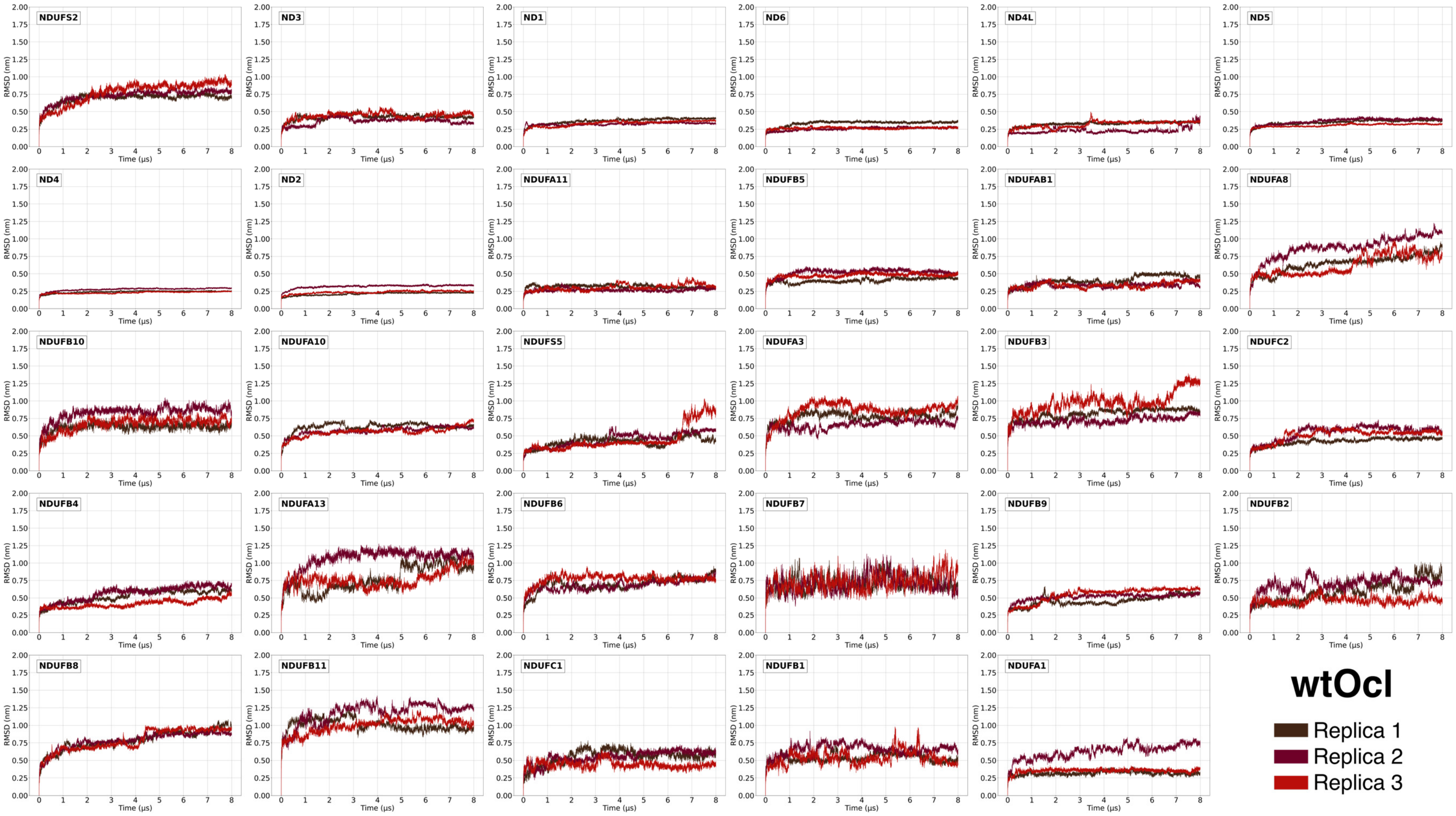

**Fig. S4. Backbone beads RMSD *vs.* simulation time plots of the P+ module subunits in the wtH$^{op}$ structures.** Each panel label corresponds to the equivalent subunit in Table S1. The RMSD values corresponding to the three replicas are in brown, maroon, and crimson, respectively.

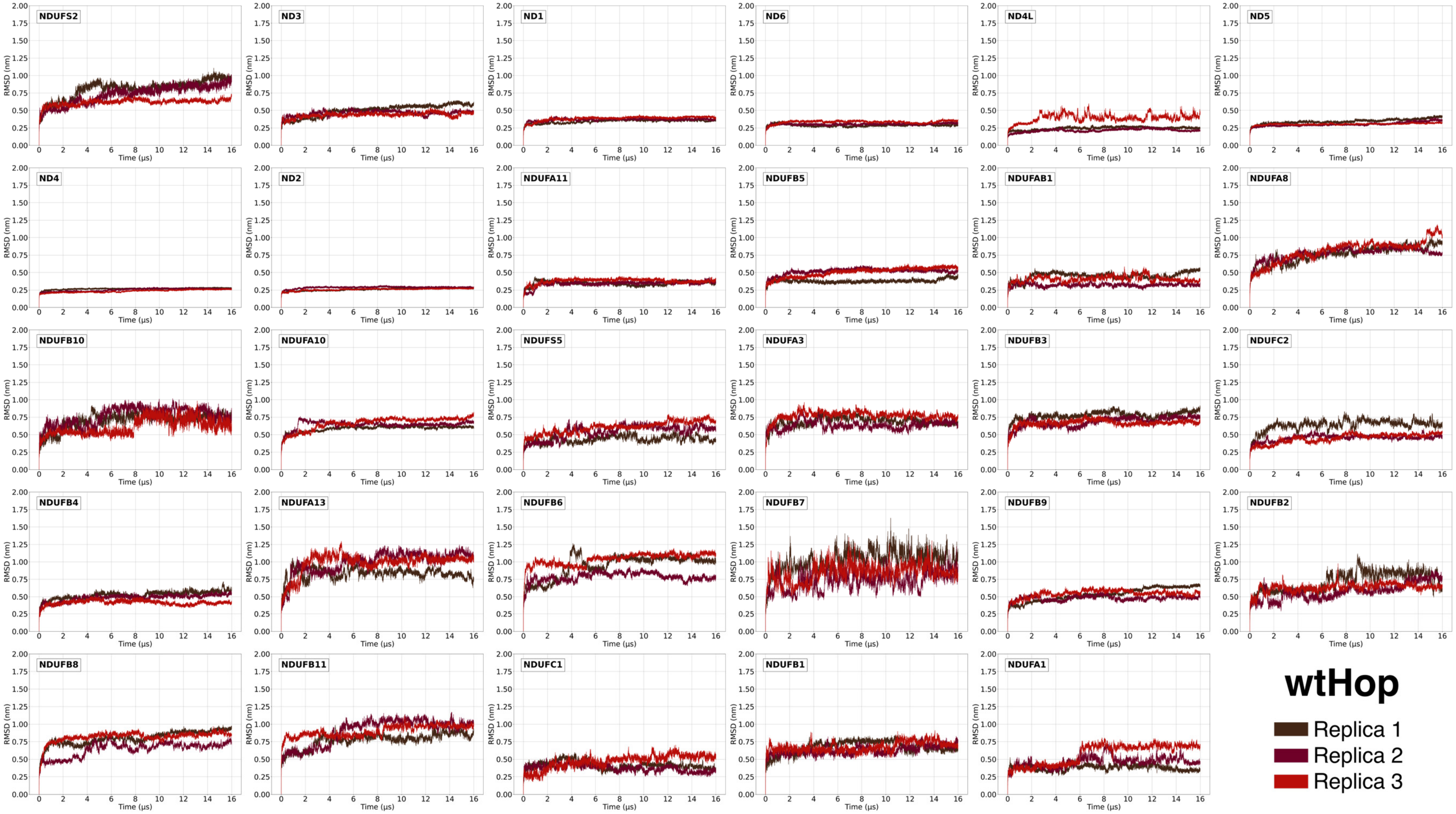

**Fig. S5. Backbone beads RMSD *vs.* simulation time plots of the P+ module subunits in the wtH$^{cl}$ structures.**
Each panel label corresponds to the equivalent subunit in Table S1. The RMSD values corresponding to the three replicas are in brown, maroon, and crimson, respectively.

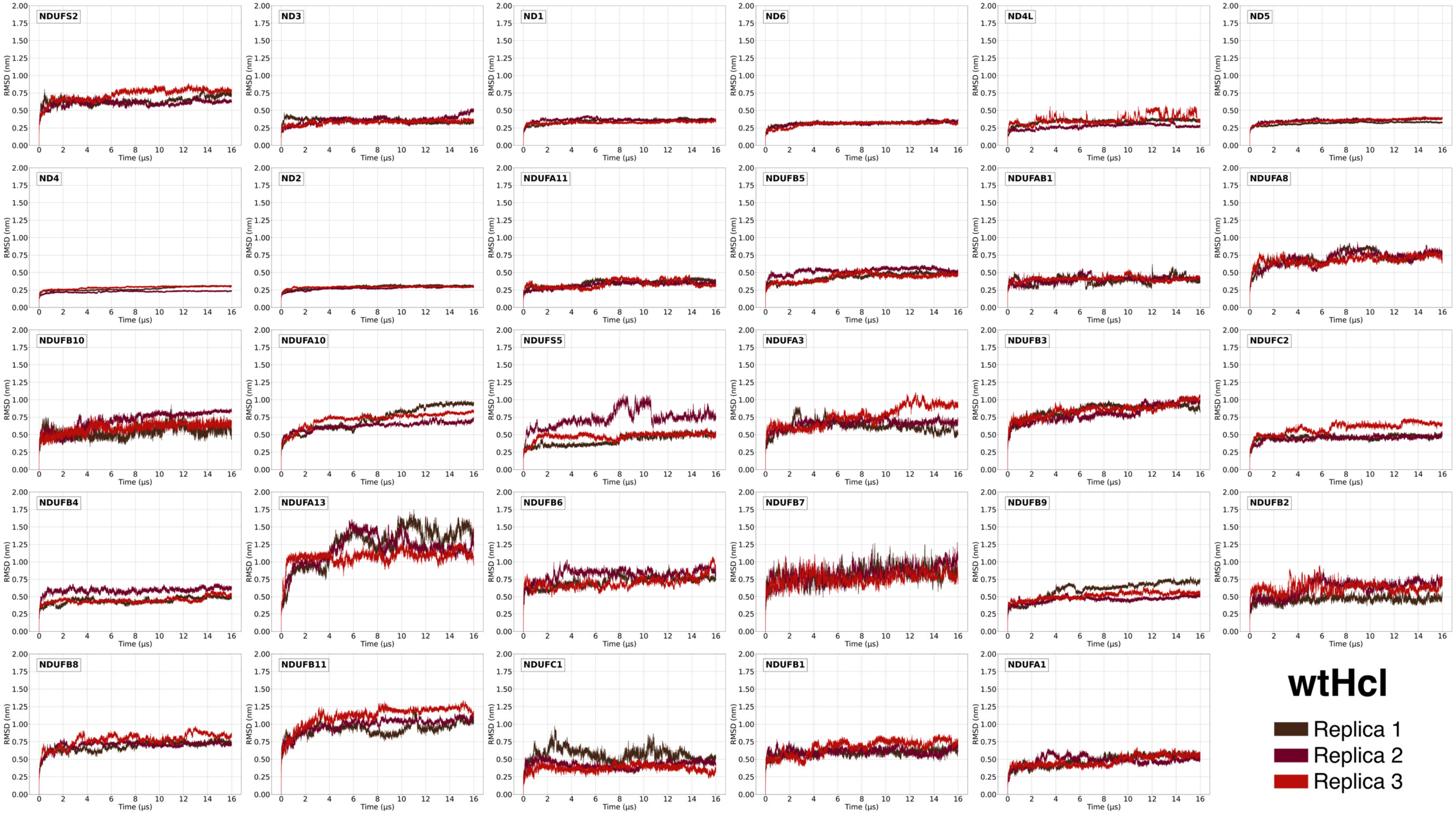

**Fig. S6. Backbone beads RMSD *vs.* simulation time plots of the P+ module subunits in the m1H$^{op}$ structures.** Each panel label corresponds to the equivalent subunit in Table S1. The RMSD values corresponding to the three replicas are in navy, royal blue, and fountain blue, respectively.

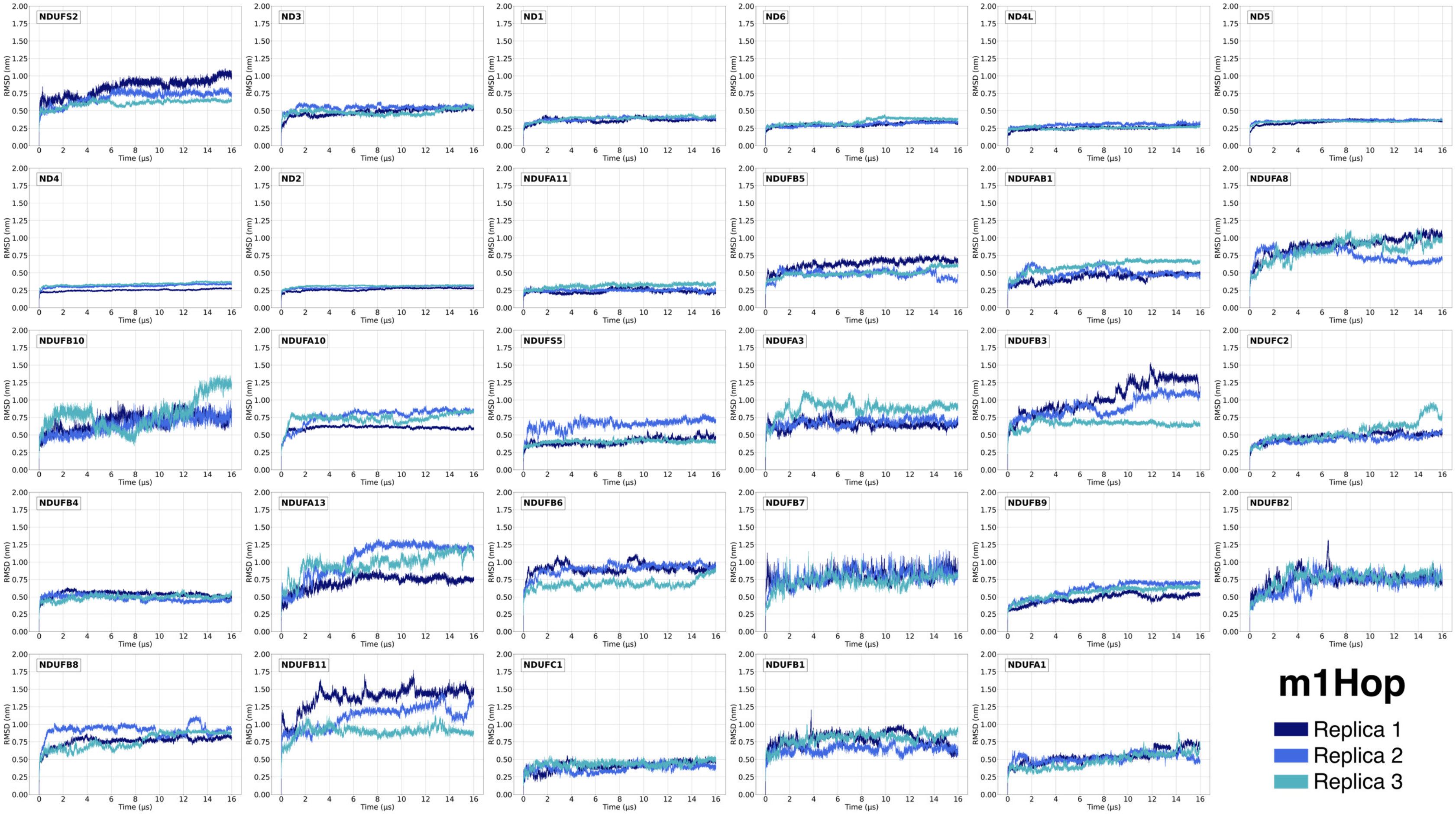

**Fig. S7. Backbone beads RMSD *vs.* simulation time plots of the P+ module subunits in the m1H$^{cl}$ structures.**
Each panel label corresponds to the equivalent subunit in Table S1. The RMSD values corresponding to the three replicas are in navy, royal blue, and fountain blue, respectively.

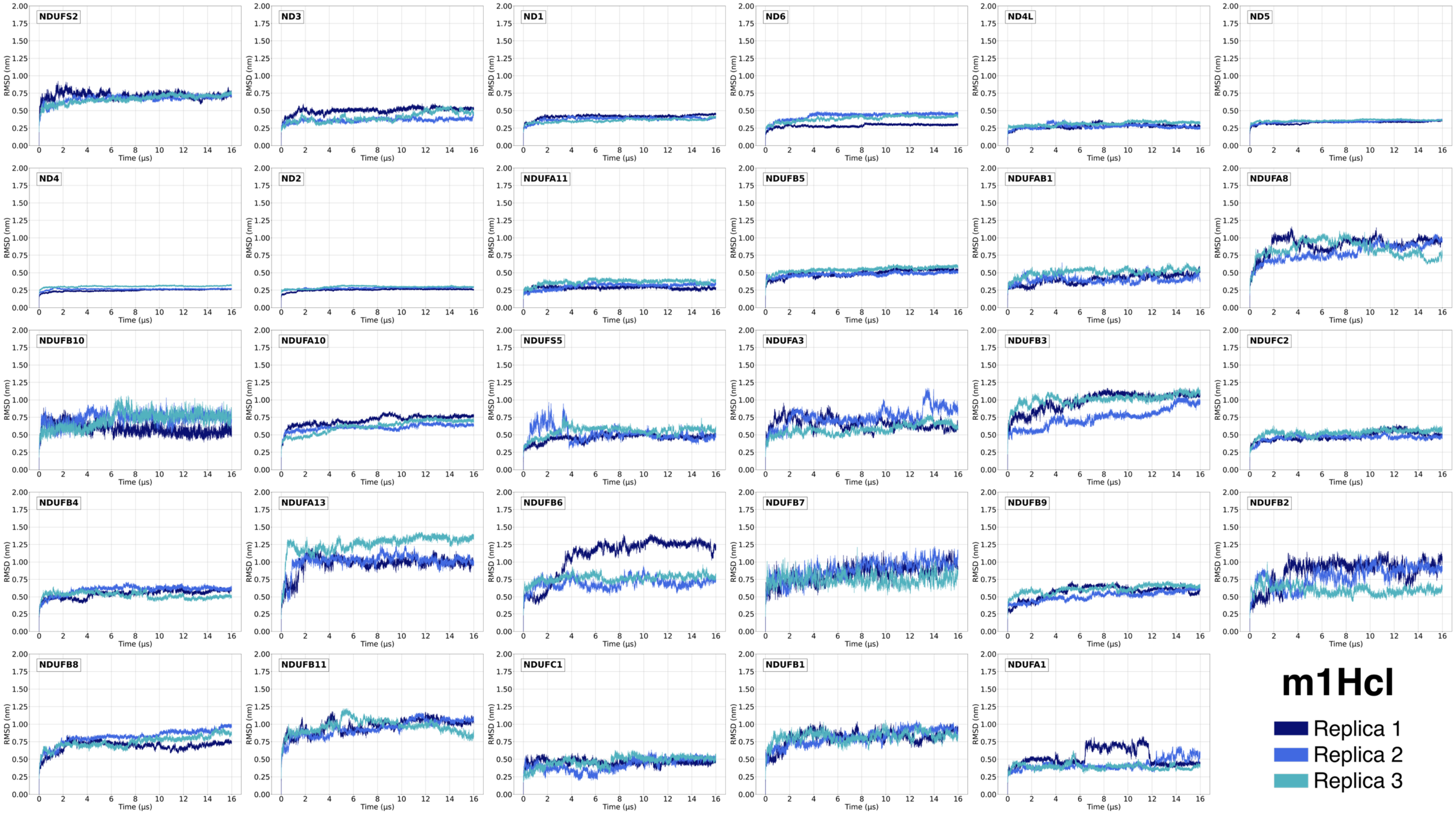

**Fig. S8. Backbone beads RMSD *vs.* simulation time plots of the P+ module subunits in the m3H$^{op}$ structures.**
Each panel label corresponds to the equivalent subunit in Table S1. The RMSD values corresponding to the three replicas are in pine, dark lime, and lime, respectively.

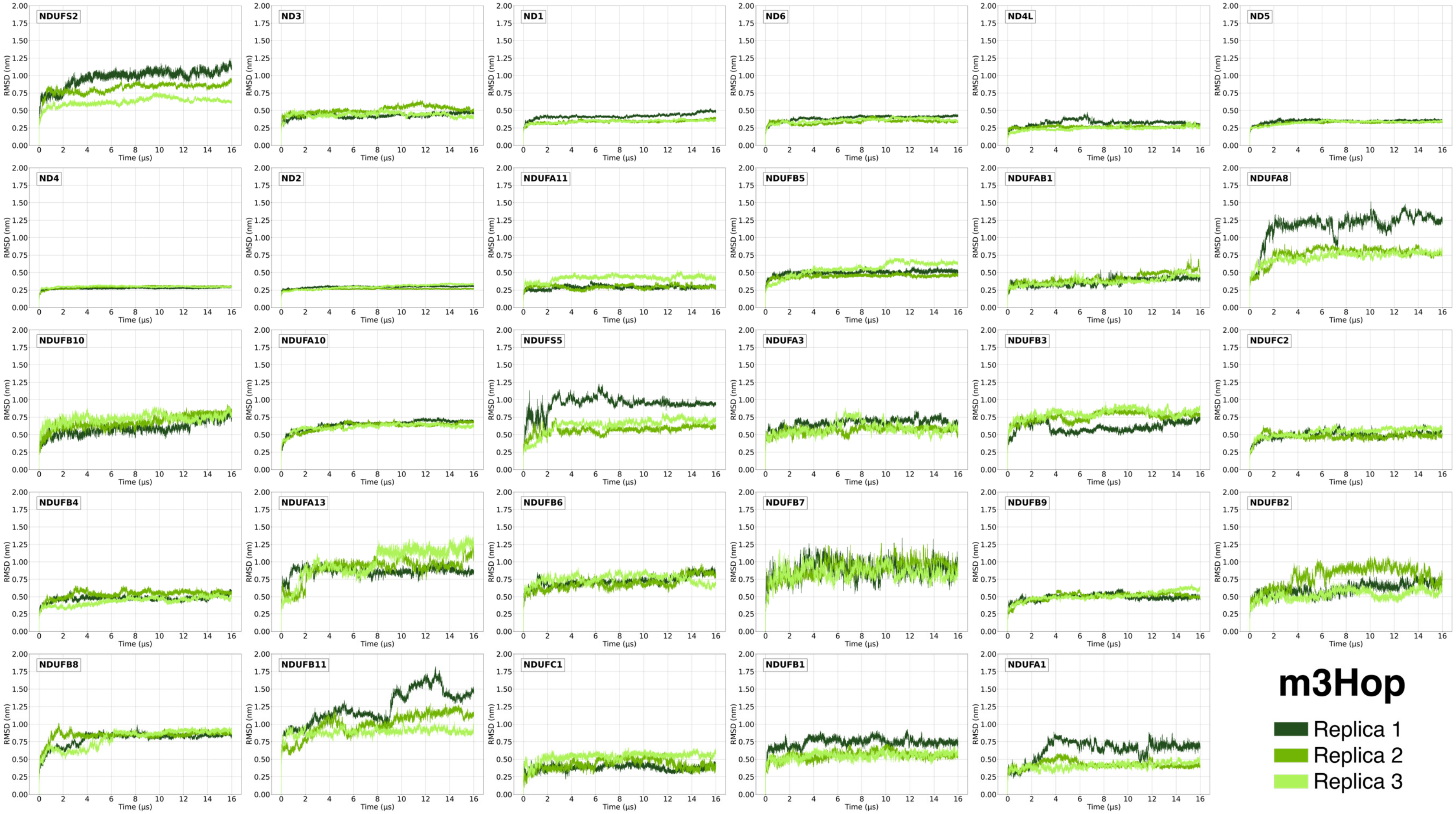

**Fig. S9. Backbone beads RMSD *vs.* simulation time plots of the P+ module subunits in the m3H$^{cl}$ structures.**
Each panel label corresponds to the equivalent subunit in Table S1. The RMSD values corresponding to the three replicas are in dark pine, dark lime, and lime, respectively.

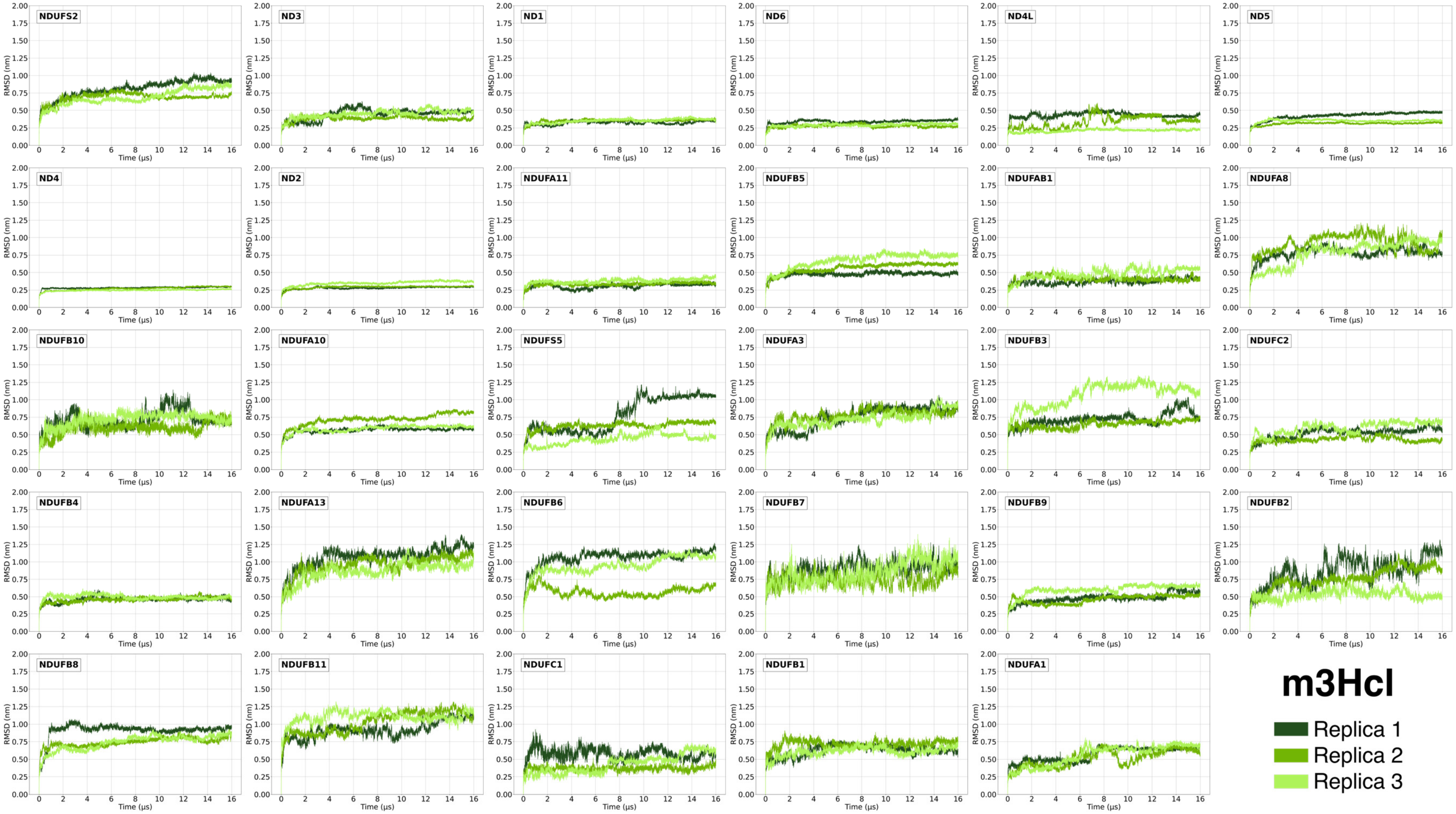

**Table S4. Average and standard deviation of RMSD values (in nm) of backbone beads for each subunit of ovine and human P+ module.** For each subunit, the ovine and human abbreviation (see Table S1) is reported. Each subunits line corresponds to a different replica.

| Human/Ovine subunit | wtO$^{op}$ | wtO$^{cl}$ | wtH$^{op}$ | wtH$^{cl}$ | m1H$^{op}$ | m1H$^{cl}$ | m3H$^{op}$ | m3H$^{cl}$ |
|---|---|---|---|---|---|---|---|---|
| 49 kDa / NDUFS2 | 0.7 ± 0.1 | 0.70 ± 0.08 | 0.8 ± 0.1 | 0.63 ± 0.06 | 0.8 ± 0.1 | 0.71 ± 0.05 | 1.0 ± 0.1 | 0.8 ± 0.1 |
| | 0.77 ± 0.08 | 0.71 ± 0.08 | 0.7 ± 0.1 | 0.60 ± 0.04 | 0.69 ± 0.09 | 0.67 ± 0.06 | 0.82 ± 0.08 | 0.69 ± 0.06 |
| | 0.62 ± 0.08 | 0.8 ± 0.2 | 0.63 ± 0.04 | 0.75 ± 0.09 | 0.61 ± 0.05 | 0.66 ± 0.06 | 0.63 ± 0.06 | 0.7 ± 0.1 |
| ND3 | 0.44 ± 0.03 | 0.42 ± 0.04 | 0.50 ± 0.08 | 0.35 ± 0.03 | 0.48 ± 0.05 | 0.50 ± 0.04 | 0.43 ± 0.03 | 0.45 ± 0.08 |
| | 0.45 ± 0.04 | 0.42 ± 0.06 | 0.45 ± 0.03 | 0.37 ± 0.05 | 0.54 ± 0.03 | 0.36 ± 0.03 | 0.47 ± 0.05 | 0.35 ± 0.02 |
| | 0.43 ± 0.04 | 0.47 ± 0.05 | 0.44 ± 0.03 | 0.32 ± 0.03 | 0.48 ± 0.04 | 0.40 ± 0.07 | 0.44 ± 0.03 | 0.47 ± 0.06 |
| ND1 | 0.35 ± 0.02 | 0.37 ± 0.04 | 0.34 ± 0.02 | 0.34 ± 0.03 | 0.36 ± 0.03 | 0.41 ± 0.03 | 0.42 ± 0.04 | 0.32 ± 0.03 |
| | 0.33 ± 0.03 | 0.33 ± 0.02 | 0.38 ± 0.02 | 0.38 ± 0.02 | 0.38 ± 0.03 | 0.38 ± 0.03 | 0.36 ± 0.02 | 0.37 ± 0.03 |
| | 0.34 ± 0.03 | 0.37 ± 0.03 | 0.40 ± 0.03 | 0.38 ± 0.02 | 0.39 ± 0.03 | 0.36 ± 0.02 | 0.36 ± 0.02 | 0.37 ± 0.02 |
| ND6 | 0.30 ± 0.02 | 0.33 ± 0.04 | 0.29 ± 0.02 | 0.32 ± 0.02 | 0.30 ± 0.03 | 0.28 ± 0.02 | 0.39 ± 0.03 | 0.34 ± 0.02 |
| | 0.32 ± 0.03 | 0.28 ± 0.02 | 0.29 ± 0.02 | 0.32 ± 0.03 | 0.31 ± 0.03 | 0.42 ± 0.05 | 0.34 ± 0.02 | 0.29 ± 0.02 |
| | 0.32 ± 0.02 | 0.28 ± 0.01 | 0.32 ± 0.02 | 0.35 ± 0.03 | 0.34 ± 0.04 | 0.38 ± 0.04 | 0.36 ± 0.03 | 0.32 ± 0.03 |
| ND4L | 0.39 ± 0.07 | 0.33 ± 0.03 | 0.24 ± 0.02 | 0.34 ± 0.03 | 0.25 ± 0.03 | 0.27 ± 0.03 | 0.32 ± 0.04 | 0.43 ± 0.04 |
| | 0.32 ± 0.04 | 0.23 ± 0.04 | 0.24 ± 0.02 | 0.29 ± 0.03 | 0.30 ± 0.02 | 0.27 ± 0.03 | 0.25 ± 0.02 | 0.37 ± 0.07 |
| | 0.26 ± 0.03 | 0.33 ± 0.04 | 0.40 ± 0.06 | 0.37 ± 0.06 | 0.25 ± 0.01 | 0.31 ± 0.03 | 0.26 ± 0.02 | 0.25 ± 0.02 |
| ND5 | 0.36 ± 0.03 | 0.35 ± 0.04 | 0.34 ± 0.03 | 0.31 ± 0.02 | 0.34 ± 0.03 | 0.33 ± 0.02 | 0.34 ± 0.02 | 0.42 ± 0.06 |
| | 0.31 ± 0.01 | 0.40 ± 0.04 | 0.31 ± 0.03 | 0.37 ± 0.02 | 0.36 ± 0.01 | 0.34 ± 0.02 | 0.34 ± 0.02 | 0.33 ± 0.02 |
| | 0.31 ± 0.02 | 0.36 ± 0.02 | 0.32 ± 0.01 | 0.37 ± 0.03 | 0.35 ± 0.01 | 0.36 ± 0.01 | 0.35 ± 0.02 | 0.37 ± 0.02 |
| ND4 | 0.26 ± 0.03 | 0.24 ± 0.02 | 0.27 ± 0.01 | 0.27 ± 0.03 | 0.25 ± 0.02 | 0.24 ± 0.02 | 0.27 ± 0.01 | 0.28 ± 0.01 |
| | 0.30 ± 0.01 | 0.27 ± 0.02 | 0.26 ± 0.02 | 0.24 ± 0.01 | 0.31 ± 0.02 | 0.26 ± 0.01 | 0.31 ± 0.02 | 0.28 ± 0.02 |
| | 0.33 ± 0.03 | 0.26 ± 0.01 | 0.25 ± 0.01 | 0.29 ± 0.02 | 0.34 ± 0.02 | 0.30 ± 0.01 | 0.31 ± 0.01 | 0.29 ± 0.01 |
| ND2 | 0.25 ± 0.02 | 0.21 ± 0.02 | 0.26 ± 0.02 | 0.29 ± 0.02 | 0.26 ± 0.02 | 0.25 ± 0.02 | 0.29 ± 0.01 | 0.28 ± 0.02 |
| | 0.30 ± 0.02 | 0.32 ± 0.02 | 0.31 ± 0.01 | 0.30 ± 0.02 | 0.29 ± 0.02 | 0.28 ± 0.01 | 0.30 ± 0.01 | 0.31 ± 0.01 |
| | 0.29 ± 0.03 | 0.28 ± 0.01 | 0.30 ± 0.01 | 0.31 ± 0.01 | 0.31 ± 0.01 | 0.29 ± 0.02 | 0.29 ± 0.03 | 0.38 ± 0.03 |
| NDUFA11 | 0.28 ± 0.03 | 0.32 ± 0.02 | 0.34 ± 0.03 | 0.35 ± 0.04 | 0.24 ± 0.03 | 0.28 ± 0.02 | 0.29 ± 0.02 | 0.31 ± 0.03 |
| | 0.28 ± 0.02 | 0.27 ± 0.02 | 0.33 ± 0.04 | 0.33 ± 0.04 | 0.25 ± 0.02 | 0.31 ± 0.04 | 0.31 ± 0.03 | 0.35 ± 0.03 |
| | 0.29 ± 0.02 | 0.33 ± 0.03 | 0.40 ± 0.04 | 0.32 ± 0.06 | 0.31 ± 0.03 | 0.36 ± 0.03 | 0.40 ± 0.04 | 0.35 ± 0.03 |
| NDUFB5 | 0.58 ± 0.07 | 0.40 ± 0.03 | 0.38 ± 0.03 | 0.43 ± 0.06 | 0.63 ± 0.07 | 0.49 ± 0.05 | 0.50 ± 0.03 | 0.48 ± 0.03 |
| | 0.44 ± 0.03 | 0.48 ± 0.04 | 0.52 ± 0.04 | 0.55 ± 0.04 | 0.48 ± 0.05 | 0.48 ± 0.03 | 0.44 ± 0.03 | 0.58 ± 0.07 |
| | 0.47 ± 0.03 | 0.46 ± 0.03 | 0.56 ± 0.07 | 0.49 ± 0.06 | 0.50 ± 0.05 | 0.54 ± 0.04 | 0.56 ± 0.09 | 0.6 ± 0.1 |
| NDUFAB1 | 0.38 ± 0.06 | 0.40 ± 0.07 | 0.45 ± 0.05 | 0.38 ± 0.05 | 0.43 ± 0.06 | 0.40 ± 0.07 | 0.37 ± 0.05 | 0.37 ± 0.04 |
| | 0.33 ± 0.06 | 0.35 ± 0.03 | 0.34 ± 0.02 | 0.43 ± 0.04 | 0.49 ± 0.07 | 0.40 ± 0.05 | 0.46 ± 0.06 | 0.40 ± 0.04 |
| | 0.34 ± 0.05 | 0.34 ± 0.04 | 0.40 ± 0.03 | 0.42 ± 0.04 | 0.60 ± 0.08 | 0.51 ± 0.05 | 0.36 ± 0.04 | 0.51 ± 0.07 |

| | | | | | | | | |
|---|---|---|---|---|---|---|---|---|
| NDUFA8 | 0.9 ± 0.1 | 0.6 ± 0.1 | 0.8 ± 0.1 | 0.70 ± 0.09 | 0.9 ± 0.1 | 0.9 ± 0.1 | 1.2 ± 0.2 | 0.78 ± 0.09 |
| | 0.7 ± 0.2 | 0.9 ± 0.2 | 0.80 ± 0.08 | 0.71 ± 0.09 | 0.75 ± 0.09 | 0.8 ± 0.1 | 0.8 ± 0.1 | 0.9 ± 0.1 |
| | 0.60 ± 0.06 | 0.6 ± 0.1 | 0.8 ± 0.1 | 0.71 ± 0.06 | 0.8 ± 0.1 | 0.8 ± 0.1 | 0.78 ± 0.07 | 0.8 ± 0.2 |
| PDSW / NDUFB10 | 0.6 ± 0.1 | 0.62 ± 0.06 | 0.7 ± 0.1 | 0.54 ± 0.08 | 0.7 ± 0.1 | 0.58 ± 0.07 | 0.59 ± 0.09 | 0.7 ± 0.1 |
| | 0.6 ± 0.1 | 0.82 ± 0.09 | 0.8 ± 0.1 | 0.7 ± 0.1 | 0.6 ± 0.1 | 0.74 ± 0.08 | 0.68 ± 0.09 | 0.61 ± 0.09 |
| | 0.60 ± 0.09 | 0.69 ± 0.09 | 0.6 ± 0.1 | 0.61 ± 0.07 | 0.8 ± 0.2 | 0.7 ± 0.1 | 0.69 ± 0.06 | 0.76 ± 0.09 |
| NDUFA10 | 0.57 ± 0.05 | 0.63 ± 0.07 | 0.59 ± 0.04 | 0.7 ± 0.2 | 0.60 ± 0.04 | 0.70 ± 0.07 | 0.64 ± 0.07 | 0.57 ± 0.04 |
| | 0.63 ± 0.06 | 0.57 ± 0.05 | 0.66 ± 0.05 | 0.63 ± 0.06 | 0.8 ± 0.1 | 0.60 ± 0.04 | 0.66 ± 0.07 | 0.72 ± 0.07 |
| | 0.50 ± 0.05 | 0.57 ± 0.06 | 0.66 ± 0.08 | 0.72 ± 0.09 | 0.74 ± 0.07 | 0.61 ± 0.09 | 0.62 ± 0.05 | 0.60 ± 0.05 |
| NDUFS5 | 0.6 ± 0.2 | 0.41 ± 0.07 | 0.43 ± 0.05 | 0.42 ± 0.07 | 0.41 ± 0.05 | 0.45 ± 0.05 | 0.9 ± 0.1 | 0.8 ± 0.2 |
| | 0.52 ± 0.08 | 0.45 ± 0.08 | 0.52 ± 0.08 | 0.7 ± 0.1 | 0.65 ± 0.07 | 0.51 ± 0.09 | 0.56 ± 0.05 | 0.64 ± 0.05 |
| | 0.8 ± 0.2 | 0.5 ± 0.2 | 0.64 ± 0.09 | 0.52 ± 0.06 | 0.40 ± 0.03 | 0.55 ± 0.07 | 0.6 ± 0.1 | 0.42 ± 0.07 |
| NDUFA3 | 0.63 ± 0.09 | 0.77 ± 0.07 | 0.69 ± 0.06 | 0.61 ± 0.08 | 0.64 ± 0.05 | 0.65 ± 0.08 | 0.66 ± 0.06 | 0.7 ± 0.1 |
| | 0.66 ± 0.08 | 0.70 ± 0.07 | 0.65 ± 0.06 | 0.65 ± 0.07 | 0.68 ± 0.05 | 0.7 ± 0.1 | 0.57 ± 0.04 | 0.74 ± 0.09 |
| | 0.73 ± 0.08 | 0.9 ± 0.1 | 0.78 ± 0.07 | 0.8 ± 0.1 | 0.9 ± 0.1 | 0.58 ± 0.07 | 0.57 ± 0.09 | 0.7 ± 0.1 |
| NDUFB3 | 0.8 ± 0.1 | 0.81 ± 0.07 | 0.78 ± 0.05 | 0.85 ± 0.09 | 1.0 ± 0.2 | 1.0 ± 0.1 | 0.61 ± 0.07 | 0.72 ± 0.09 |
| | 0.77 ± 0.04 | 0.81 ± 0.05 | 0.63 ± 0.06 | 0.8 ± 0.1 | 0.9 ± 0.1 | 0.7 ± 0.1 | 0.77 ± 0.07 | 0.66 ± 0.06 |
| | 1.1 ± 0.1 | 0.9 ± 0.1 | 0.66 ± 0.05 | 0.9 ± 0.1 | 0.66 ± 0.05 | 1.01 ± 0.08 | 0.78 ± 0.07 | 1.1 ± 0.2 |
| NDUFC2 | 0.51 ± 0.08 | 0.42 ± 0.05 | 0.64 ± 0.08 | 0.46 ± 0.03 | 0.48 ± 0.06 | 0.48 ± 0.06 | 0.50 ± 0.05 | 0.52 ± 0.07 |
| | 0.43 ± 0.08 | 0.54 ± 0.09 | 0.45 ± 0.04 | 0.45 ± 0.03 | 0.44 ± 0.05 | 0.46 ± 0.04 | 0.49 ± 0.05 | 0.44 ± 0.03 |
| | 0.55 ± 0.07 | 0.54 ± 0.08 | 0.46 ± 0.07 | 0.60 ± 0.07 | 0.5 ± 0.1 | 0.53 ± 0.05 | 0.55 ± 0.06 | 0.58 ± 0.09 |
| NDUFB4 | 0.38 ± 0.04 | 0.52 ± 0.09 | 0.52 ± 0.05 | 0.44 ± 0.04 | 0.54 ± 0.04 | 0.54 ± 0.05 | 0.47 ± 0.04 | 0.46 ± 0.04 |
| | 0.53 ± 0.04 | 0.60 ± 0.09 | 0.51 ± 0.04 | 0.59 ± 0.04 | 0.48 ± 0.04 | 0.60 ± 0.06 | 0.56 ± 0.05 | 0.50 ± 0.03 |
| | 0.51 ± 0.07 | 0.46 ± 0.04 | 0.46 ± 0.04 | 0.44 ± 0.04 | 0.49 ± 0.04 | 0.52 ± 0.04 | 0.47 ± 0.06 | 0.51 ± 0.03 |
| NDUFA13 | 1.0 ± 0.3 | 0.8 ± 0.2 | 0.83 ± 0.08 | 1.2 ± 0.3 | 0.7 ± 0.1 | 1.0 ± 0.1 | 0.86 ± 0.06 | 1.1 ± 0.1 |
| | 1.0 ± 0.2 | 1.0 ± 0.1 | 1.0 ± 0.2 | 1.2 ± 0.2 | 1.0 ± 0.2 | 1.0 ± 0.1 | 0.9 ± 0.1 | 0.9 ± 0.1 |
| | 0.8 ± 0.2 | 0.8 ± 0.1 | 1.0 ± 0.1 | 1.1 ± 0.1 | 1.0 ± 0.1 | 1.2 ± 0.1 | 1.0 ± 0.2 | 0.9 ± 0.1 |
| NDUFB6 | 0.8 ± 0.1 | 0.69 ± 0.09 | 0.9 ± 0.2 | 0.72 ± 0.08 | 0.91 ± 0.07 | 1.1 ± 0.3 | 0.73 ± 0.07 | 1.1 ± 0.1 |
| | 0.89 ± 0.07 | 0.70 ± 0.07 | 0.81 ± 0.06 | 0.83 ± 0.08 | 0.90 ± 0.08 | 0.69 ± 0.07 | 0.73 ± 0.09 | 0.57 ± 0.06 |
| | 0.74 ± 0.08 | 0.77 ± 0.06 | 1.03 ± 0.08 | 0.71 ± 0.09 | 0.69 ± 0.07 | 0.77 ± 0.05 | 0.76 ± 0.06 | 0.9 ± 0.1 |
| NDUFB7 | 0.8 ± 0.1 | 0.71 ± 0.09 | 1.0 ± 0.1 | 0.8 ± 0.1 | 0.79 ± 0.09 | 0.8 ± 0.1 | 0.9 ± 0.1 | 0.9 ± 0.1 |
| | 0.8 ± 0.1 | 0.7 ± 0.1 | 0.7 ± 0.1 | 0.8 ± 0.1 | 0.8 ± 0.1 | 0.9 ± 0.1 | 0.9 ± 0.1 | 0.73 ± 0.09 |
| | 0.8 ± 0.1 | 0.8 ± 0.1 | 0.9 ± 0.1 | 0.80 ± 0.09 | 0.7 ± 0.1 | 0.74 ± 0.08 | 0.9 ± 0.1 | 0.8 ± 0.1 |
| NDUFB9 | 0.51 ± 0.06 | 0.45 ± 0.07 | 0.53 ± 0.09 | 0.6 ± 0.1 | 0.47 ± 0.07 | 0.57 ± 0.09 | 0.49 ± 0.04 | 0.48 ± 0.07 |
| | 0.59 ± 0.06 | 0.55 ± 0.05 | 0.50 ± 0.04 | 0.47 ± 0.04 | 0.6 ± 0.1 | 0.51 ± 0.07 | 0.52 ± 0.05 | 0.44 ± 0.05 |
| | 0.51 ± 0.05 | 0.61 ± 0.09 | 0.57 ± 0.05 | 0.50 ± 0.05 | 0.56 ± 0.08 | 0.61 ± 0.06 | 0.52 ± 0.06 | 0.58 ± 0.07 |
| NDUFB2 | 0.8 ± 0.1 | 0.6 ± 0.1 | 0.7 ± 0.1 | 0.46 ± 0.04 | 0.7 ± 0.1 | 0.9 ± 0.2 | 0.61 ± 0.08 | 0.9 ± 0.2 |
| | 0.7 ± 0.2 | 0.72 ± 0.09 | 0.6 ± 0.1 | 0.7 ± 0.1 | 0.7 ± 0.1 | 0.8 ± 0.1 | 0.8 ± 0.1 | 0.8 ± 0.1 |
| | 1.0 ± 0.1 | 0.48 ± 0.05 | 0.62 ± 0.06 | 0.67 ± 0.08 | 0.8 ± 0.1 | 0.60 ± 0.06 | 0.61 ± 0.07 | 0.56 ± 0.07 |

| | | | | | | | | |
|---|---|---|---|---|---|---|---|---|
| NDUFB8 | 0.9 ± 0.2 | 0.8 ± 0.1 | 0.80 ± 0.09 | 0.68 ± 0.07 | 0.76 ± 0.07 | 0.69 ± 0.07 | 0.79 ± 0.09 | 0.9 ± 0.1 |
| | 0.67 ± 0.08 | 0.8 ± 0.1 | 0.67 ± 0.09 | 0.72 ± 0.06 | 0.92 ± 0.08 | 0.8 ± 0.1 | 0.85 ± 0.07 | 0.76 ± 0.06 |
| | 0.6 ± 0.1 | 0.8 ± 0.1 | 0.85 ± 0.08 | 0.76 ± 0.09 | 0.8 ± 0.1 | 0.75 ± 0.08 | 0.8 ± 0.1 | 0.75 ± 0.09 |
| ESSS / NDUFB11 | 1.03 ± 0.09 | 1.00 ± 0.08 | 0.8 ± 0.1 | 0.93 ± 0.09 | 1.4 ± 0.2 | 1.0 ± 0.1 | 1.2 ± 0.2 | 0.9 ± 0.1 |
| | 0.9 ± 0.1 | 1.2 ± 0.1 | 0.9 ± 0.2 | 0.98 ± 0.09 | 1.1 ± 0.2 | 0.94 ± 0.11 | 1.0 ± 0.2 | 1.0 ± 0.2 |
| | 1.03 ± 0.05 | 1.0 ± 0.1 | 0.91 ± 0.08 | 1.1 ± 0.1 | 0.89 ± 0.08 | 0.96 ± 0.09 | 0.88 ± 0.05 | 1.10 ± 0.09 |
| KFYI / NDUFC1 | 0.33 ± 0.04 | 0.6 ± 0.1 | 0.42 ± 0.05 | 0.56 ± 0.09 | 0.41 ± 0.06 | 0.46 ± 0.04 | 0.38 ± 0.04 | 0.58 ± 0.08 |
| | 0.48 ± 0.07 | 0.49 ± 0.07 | 0.40 ± 0.05 | 0.41 ± 0.04 | 0.37 ± 0.05 | 0.41 ± 0.08 | 0.37 ± 0.07 | 0.39 ± 0.04 |
| | 0.46 ± 0.08 | 0.35 ± 0.04 | 0.46 ± 0.09 | 0.41 ± 0.04 | 0.43 ± 0.05 | 0.47 ± 0.07 | 0.51 ± 0.06 | 0.4 ± 0.2 |
| MNLL/ NDUFB1 | 0.64 ± 0.06 | 0.53 ± 0.06 | 0.68 ± 0.09 | 0.60 ± 0.04 | 0.79 ± 0.09 | 0.8 ± 0.1 | 0.74 ± 0.06 | 0.63 ± 0.08 |
| | 0.64 ± 0.05 | 0.61 ± 0.07 | 0.62 ± 0.07 | 0.63 ± 0.05 | 0.65 ± 0.06 | 0.8 ± 0.1 | 0.57 ± 0.05 | 0.69 ± 0.05 |
| | 0.75 ± 0.08 | 0.57 ± 0.09 | 0.67 ± 0.08 | 0.64 ± 0.08 | 0.80 ± 0.09 | 0.82 ± 0.07 | 0.56 ± 0.05 | 0.60 ± 0.07 |
| MWFE / NDUFA1 | 0.49 ± 0.05 | 0.32 ± 0.03 | 0.38 ± 0.04 | 0.48 ± 0.08 | 0.54 ± 0.09 | 0.5 ± 0.1 | 0.6 ± 0.1 | 0.57 ± 0.09 |
| | 0.4 ± 0.1 | 0.65 ± 0.9 | 0.48 ± 0.07 | 0.49 ± 0.06 | 0.52 ± 0.06 | 0.42 ± 0.08 | 0.42 ± 0.05 | 0.6 ± 0.1 |
| | 0.44 ± 0.08 | 0.33 ± 0.02 | 0.6 ± 0.1 | 0.47 ± 0.06 | 0.5 ± 0.1 | 0.39 ± 0.04 | 0.40 ± 0.05 | 0.6 ± 0.2 |

The NDUFS2 (49 kDa in the ovine naming, Table S2) subunit has higher average RMSD values (between 0.6 and 1.0 nm). This is probably caused by the absence of the Q module subunits that usually surround and stabilize this subunit, and, consequently, the solvent exposure. The NDUFA11, NDUFB5, NDUFAB1, NDUFC1 (KFYI) and the NDUFA1 (MWFE) subunits have the lowest average RMSD values between the supernumerary subunits (below 0.5 nm). On the other hand, the NDUFA13, NDUFB6, NDUFB7 and NDUFB11 (ESSS) subunits shows the higher average RMSD values (over 0.8 nm) because of the large oscillations of their C-terminals (see below). The other supernumerary subunits have generally intermediate RMSD values (between 0.5 and 0.8 nm).

**Fig. S10. Backbone beads RMSF *vs.* residue number of the P+ module subunits in the wtO$^{op}$ structures.**
Each panel label corresponds to the equivalent subunit in Table S1. The RMSF values corresponding to the three replicas are in brown, maroon, and crimson, respectively.

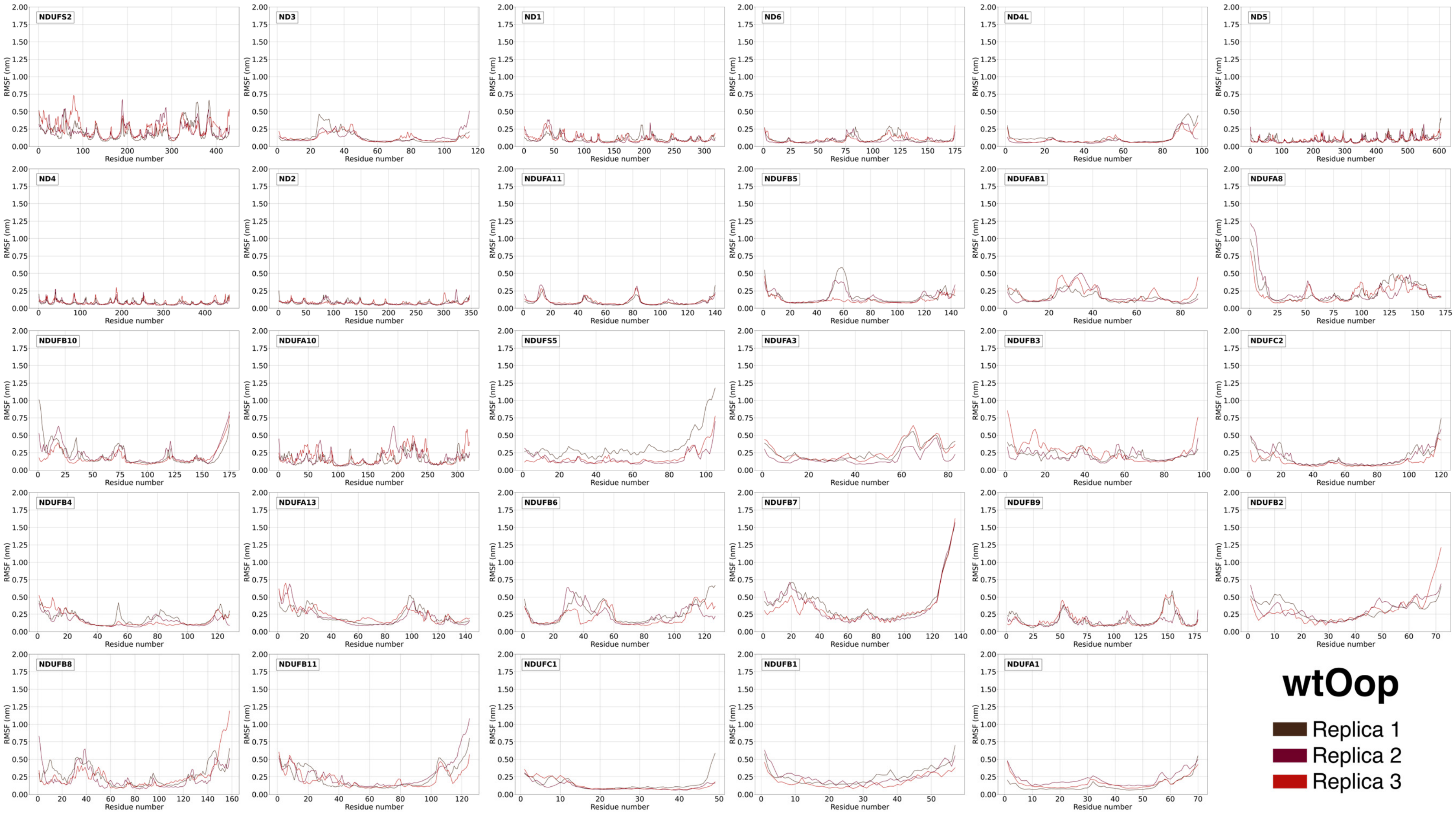

**Fig. S11. Backbone beads RMSF *vs.* residue number plots of the P+ module subunits in the wtO$^{cl}$ structures.**
Each panel label corresponds to the equivalent subunit in Table S1. The RMSF values corresponding to the three replicas are in brown, maroon, and crimson, respectively.

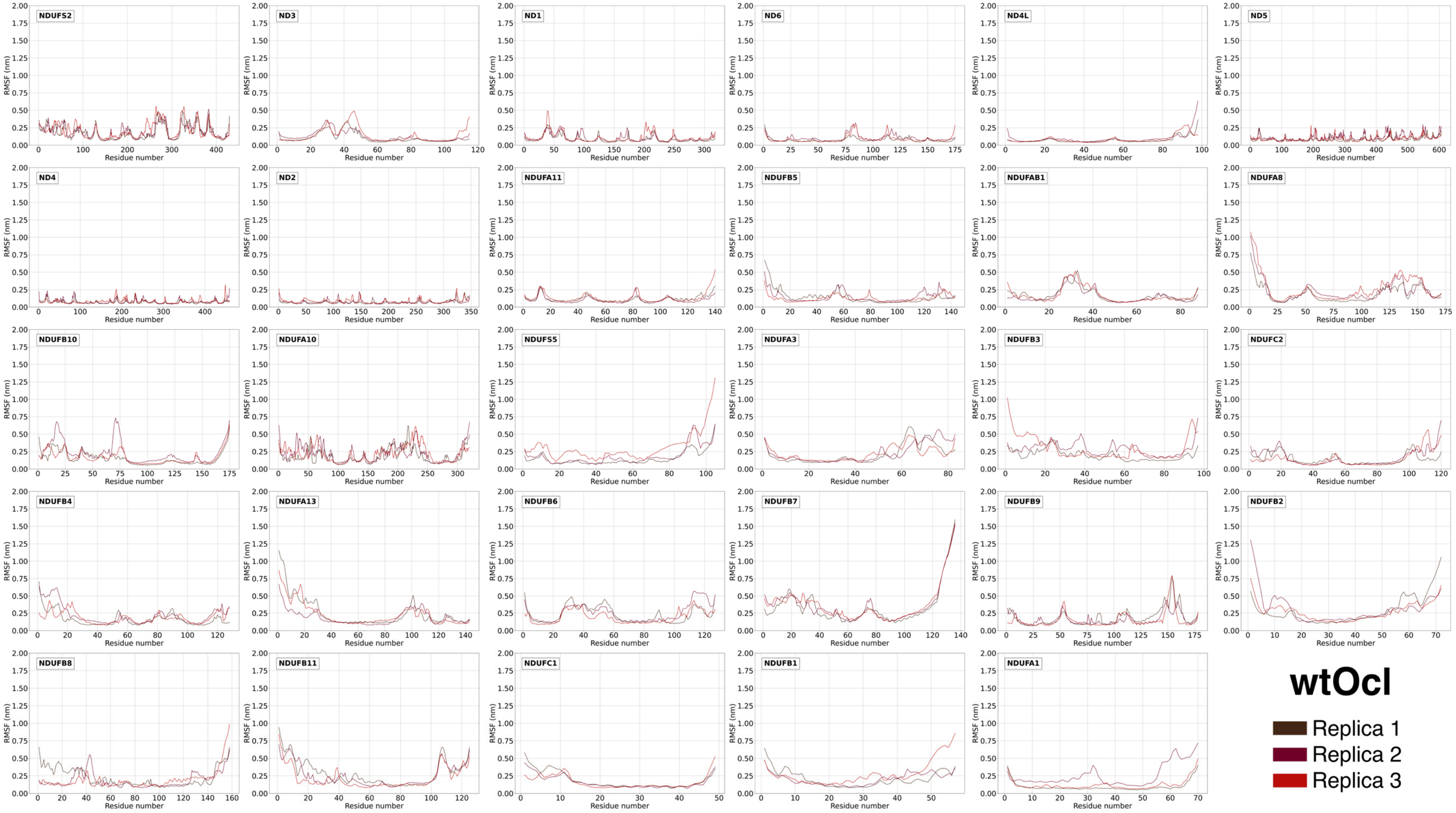

**Fig. S12. Backbone beads RMSF *vs.* residue number plots of the P+ module subunits in the wtH$^{op}$ structures.**
Each panel label corresponds to the equivalent subunit in Table S1. The RMSF values corresponding to the three replicas are in brown, maroon, and crimson, respectively.

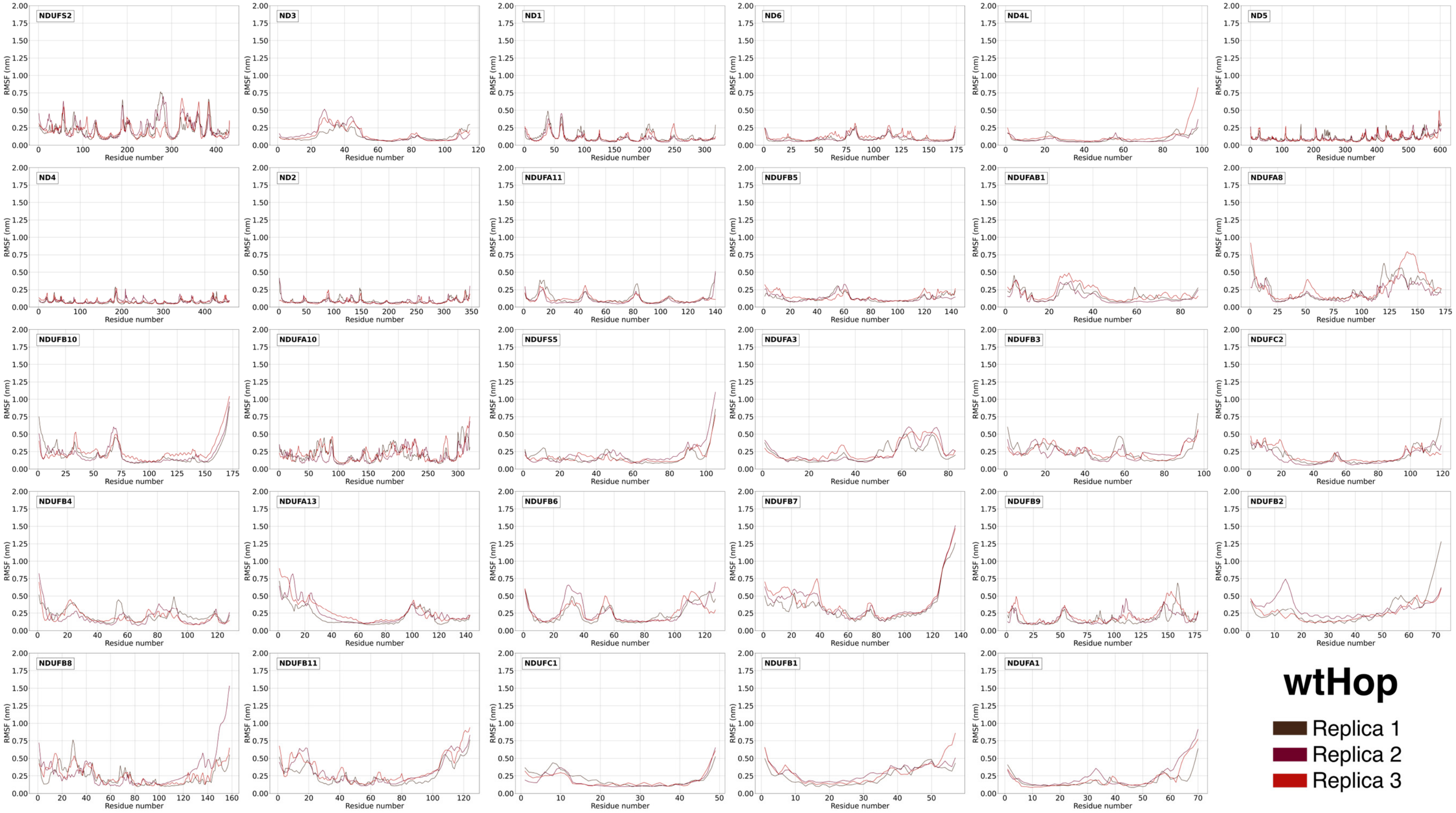

**Fig. S13. Backbone beads RMSF *vs.* residue number plots of the P+ module subunits in the wtH$^{cl}$ structures.**
Each panel label corresponds to the equivalent subunit in Table S1. The RMSF values corresponding to the three replicas are in brown, maroon, and crimson, respectively.

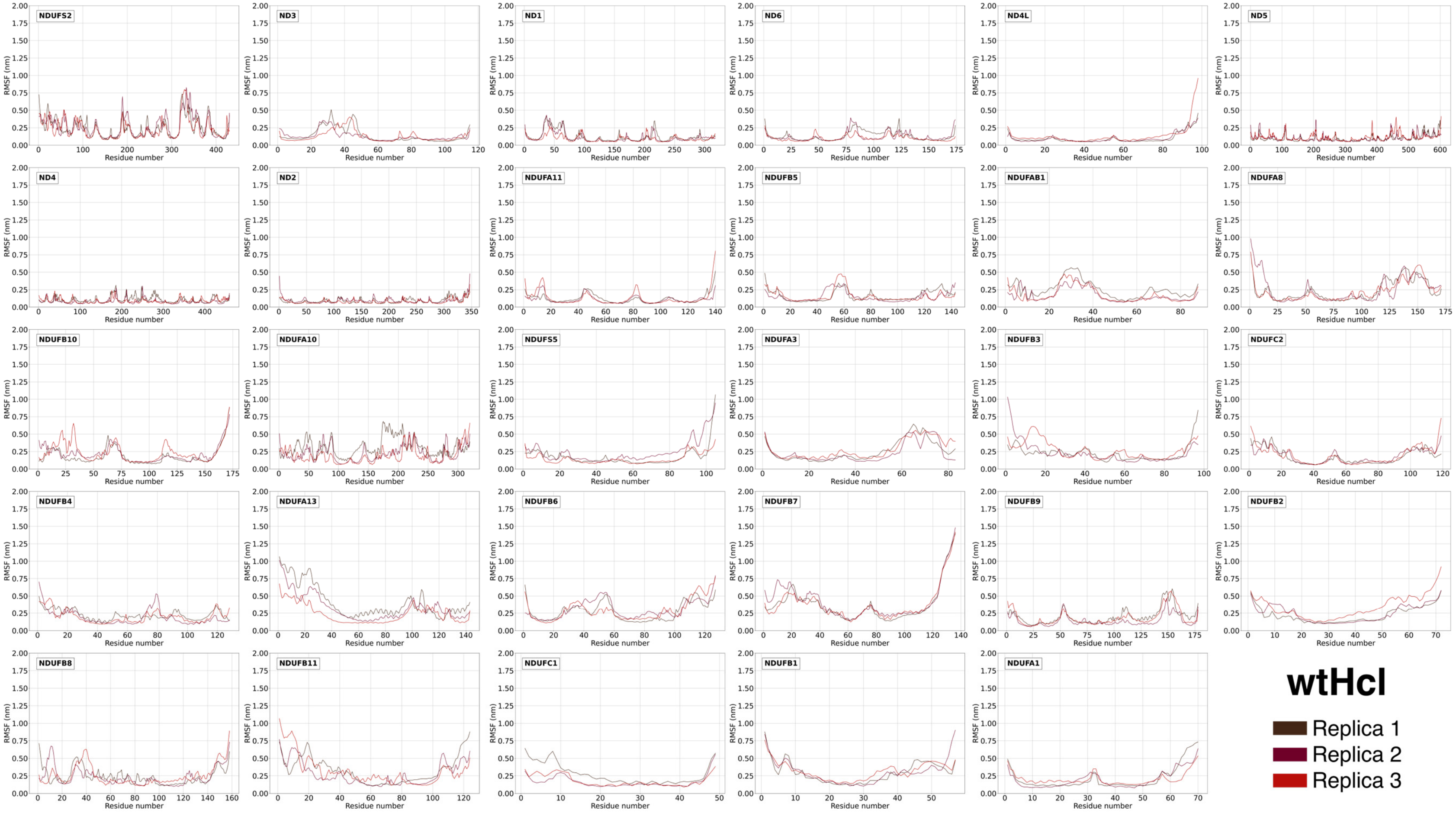

**Fig. S14. Backbone beads RMSF *vs.* residue number plots of the P+ module subunits in the m1H$^{op}$ structures.**
Each panel label corresponds to the equivalent subunit in Table S1. The RMSF values corresponding to the three replicas are in navy, royal blue, and fountain blue, respectively.

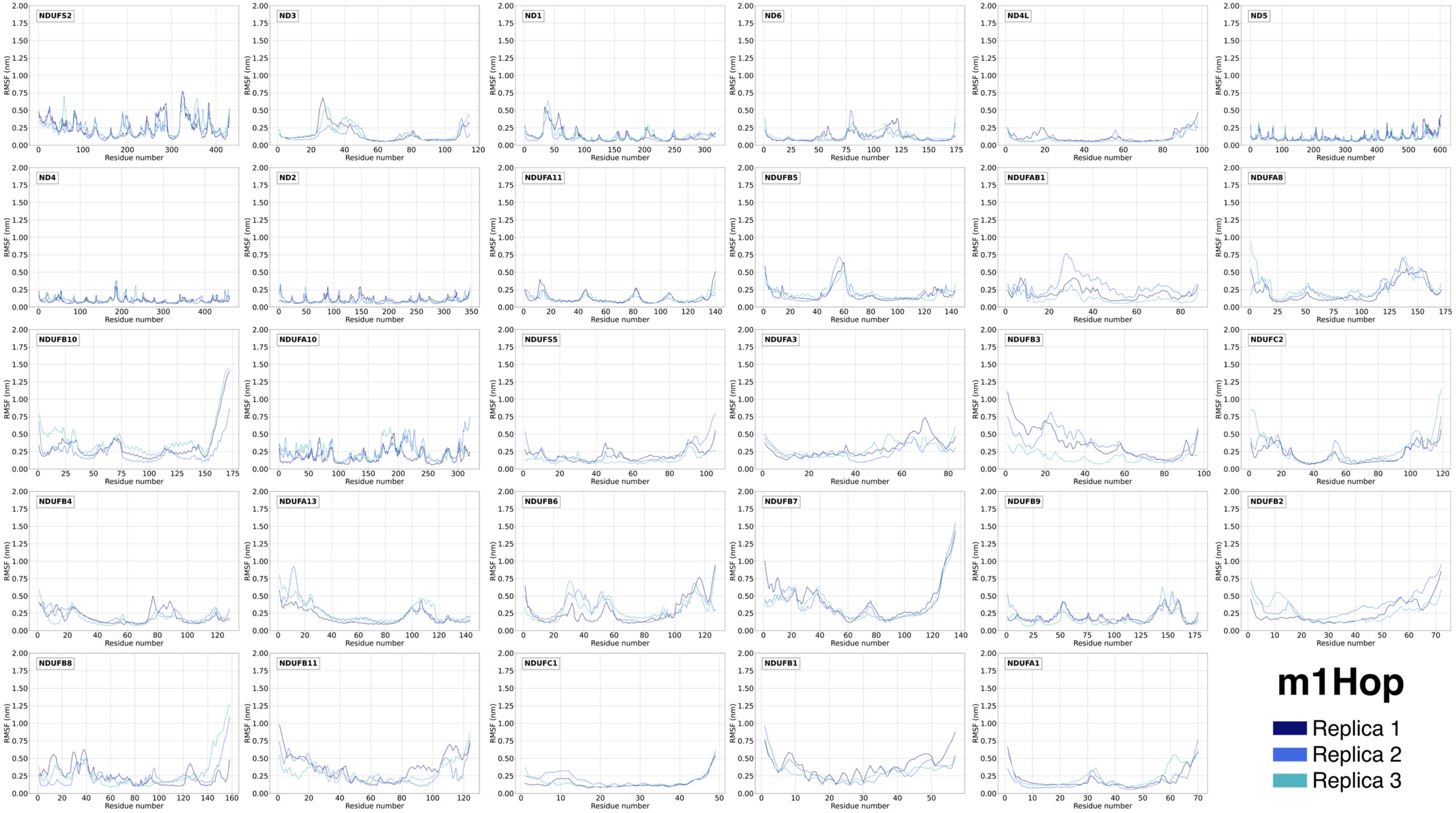

**Fig. S15. Backbone beads RMSF *vs.* residue number plots of the P+ module subunits in the m1H$^{cl}$ structures.**
Each panel label corresponds to the equivalent subunit in Table S1. The RMSF values corresponding to the three replicas are in navy, royal blue, and fountain blue, respectively.

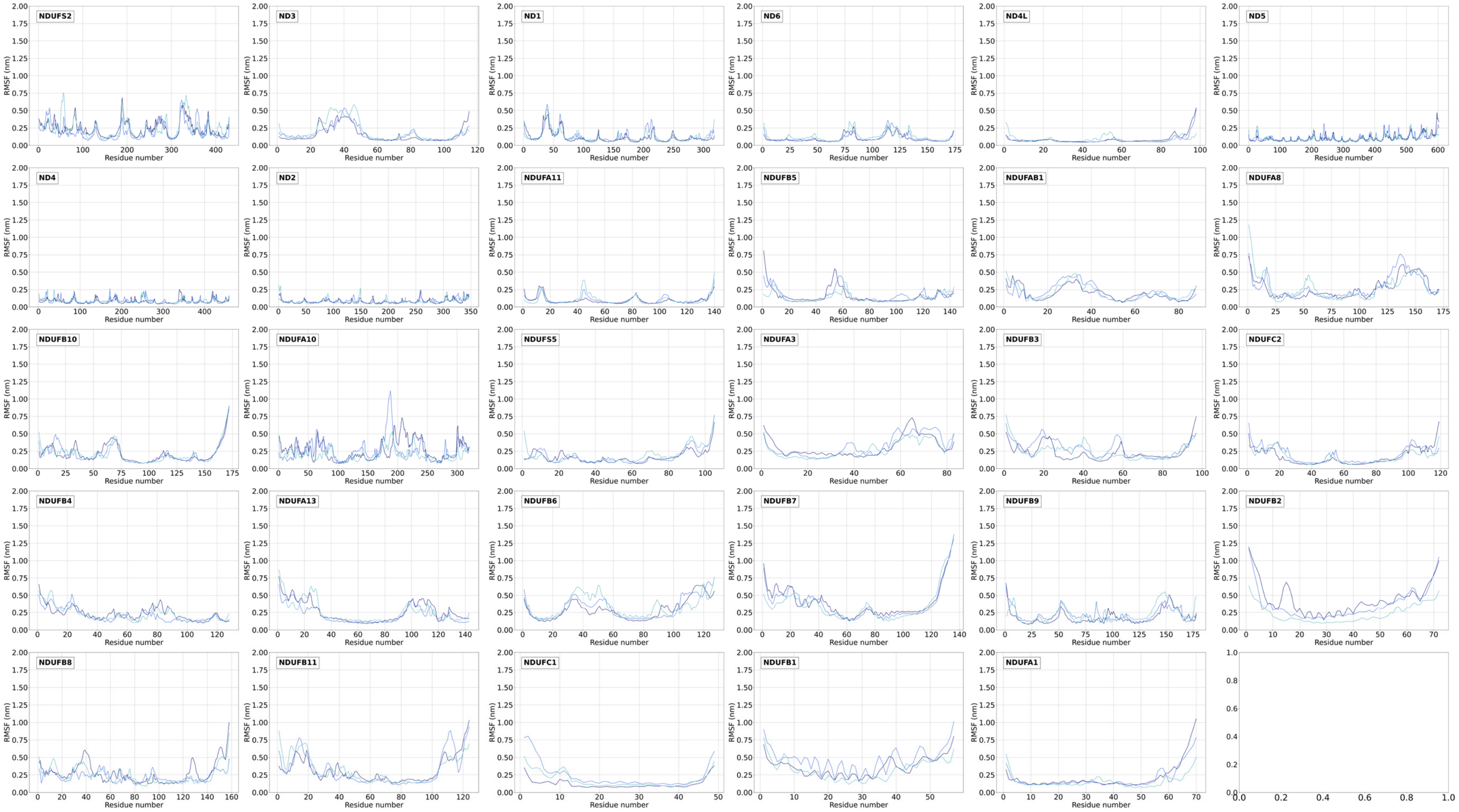

**Fig. S16. Backbone beads RMSF *vs.* residue number plots of the P+ module subunits in the m3H$^{op}$ structures.**
Each panel label corresponds to the equivalent subunit in Table S1. The RMSF values corresponding to the three replicas are in pine, dark lime, and lime, respectively.

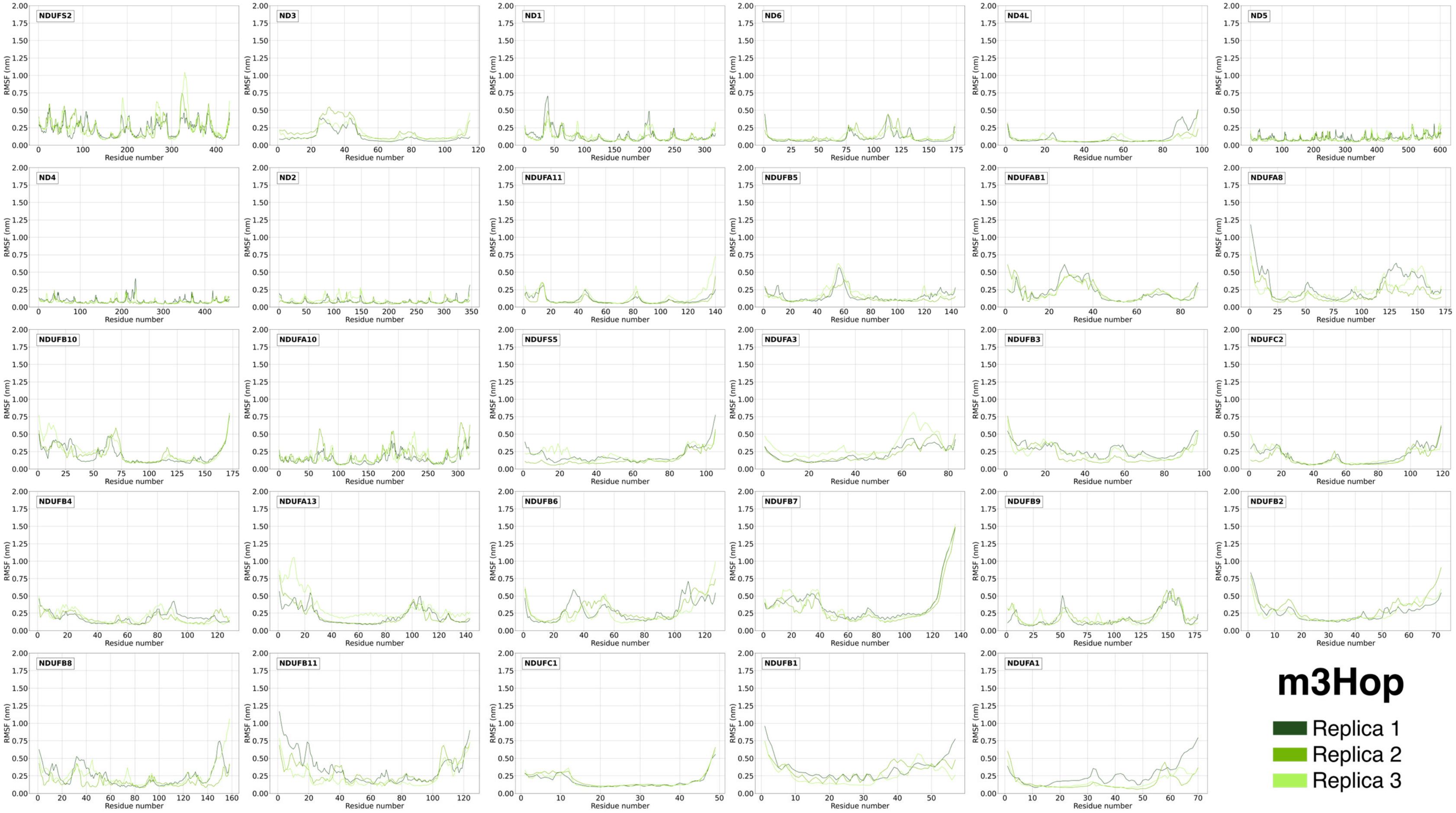

**Fig. S17. Backbone beads RMSF *vs.* residue number plots of the P+ module subunits in the m3H$^{cl}$ structures.**
Each panel label corresponds to the equivalent subunit in Table S1. The RMSF values corresponding to the three replicas are in pine, dark lime, and lime, respectively.

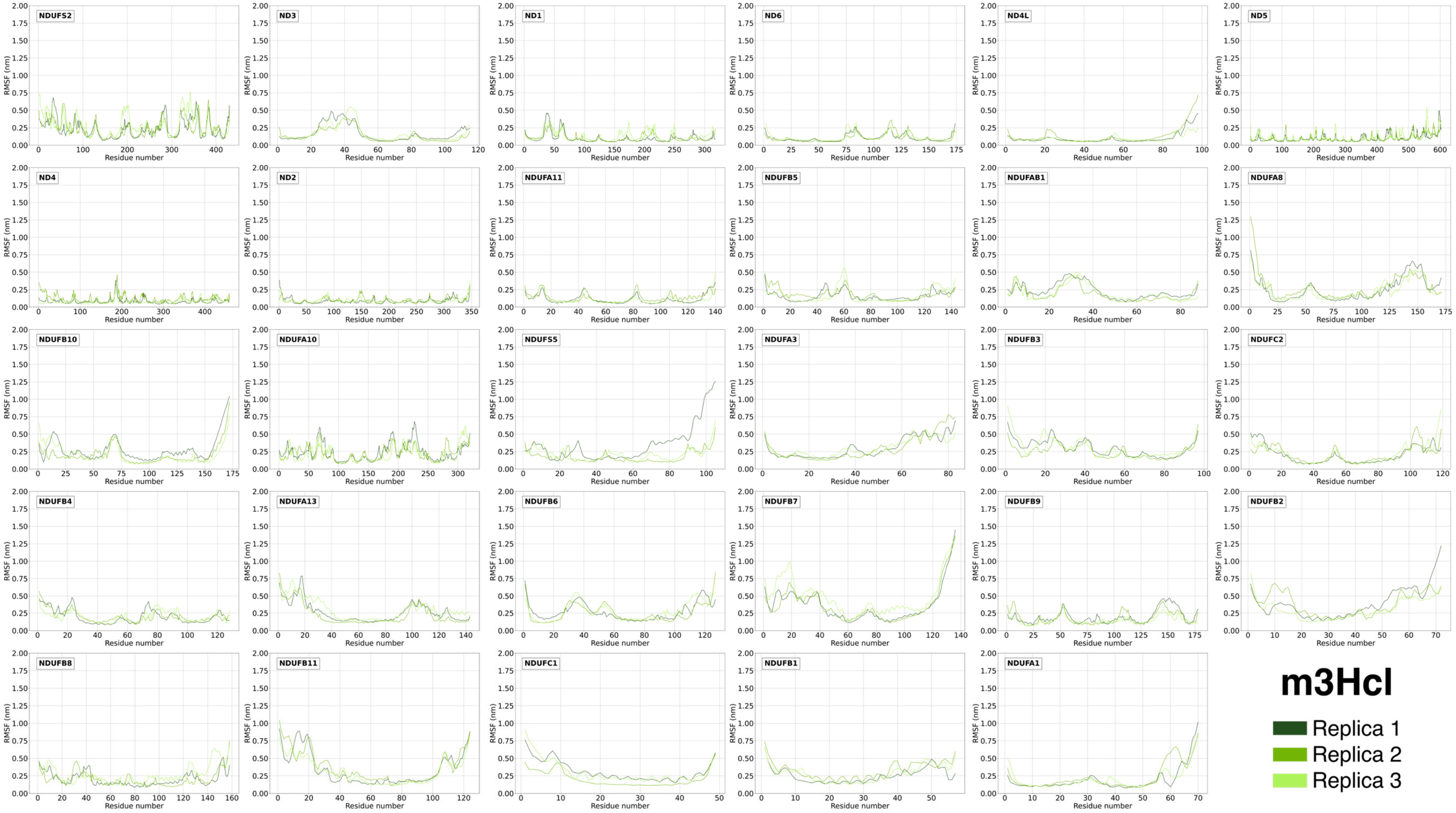

**Table S5. Average and standard deviation of RMSF values (in nm) of backbone beads for each subunit of ovine and human P+ module.** For each subunit, the ovine and human abbreviation (see Table S1) is reported. Each subunits line corresponds to a different replica.

| Human/Ovine subunit | wtO$^{op}$ | wtO$^{cl}$ | wtH$^{op}$ | wtH$^{cl}$ | m1H$^{op}$ | m1H$^{cl}$ | m3H$^{op}$ | m3H$^{cl}$ |
|---|---|---|---|---|---|---|---|---|
| 49 kDa / NDUFS2 | 0.3 ± 0.1 | 0.3 ± 0.1 | 0.3 ± 0.1 | 0.3 ± 0.2 | 0.3 ± 0.2 | 0.3 ± 0.1 | 0.3 ± 0.1 | 0.3 ± 0.2 |
| | 0.3 ± 0.1 | 0.3 ± 0.1 | 0.3 ± 0.2 | 0.3 ± 0.2 | 0.3 ± 0.2 | 0.3 ± 0.1 | 0.3 ± 0.2 | 0.3 ± 0.1 |
| | 0.3 ± 0.1 | 0.3 ± 0.1 | 0.3 ± 0.1 | 0.3 ± 0.2 | 0.3 ± 0.1 | 0.3 ± 0.1 | 0.3 ± 0.2 | 0.3 ± 0.2 |
| ND3 | 0.14 ± 0.09 | 0.2 ± 0.1 | 0.2 ± 0.1 | 0.2 ± 0.1 | 0.2 ± 0.1 | 0.2 ± 0.1 | 0.15 ± 0.09 | 0.2 ± 0.1 |
| | 0.2 ± 0.1 | 0.2 ± 0.1 | 0.2 ± 0.1 | 0.15 ± 0.07 | 0.15 ± 0.07 | 0.2 ± 0.1 | 0.2 ± 0.1 | 0.2 ± 0.1 |
| | 0.16 ± 0.07 | 0.2 ± 0.1 | 0.2 ± 0.1 | 0.15 ± 0.09 | 0.2 ± 0.1 | 0.2 ± 0.1 | 0.2 ± 0.1 | 0.2 ± 0.1 |
| ND1 | 0.12 ± 0.05 | 0.13 ± 0.08 | 0.14 ± 0.08 | 0.15 ± 0.08 | 0.14 ± 0.09 | 0.14 ± 0.08 | 0.1 ± 0.1 | 0.12 ± 0.08 |
| | 0.13 ± 0.07 | 0.13 ± 0.07 | 0.12 ± 0.07 | 0.13 ± 0.07 | 0.14 ± 0.07 | 0.15 ± 0.09 | 0.13 ± 0.08 | 0.13 ± 0.07 |
| | 0.14 ± 0.07 | 0.14 ± 0.08 | 0.14 ± 0.07 | 0.10 ± 0.05 | 0.16 ± 0.09 | 0.14 ± 0.08 | 0.13 ± 0.06 | 0.15 ± 0.07 |
| ND6 | 0.12 ± 0.05 | 0.12 ± 0.05 | 0.12 ± 0.05 | 0.15 ± 0.07 | 0.13 ± 0.07 | 0.12 ± 0.05 | 0.12 ± 0.07 | 0.12 ± 0.06 |
| | 0.11 ± 0.05 | 0.11 ± 0.05 | 0.10 ± 0.05 | 0.14 ± 0.07 | 0.13 ± 0.07 | 0.14 ± 0.07 | 0.16 ± 0.09 | 0.11 ± 0.06 |
| | 0.13 ± 0.06 | 0.11 ± 0.05 | 0.13 ± 0.05 | 0.12 ± 0.06 | 0.14 ± 0.08 | 0.17 ± 0.08 | 0.14 ± 0.06 | 0.14 ± 0.07 |
| ND4L | 0.2 ± 0.1 | 0.10 ± 0.06 | 0.11 ± 0.05 | 0.11 ± 0.07 | 0.13 ± 0.09 | 0.11 ± 0.08 | 0.1 ± 0.1 | 0.2 ± 0.1 |
| | 0.10 ± 0.07 | 0.09 ± 0.08 | 0.09 ± 0.06 | 0.11 ± 0.08 | 0.11 ± 0.08 | 0.09 ± 0.07 | 0.10 ± 0.05 | 0.2 ± 0.1 |
| | 0.13 ± 0.09 | 0.1 ± 0.1 | 0.1 ± 0.1 | 0.1 ± 0.1 | 0.09 ± 0.06 | 0.10 ± 0.06 | 0.10 ± 0.06 | 0.10 ± 0.05 |
| ND5 | 0.13 ± 0.07 | 0.12 ± 0.05 | 0.12 ± 0.06 | 0.11 ± 0.06 | 0.12 ± 0.06 | 0.13 ± 0.07 | 0.12 ± 0.05 | 0.13 ± 0.07 |
| | 0.10 ± 0.05 | 0.14 ± 0.05 | 0.11 ± 0.05 | 0.12 ± 0.06 | 0.12 ± 0.06 | 0.12 ± 0.06 | 0.11 ± 0.05 | 0.12 ± 0.06 |
| | 0.11 ± 0.05 | 0.12 ± 0.05 | 0.11 ± 0.06 | 0.13 ± 0.05 | 0.12 ± 0.06 | 0.11 ± 0.05 | 0.12 ± 0.06 | 0.13 ± 0.07 |
| ND4 | 0.11 ± 0.04 | 0.10 ± 0.04 | 0.08 ± 0.03 | 0.11 ± 0.05 | 0.09 ± 0.04 | 0.10 ± 0.04 | 0.09 ± 0.04 | 0.09 ± 0.05 |
| | 0.10 ± 0.04 | 0.10 ± 0.04 | 0.09 ± 0.03 | 0.09 ± 0.04 | 0.10 ± 0.04 | 0.09 ± 0.04 | 0.09 ± 0.04 | 0.13 ± 0.06 |
| | 0.11 ± 0.05 | 0.09 ± 0.04 | 0.09 ± 0.04 | 0.10 ± 0.04 | 0.11 ± 0.06 | 0.09 ± 0.04 | 0.09 ± 0.04 | 0.09 ± 0.04 |
| ND2 | 0.09 ± 0.04 | 0.09 ± 0.04 | 0.10 ± 0.04 | 0.11 ± 0.04 | 0.11 ± 0.05 | 0.09 ± 0.04 | 0.09 ± 0.04 | 0.10 ± 0.05 |
| | 0.10 ± 0.04 | 0.09 ± 0.04 | 0.09 ± 0.05 | 0.11 ± 0.05 | 0.11 ± 0.05 | 0.09 ± 0.04 | 0.09 ± 0.03 | 0.12 ± 0.04 |
| | 0.10 ± 0.04 | 0.09 ± 0.04 | 0.09 ± 0.04 | 0.09 ± 0.04 | 0.09 ± 0.04 | 0.09 ± 0.05 | 0.11 ± 0.05 | 0.12 ± 0.05 |
| NDUFA11 | 0.11 ± 0.06 | 0.12 ± 0.06 | 0.14 ± 0.09 | 0.13 ± 0.08 | 0.12 ± 0.08 | 0.11 ± 0.07 | 0.13 ± 0.07 | 0.13 ± 0.08 |
| | 0.10 ± 0.06 | 0.12 ± 0.05 | 0.11 ± 0.07 | 0.12 ± 0.06 | 0.12 ± 0.06 | 0.13 ± 0.08 | 0.10 ± 0.07 | 0.15 ± 0.06 |
| | 0.12 ± 0.07 | 0.16 ± 0.07 | 0.13 ± 0.06 | 0.1 ± 0.1 | 0.13 ± 0.06 | 0.11 ± 0.08 | 0.2 ± 0.1 | 0.13 ± 0.07 |
| NDUFB5 | 0.2 ± 0.1 | 0.18 ± 0.09 | 0.14 ± 0.05 | 0.18 ± 0.09 | 0.2 ± 0.1 | 0.2 ± 0.1 | 0.17 ± 0.09 | 0.19 ± 0.08 |
| | 0.19 ± 0.09 | 0.18 ± 0.09 | 0.16 ± 0.06 | 0.17 ± 0.08 | 0.22 ± 0.08 | 0.18 ± 0.09 | 0.16 ± 0.08 | 0.21 ± 0.08 |
| | 0.17 ± 0.06 | 0.16 ± 0.08 | 0.18 ± 0.08 | 0.2 ± 0.1 | 0.20 ± 0.08 | 0.17 ± 0.08 | 0.2 ± 0.1 | 0.2 ± 0.1 |
| NDUFAB1 | 0.19 ± 0.09 | 0.21 ± 0.09 | 0.18 ± 0.08 | 0.2 ± 0.1 | 0.22 ± 0.09 | 0.2 ± 0.1 | 0.2 ± 0.1 | 0.2 ± 0.1 |
| | 0.2 ± 0.1 | 0.18 ± 0.09 | 0.14 ± 0.09 | 0.2 ± 0.1 | 0.3 ± 0.1 | 0.2 ± 0.1 | 0.3 ± 0.1 | 0.2 ± 0.1 |
| | 0.2 ± 0.1 | 0.2 ± 0.1 | 0.2 ± 0.1 | 0.2 ± 0.1 | 0.20 ± 0.09 | 0.2 ± 0.1 | 0.2 ± 0.1 | 0.2 ± 0.1 |

| | | | | | | | | |
|---|---|---|---|---|---|---|---|---|
| NDUFA8 | 0.3 ± 0.2 | 0.2 ± 0.1 | 0.3 ± 0.2 | 0.3 ± 0.2 | 0.3 ± 0.2 | 0.3 ± 0.2 | 0.4 ± 0.2 | 0.3 ± 0.2 |
| | 0.3 ± 0.2 | 0.3 ± 0.2 | 0.3 ± 0.1 | 0.3 ± 0.2 | 0.3 ± 0.2 | 0.3 ± 0.2 | 0.2 ± 0.2 | 0.3 ± 0.2 |
| | 0.3 ± 0.1 | 0.3 ± 0.2 | 0.3 ± 0.2 | 0.3 ± 0.1 | 0.3 ± 0.2 | 0.3 ± 0.2 | 0.3 ± 0.1 | 0.3 ± 0.2 |
| PDSW / NDUFB10 | 0.3 ± 0.1 | 0.2 ± 0.1 | 0.3 ± 0.1 | 0.2 ± 0.1 | 0.3 ± 0.2 | 0.2 ± 0.1 | 0.2 ± 0.1 | 0.3 ± 0.2 |
| | 0.3 ± 0.1 | 0.3 ± 0.2 | 0.2 ± 0.2 | 0.3 ± 0.2 | 0.3 ± 0.2 | 0.2 ± 0.1 | 0.2 ± 0.1 | 0.2 ± 0.1 |
| | 0.3 ± 0.2 | 0.2 ± 0.1 | 0.3 ± 0.2 | 0.3 ± 0.1 | 0.4 ± 0.2 | 0.2 ± 0.1 | 0.3 ± 0.2 | 0.2 ± 0.1 |
| NDUFA10 | 0.2 ± 0.1 | 0.3 ± 0.1 | 0.2 ± 0.1 | 0.4 ± 0.2 | 0.21 ± 0.09 | 0.3 ± 0.2 | 0.2 ± 0.1 | 0.3 ± 0.1 |
| | 0.2 ± 0.1 | 0.3 ± 0.1 | 0.2 ± 0.1 | 0.2 ± 0.1 | 0.3 ± 0.1 | 0.3 ± 0.2 | 0.3 ± 0.1 | 0.3 ± 0.1 |
| | 0.2 ± 0.1 | 0.3 ± 0.1 | 0.3 ± 0.1 | 0.3 ± 0.2 | 0.3 ± 0.1 | 0.3 ± 0.1 | 0.20 ± 0.09 | 0.2 ± 0.1 |
| NDUFS5 | 0.4 ± 0.2 | 0.2 ± 0.1 | 0.2 ± 0.1 | 0.2 ± 0.1 | 0.20 ± 0.08 | 0.19 ± 0.09 | 0.3 ± 0.2 | 0.4 ± 0.3 |
| | 0.2 ± 0.1 | 0.2 ± 0.1 | 0.2 ± 0.2 | 0.3 ± 0.2 | 0.3 ± 0.2 | 0.3 ± 0.1 | 0.2 ± 0.1 | 0.2 ± 0.1 |
| | 0.3 ± 0.2 | 0.3 ± 0.2 | 0.2 ± 0.1 | 0.17 ± 0.08 | 0.14 ± 0.06 | 0.2 ± 0.1 | 0.3 ± 0.1 | 0.2 ± 0.1 |
| NDUFA3 | 0.3 ± 0.1 | 0.3 ± 0.1 | 0.2 ± 0.1 | 0.3 ± 0.2 | 0.4 ± 0.2 | 0.3 ± 0.1 | 0.3 ± 0.1 | 0.4 ± 0.2 |
| | 0.23 ± 0.09 | 0.3 ± 0.1 | 0.3 ± 0.1 | 0.3 ± 0.1 | 0.3 ± 0.1 | 0.4 ± 0.2 | 0.3 ± 0.1 | 0.4 ± 0.2 |
| | 0.3 ± 0.1 | 0.3 ± 0.1 | 0.3 ± 0.1 | 0.3 ± 0.2 | 0.4 ± 0.1 | 0.3 ± 0.1 | 0.4 ± 0.2 | 0.3 ± 0.1 |
| NDUFB3 | 0.3 ± 0.2 | 0.3 ± 0.1 | 0.3 ± 0.1 | 0.3 ± 0.1 | 0.4 ± 0.2 | 0.3 ± 0.1 | 0.3 ± 0.2 | 0.3 ± 0.1 |
| | 0.21 ± 0.09 | 0.3 ± 0.1 | 0.3 ± 0.1 | 0.3 ± 0.2 | 0.4 ± 0.2 | 0.4 ± 0.1 | 0.3 ± 0.2 | 0.30 ± 0.09 |
| | 0.4 ± 0.1 | 0.4 ± 0.2 | 0.3 ± 0.1 | 0.4 ± 0.2 | 0.21 ± 0.09 | 0.3 ± 0.1 | 0.3 ± 0.1 | 0.4 ± 0.2 |
| NDUFC2 | 0.2 ± 0.1 | 0.19 ± 0.09 | 0.2 ± 0.1 | 0.2 ± 0.1 | 0.2 ± 0.1 | 0.2 ± 0.1 | 0.2 ± 0.1 | 0.2 ± 0.1 |
| | 0.2 ± 0.1 | 0.2 ± 0.2 | 0.2 ± 0.1 | 0.2 ± 0.1 | 0.2 ± 0.1 | 0.2 ± 0.1 | 0.2 ± 0.1 | 0.2 ± 0.1 |
| | 0.2 ± 0.1 | 0.2 ± 0.1 | 0.2 ± 0.1 | 0.2 ± 0.2 | 0.3 ± 0.2 | 0.2 ± 0.1 | 0.2 ± 0.1 | 0.2 ± 0.1 |
| NDUFB4 | 0.2 ± 0.1 | 0.3 ± 0.1 | 0.24 ± 0.08 | 0.24 ± 0.08 | 0.2 ± 0.1 | 0.3 ± 0.1 | 0.21 ± 0.08 | 0.2 ± 0.1 |
| | 0.20 ± 0.08 | 0.3 ± 0.1 | 0.2 ± 0.1 | 0.2 ± 0.1 | 0.2 ± 0.1 | 0.2 ± 0.1 | 0.21 ± 0.08 | 0.22 ± 0.08 |
| | 0.20 ± 0.1 | 0.3 ± 0.1 | 0.22 ± 0.09 | 0.21 ± 0.08 | 0.2 ± 0.1 | 0.2 ± 0.1 | 0.22 ± 0.09 | 0.24 ± 0.08 |
| NDUFA13 | 0.4 ± 0.3 | 0.3 ± 0.2 | 0.2 ± 0.1 | 0.5 ± 0.3 | 0.2 ± 0.2 | 0.3 ± 0.2 | 0.2 ± 0.1 | 0.3 ± 0.2 |
| | 0.3 ± 0.2 | 0.3 ± 0.2 | 0.3 ± 0.2 | 0.4 ± 0.2 | 0.3 ± 0.2 | 0.3 ± 0.1 | 0.3 ± 0.2 | 0.3 ± 0.2 |
| | 0.4 ± 0.2 | 0.3 ± 0.2 | 0.4 ± 0.2 | 0.3 ± 0.2 | 0.3 ± 0.2 | 0.3 ± 0.2 | 0.4 ± 0.2 | 0.3 ± 0.2 |
| NDUFB6 | 0.4 ± 0.2 | 0.3 ± 0.1 | 0.3 ± 0.2 | 0.3 ± 0.1 | 0.3 ± 0.2 | 0.5 ± 0.3 | 0.3 ± 0.2 | 0.4 ± 0.2 |
| | 0.3 ± 0.1 | 0.3 ± 0.1 | 0.3 ± 0.1 | 0.4 ± 0.2 | 0.4 ± 0.2 | 0.3 ± 0.2 | 0.3 ± 0.1 | 0.3 ± 0.1 |
| | 0.3 ± 0.1 | 0.3 ± 0.1 | 0.3 ± 0.1 | 0.3 ± 0.2 | 0.3 ± 0.1 | 0.4 ± 0.1 | 0.3 ± 0.2 | 0.3 ± 0.1 |
| NDUFB7 | 0.4 ± 0.3 | 0.4 ± 0.3 | 0.4 ± 0.2 | 0.4 ± 0.3 | 0.5 ± 0.3 | 0.4 ± 0.2 | 0.4 ± 0.3 | 0.4 ± 0.2 |
| | 0.4 ± 0.3 | 0.4 ± 0.3 | 0.4 ± 0.3 | 0.4 ± 0.3 | 0.4 ± 0.3 | 0.4 ± 0.3 | 0.4 ± 0.3 | 0.4 ± 0.2 |
| | 0.4 ± 0.3 | 0.4 ± 0.3 | 0.5 ± 0.3 | 0.4 ± 0.3 | 0.4 ± 0.2 | 0.4 ± 0.2 | 0.4 ± 0.3 | 0.5 ± 0.2 |
| NDUFB9 | 0.2 ± 0.1 | 0.2 ± 0.1 | 0.2 ± 0.1 | 0.3 ± 0.1 | 0.22 ± 0.08 | 0.2 ± 0.1 | 0.2 ± 0.1 | 0.21 ± 0.09 |
| | 0.18 ± 0.08 | 0.20 ± 0.09 | 0.20 ± 0.09 | 0.18 ± 0.08 | 0.26 ± 0.09 | 0.2 ± 0.1 | 0.2 ± 0.1 | 0.2 ± 0.1 |
| | 0.2 ± 0.1 | 0.2 ± 0.1 | 0.2 ± 0.1 | 0.2 ± 0.1 | 0.2 ± 0.1 | 0.2 ± 0.1 | 0.2 ± 0.1 | 0.18 ± 0.07 |
| NDUFB2 | 0.4 ± 0.2 | 0.3 ± 0.2 | 0.3 ± 0.2 | 0.2 ± 0.1 | 0.3 ± 0.2 | 0.5 ± 0.2 | 0.3 ± 0.1 | 0.4 ± 0.2 |
| | 0.4 ± 0.2 | 0.3 ± 0.2 | 0.3 ± 0.1 | 0.3 ± 0.2 | 0.4 ± 0.2 | 0.4 ± 0.2 | 0.4 ± 0.2 | 0.4 ± 0.2 |
| | 0.3 ± 0.2 | 0.3 ± 0.1 | 0.3 ± 0.1 | 0.4 ± 0.2 | 0.3 ± 0.2 | 0.3 ± 0.1 | 0.3 ± 0.1 | 0.3 ± 0.1 |

| | | | | | | | | |
|---|---|---|---|---|---|---|---|---|
| NDUFB8 | 0.4 ± 0.2 | 0.3 ± 0.1 | 0.3 ± 0.1 | 0.3 ± 0.1 | 0.3 ± 0.1 | 0.3 ± 0.2 | 0.3 ± 0.2 | 0.3 ± 0.1 |
| | 0.3 ± 0.1 | 0.3 ± 0.1 | 0.3 ± 0.2 | 0.3 ± 0.1 | 0.3 ± 0.1 | 0.3 ± 0.1 | 0.21 ± 0.09 | 0.2 ± 0.1 |
| | 0.3 ± 0.2 | 0.3 ± 0.2 | 0.3 ± 0.1 | 0.3 ± 0.2 | 0.34 ± 0.3 | 0.3 ± 0.1 | 0.3 ± 0.2 | 0.3 ± 0.1 |
| ESSS / NDUFB11 | 0.3 ± 0.2 | 0.3 ± 0.2 | 0.3 ± 0.2 | 0.3 ± 0.2 | 0.4 ± 0.2 | 0.4 ± 0.2 | 0.4 ± 0.2 | 0.4 ± 0.2 |
| | 0.4 ± 0.2 | 0.4 ± 0.2 | 0.4 ± 0.2 | 0.3 ± 0.2 | 0.4 ± 0.2 | 0.4 ± 0.2 | 0.4 ± 0.2 | 0.4 ± 0.2 |
| | 0.3 ± 0.1 | 0.3 ± 0.2 | 0.3 ± 0.2 | 0.4 ± 0.2 | 0.3 ± 0.2 | 0.4 ± 0.2 | 0.3 ± 0.1 | 0.3 ± 0.2 |
| KFYI / NDUFC1 | 0.2 ± 0.1 | 0.2 ± 0.1 | 0.2 ± 0.1 | 0.3 ± 0.1 | 0.2 ± 0.1 | 0.14 ± 0.07 | 0.2 ± 0.1 | 0.3 ± 0.2 |
| | 0.17 ± 0.08 | 0.2 ± 0.1 | 0.2 ± 0.1 | 0.2 ± 0.1 | 0.2 ± 0.1 | 0.2 ± 0.2 | 0.2 ± 0.1 | 0.2 ± 0.1 |
| | 0.2 ± 0.1 | 0.2 ± 0.1 | 0.2 ± 0.1 | 0.18 ± 0.08 | 0.17 ± 0.09 | 0.2 ± 0.1 | 0.2 ± 0.1 | 0.3 ± 0.2 |
| MNLL/ NDUFB1 | 0.3 ± 0.1 | 0.3 ± 0.1 | 0.3 ± 0.1 | 0.3 ± 0.1 | 0.4 ± 0.2 | 0.4 ± 0.1 | 0.4 ± 0.2 | 0.3 ± 0.09 |
| | 0.3 ± 0.1 | 0.3 ± 0.1 | 0.3 ± 0.1 | 0.3 ± 0.2 | 0.3 ± 0.2 | 0.5 ± 0.2 | 0.3 ± 0.1 | 0.4 ± 0.1 |
| | 0.3 ± 0.1 | 0.3 ± 0.1 | 0.3 ± 0.2 | 0.4 ± 0.1 | 0.4 ± 0.1 | 0.4 ± 0.1 | 0.3 ± 0.1 | 0.3 ± 0.1 |
| MWFE / NDUFA1 | 0.2 ± 0.1 | 0.12 ± 0.08 | 0.2 ± 0.1 | 0.3 ± 0.2 | 0.2 ± 0.1 | 0.2 ± 0.2 | 0.3 ± 0.2 | 0.2 ± 0.2 |
| | 0.2 ± 0.1 | 0.3 ± 0.1 | 0.3 ± 0.2 | 0.2 ± 0.1 | 0.2 ± 0.1 | 0.2 ± 0.1 | 0.2 ± 0.1 | 0.3 ± 0.2 |
| | 0.2 ± 0.1 | 0.14 ± 0.07 | 0.3 ± 0.2 | 0.21 ± 0.09 | 0.3 ± 0.1 | 0.2 ± 0.1 | 0.2 ± 0.1 | 0.3 ± 0.2 |

The NDUFS2 (49 kDa in the ovine naming) subunit showed high average RMSF values (0.3 nm). As discussed above, these fluctuations are probably caused by the hydrophobic residues on the surface of this subunit, which are exposed to the solvent. The NDUFA11 and NDUFB5 subunits show the lower average RMSF values than the other supernumerary subunits (below 0.2 nm). The NDUFA3, NDUFA13, NDUFB6, NDUFB7, NDUFB2, NDUFB11 (ESSS) and NDUFB1 subunits have usually the highest average RMSF values (over 0.3 nm), which are caused by the large fluctuations of the N- and C-terminals.

**Fig. S18. Lipid density maps calculated on the summed trajectories of the wtH$^{op}$ system.** Cardiolipin, POPC, POPE and POPI maps are in blue, orange, light blue and green, respectively. The section of P+ module is reported, and the subunits are colored according to Fig. 1B scheme.

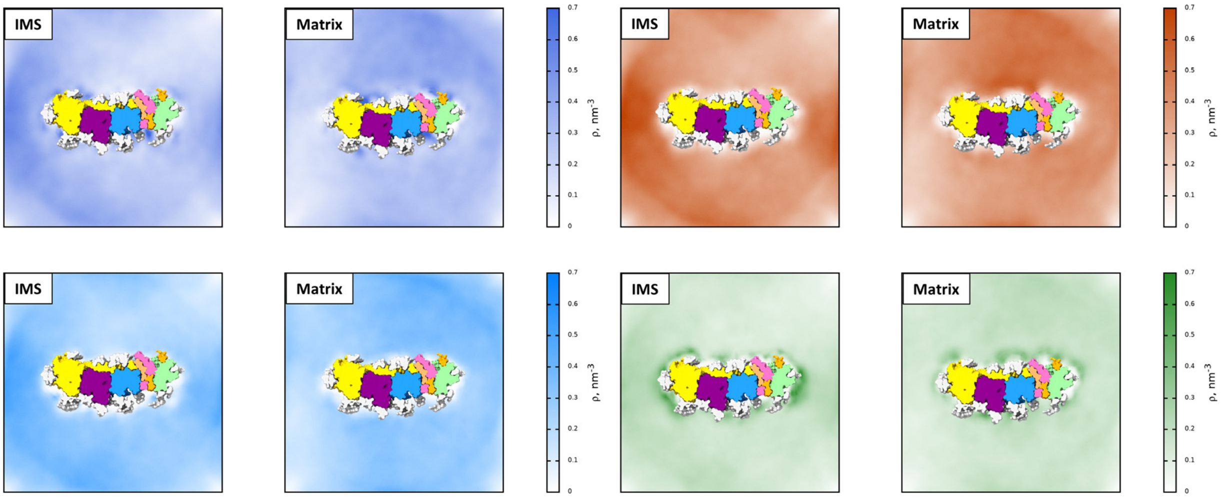


**Fig. S19. Lipid density maps calculated on the summed trajectories of the 1mH$^{op}$ system.** Cardiolipin, POPC, POPE and POPI maps are in blue, orange, light blue and green, respectively. The section of P+ module is reported, and the subunits are colored according to Fig. 1B scheme.

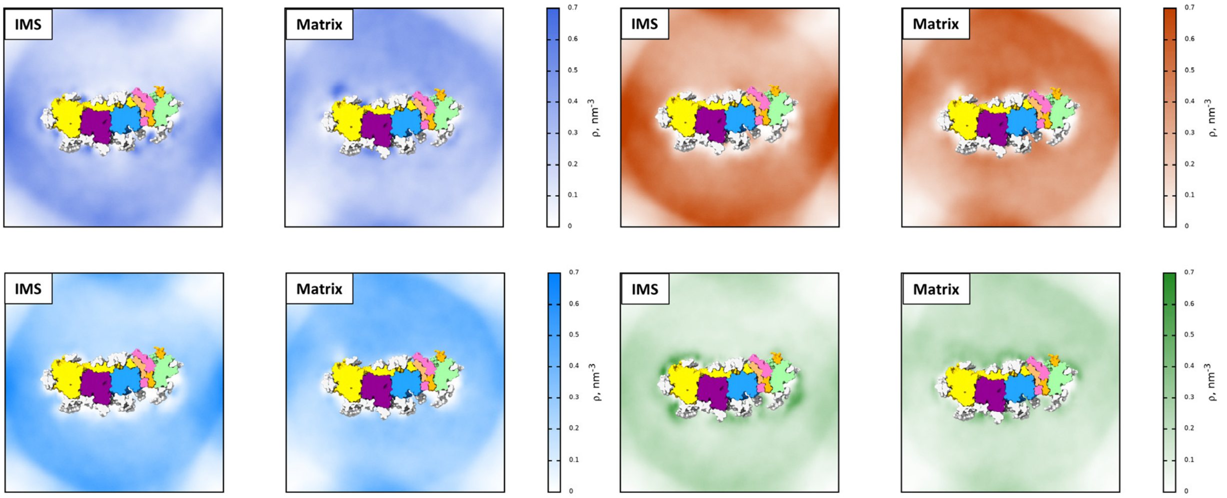

**Fig. S20. Maps of lipid density on the summed trajectories of the 1mH$^{op}$ system.** Cardiolipin, POPC, POPE and POPI maps are in blue, orange, light blue and green, respectively. The section of P+ module is reported, and the subunits are colored according to Fig. 1B scheme.

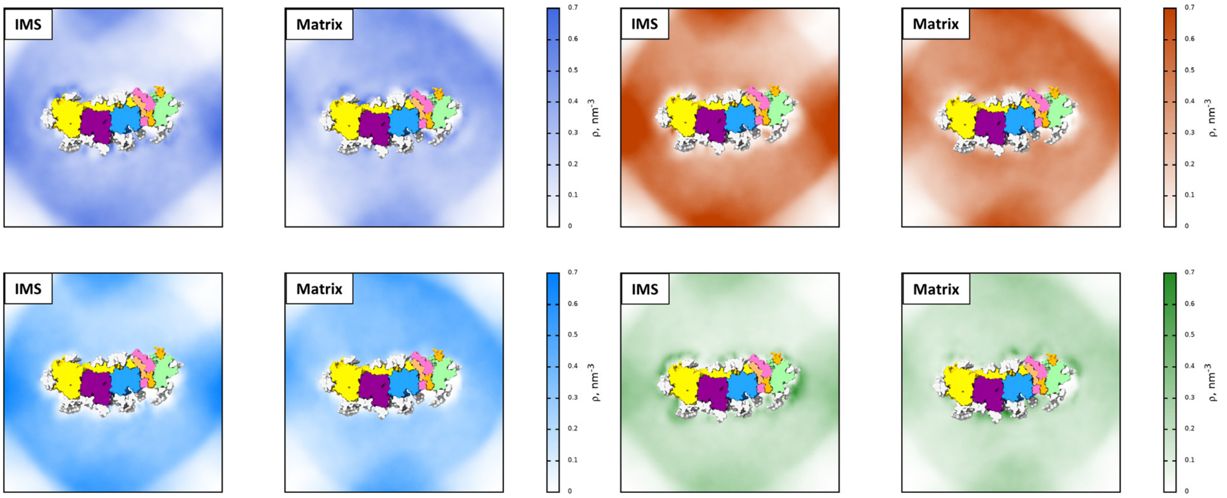


**Fig. S21. Maps of lipid density on the summed trajectories of the wtH$^{cl}$ system.** Cardiolipin, POPC, POPE and POPI maps are in blue, orange, light blue and green, respectively. The section of P+ module is reported, and the subunits are colored according to Fig. 1B scheme.

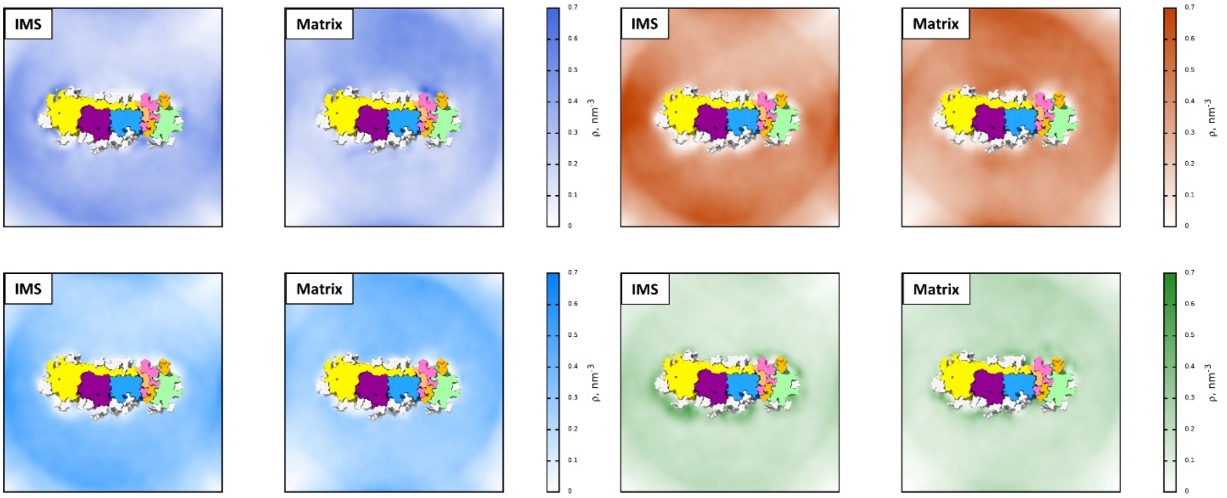

**Fig. S22. Maps of lipid density on the summed trajectories of the 1mH$^{cl}$ system.** Cardiolipin, POPC, POPE and POPI maps are in blue, orange, light blue and green, respectively. The section of P+ module is reported, and the subunits are colored according to Fig. 1B scheme.

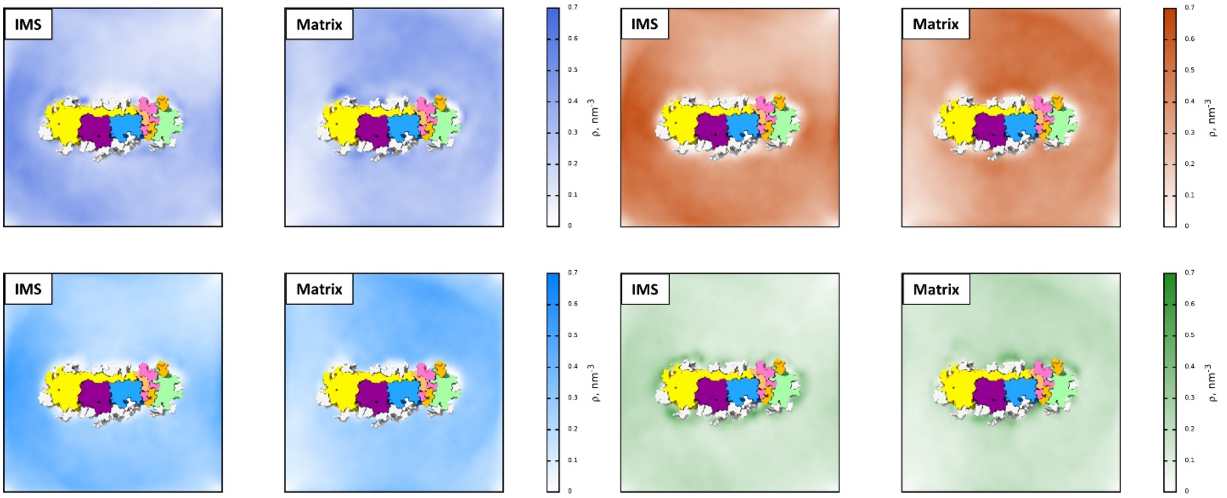


**Fig. S23. Maps of lipid density on the summed trajectories of the 3mH$^{cl}$ system.** Cardiolipin, POPC, POPE and POPI maps are in blue, orange, light blue and green, respectively. The section of P+ module is reported, and the subunits are colored according to Fig. 1B scheme.

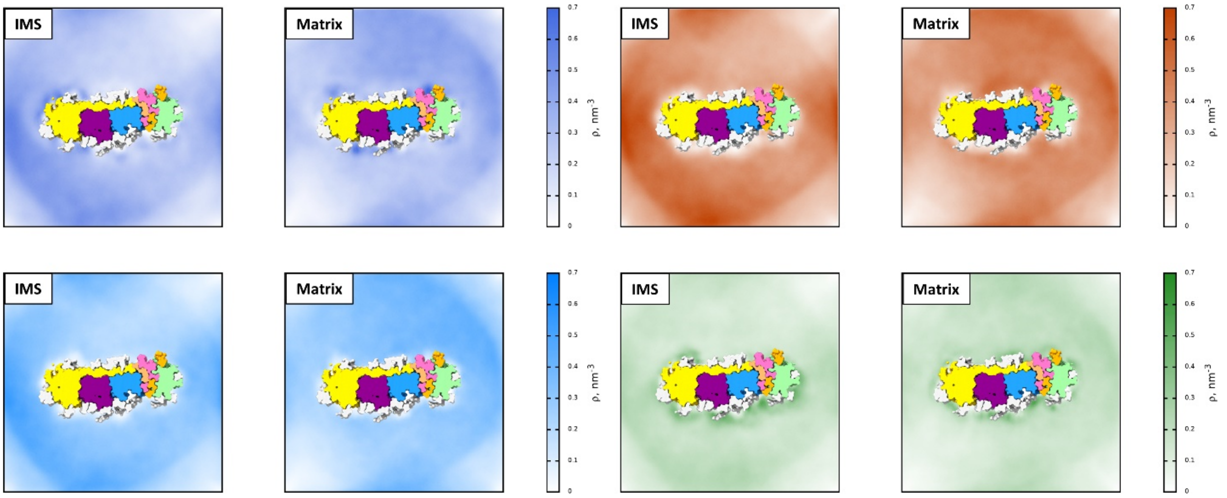

**Fig. S24. Maps of lipid density on the summed trajectories of the wtO$^{op}$ system.** Cardiolipin, POPC, POPE and POPI maps are in blue, orange, light blue and green, respectively. The section of P+ module is reported, and the subunits are colored according to Fig. 1B scheme.

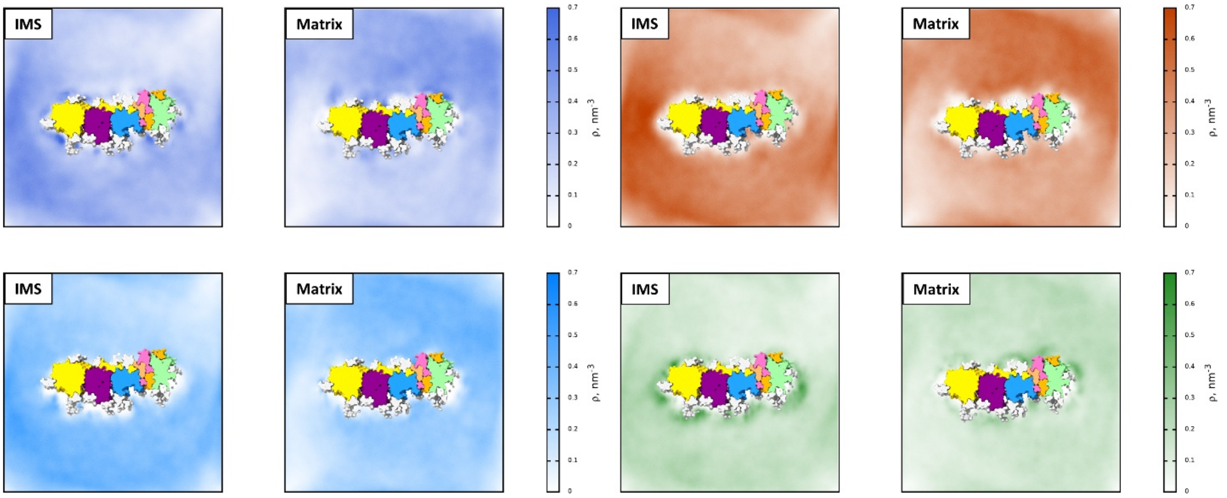


**Fig. S25. Maps of lipid density on the summed trajectories of the wtO$^{cl}$ system.** Cardiolipin, POPC, POPE and POPI maps are in blue, orange, light blue and green, respectively. The section of P+ module is reported, and the subunits are colored according to Fig. 1B scheme.

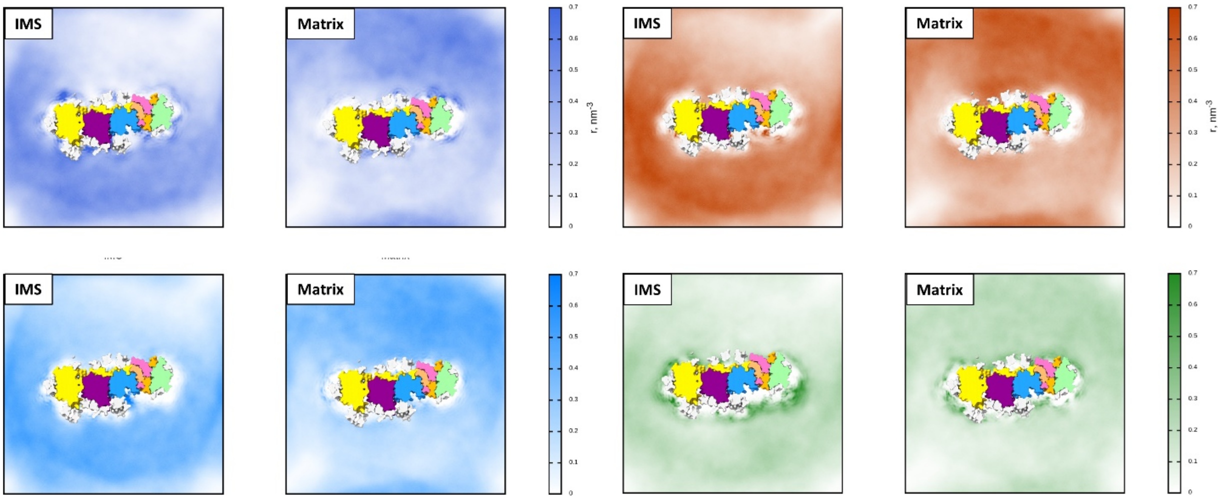

**Fig. S26. Average RMSF *vs.* residue number plots of the ND1 (top panels) and ND3 (bottom panels) subunits in the human systems.**

Left and right panels report the results for the close and open system, respectively. The average RMSF of the *wild type* subunits are in red, while the single and the triple mutants are in blue and green, respectively. The width of the shading shows the standard deviation calculated on the three replicas of each simulation. The position of the TMHs is reported above the plots.

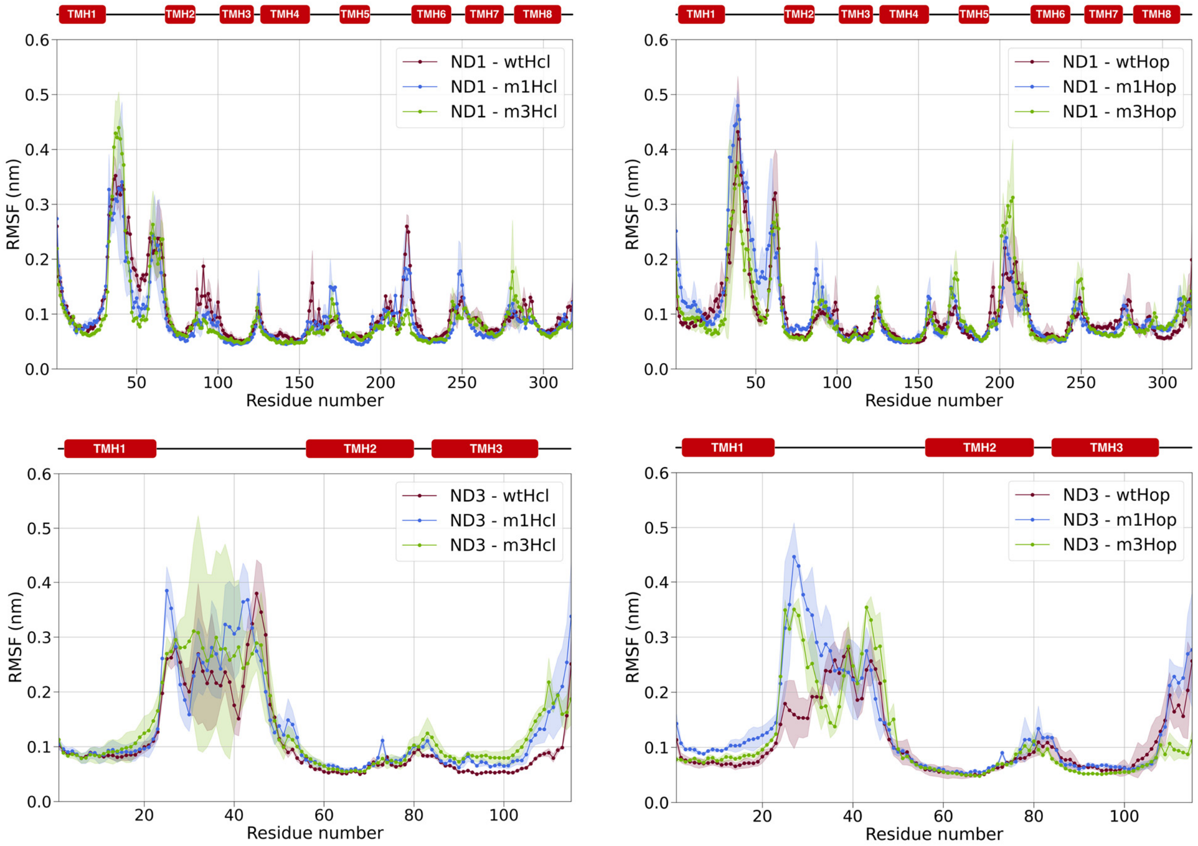

**Fig. S27. Average RMSF *vs.* residue number plots of the ND2 (top panels) and ND4 (bottom panels) subunits in the human systems.**

Left and right panels report the results for the close and open system, respectively. The average RMSF of the *wild type* subunits are in red, while the single and the triple mutants are in blue and green, respectively. The width of the shading shows the standard deviation calculated on the three replicas of each simulation. The position of the TMHs is reported above the plots.

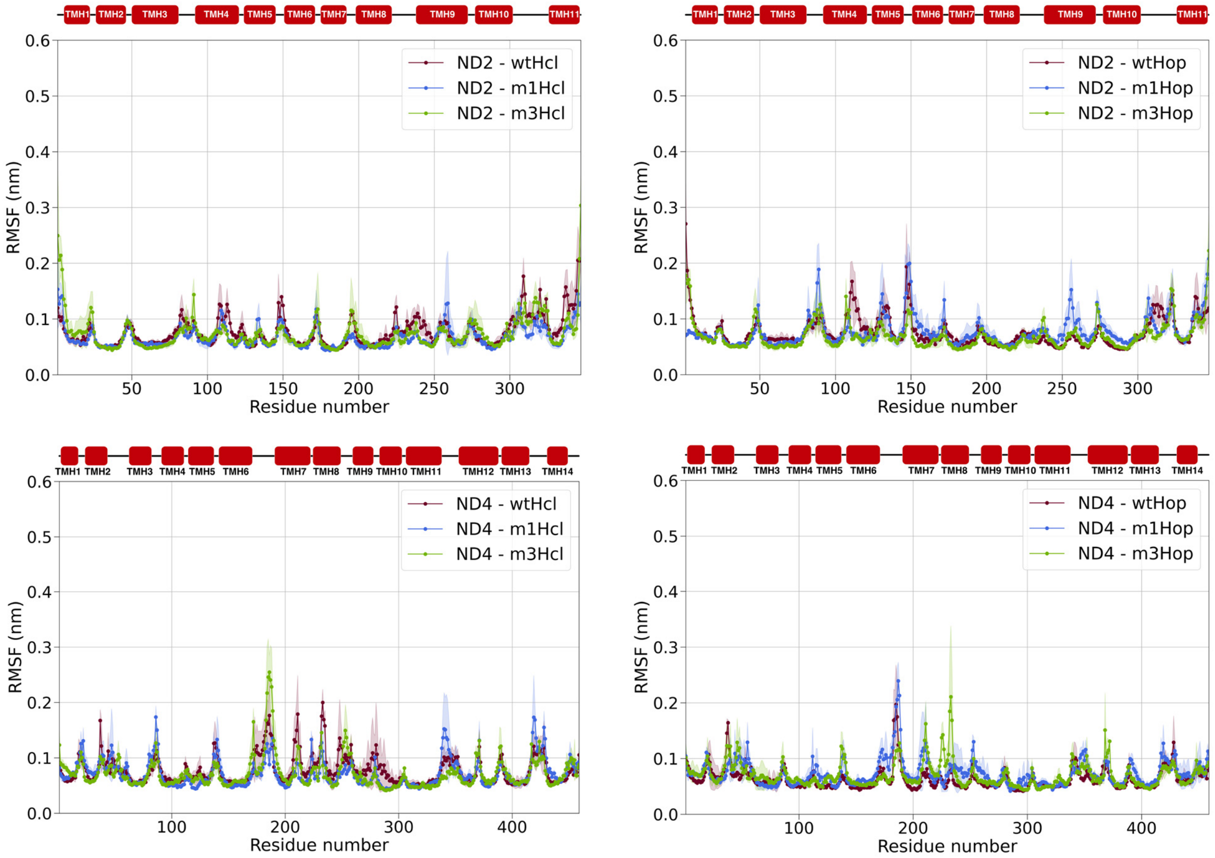

**Fig. S28. Average root mean square fluctuations values as a function of the residue number for ND4L in the human close and open conformation, respectively.**

The *wild type* ND4L is in red, while the single and the triple mutants are in blue and green, respectively. The width of the shading shows the standard deviation calculated on the three replicas of each simulation. The mutation position is indicated by a black arrow, while the secondary structure is reported above the plots. The right panels report a close up of the average RMSF in the region around the p.A71T/ND4L variant, which includes the TMH3 and the nearby loops, for the human close and open systems, respectively.

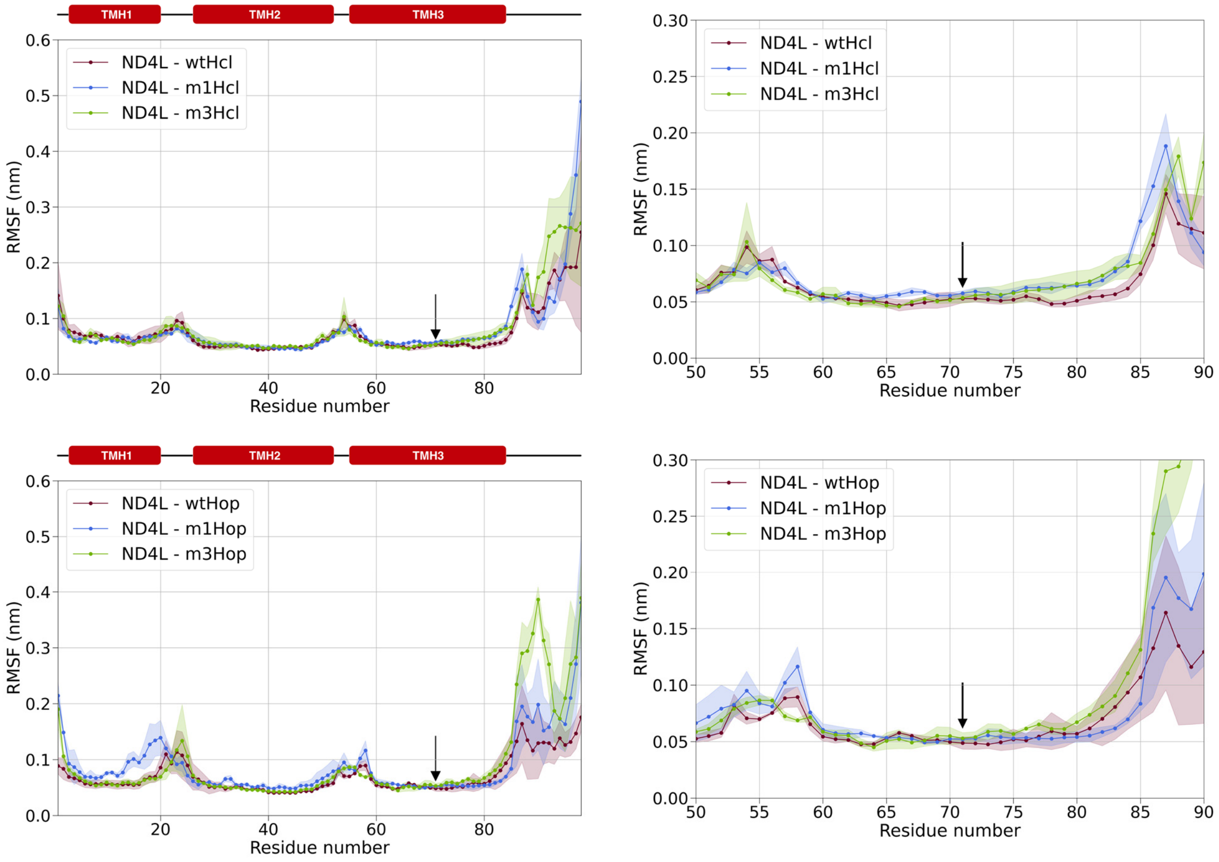

**Fig. S29. Average root mean square fluctuations values as a function of the residue number for ND5 in the human close and open conformation, respectively.**
The *wild type* ND5 is in red, while the single and the triple mutants are in blue and green, respectively. The width of the shading shows the standard deviation calculated on the three replicas of each simulation. The mutation position is indicated by a black arrow, while the position of the TMHs is reported above the plots. The right panels report a detail of the average RMSF in the region around the p.T536A/ND5 mutation, which includes the lateral helix (HL) and the nearby loops, for the human close and open systems, respectively.

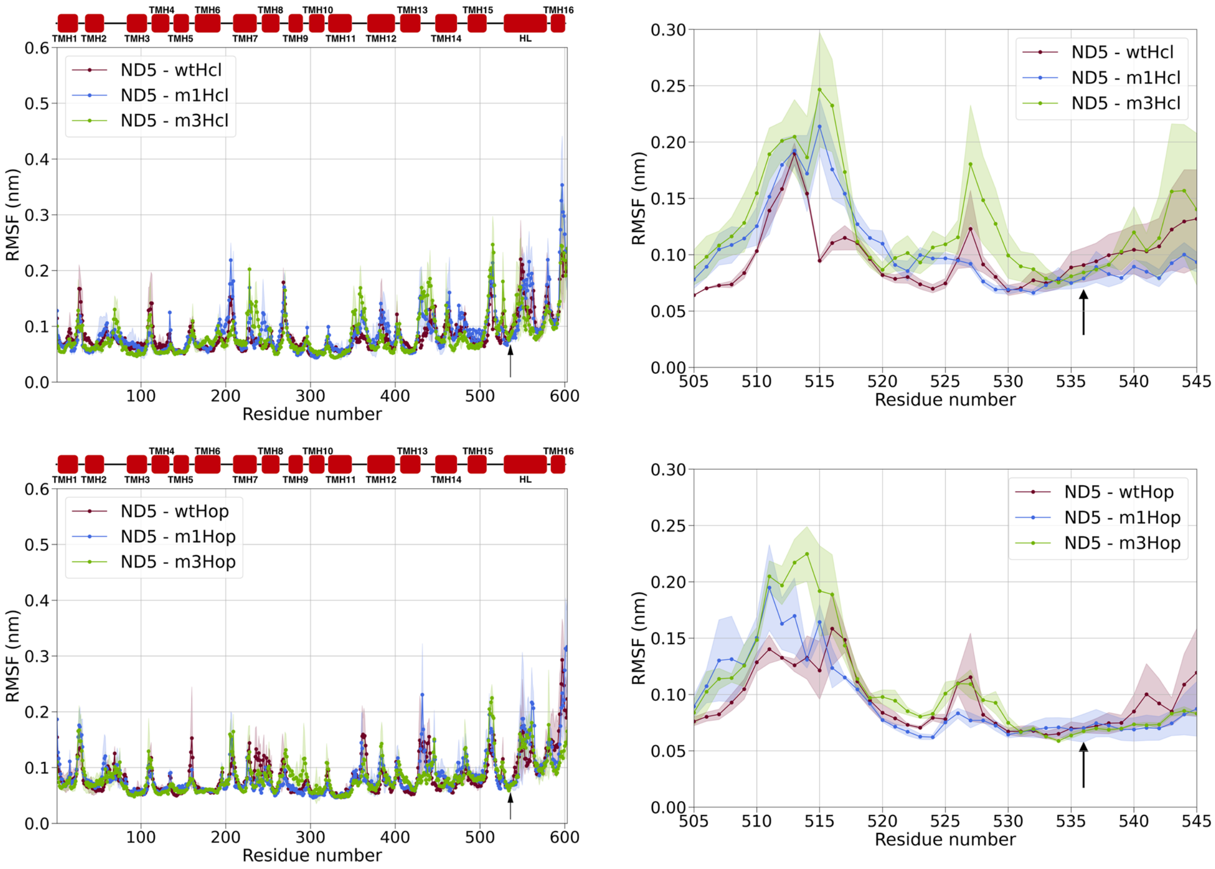

**Fig. S30. Average RMSF *vs.* residue number plots of the ND1 (top panels) and ND3 (bottom panels) subunits in the ovine systems.**

Left and right panels report the results for the close and open system, respectively. The average RMSF of the *wild type* subunits are in red. The width of the shading shows the standard deviation calculated on the three replicas of each simulation. The position of the TMHs is reported above the plots.

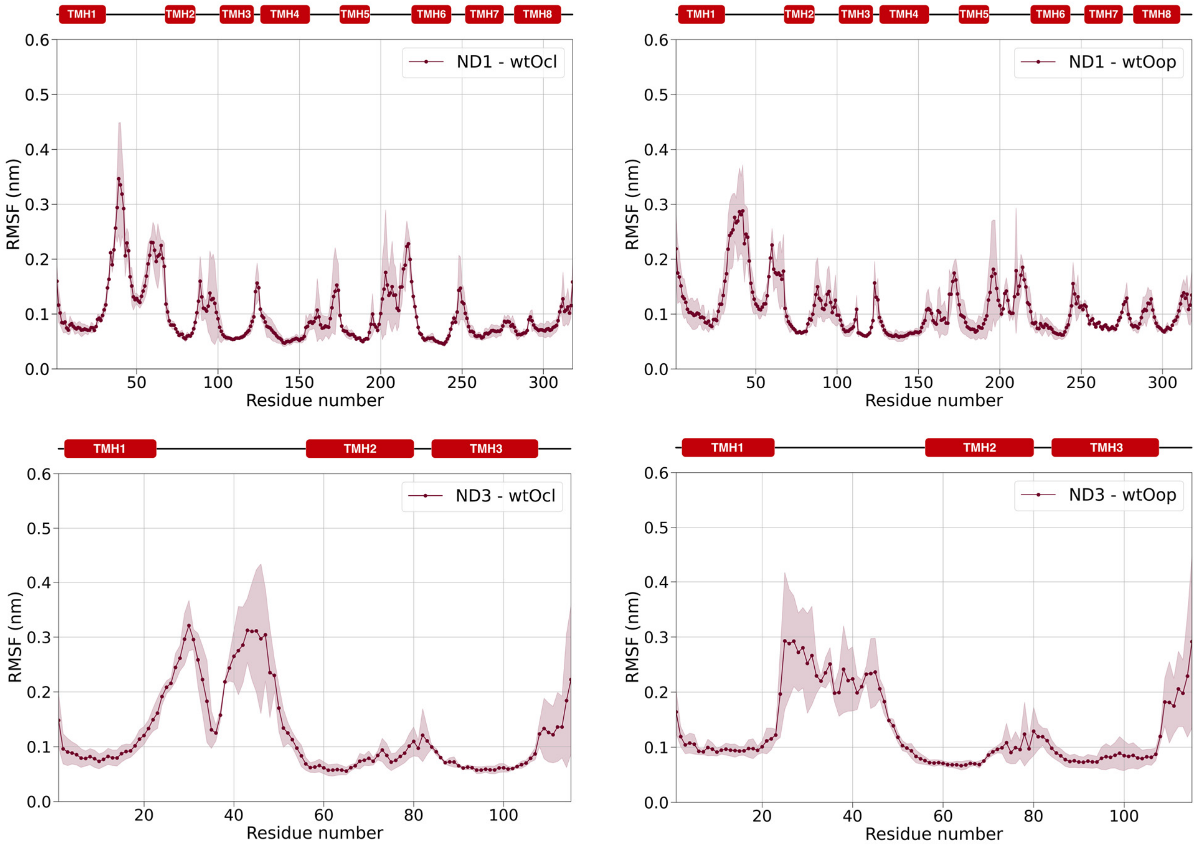

**Fig. S31. Average RMSF *vs.* residue number plots of the ND2 (top panels) and ND4 (bottom panels) subunits in the ovine systems.**

Left and right panels report the results for the close and open system, respectively. The average RMSF of the *wild type* subunits are in red. The width of the shading shows the standard deviation calculated on the three replicas of each simulation. The position of the TMHs is reported above the plots.

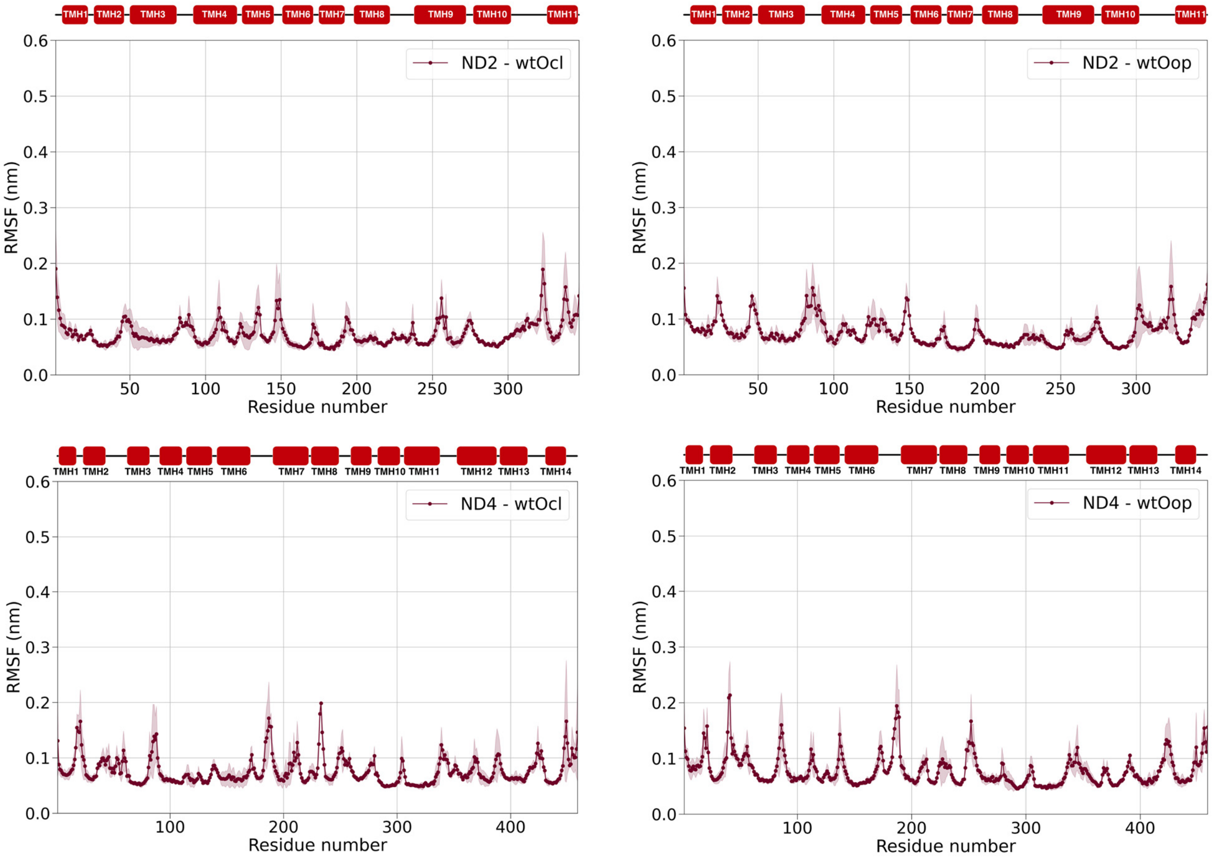

**Fig. S32. Average root mean square fluctuations values as a function of the residue number for the ND6 (top panels), ND4L (middle panels) and ND5 (bottom panels) in the ovine close (left panels) and open (right panels) conformation, respectively.**
The average RMSF of the *wild type* subunits are in red. The width of the shading shows the standard deviation calculated on the three replicas of each simulation. The positions of the mutated residues in the human subunits are indicated by a black arrow, while the position of the TMHs is reported above the plots.

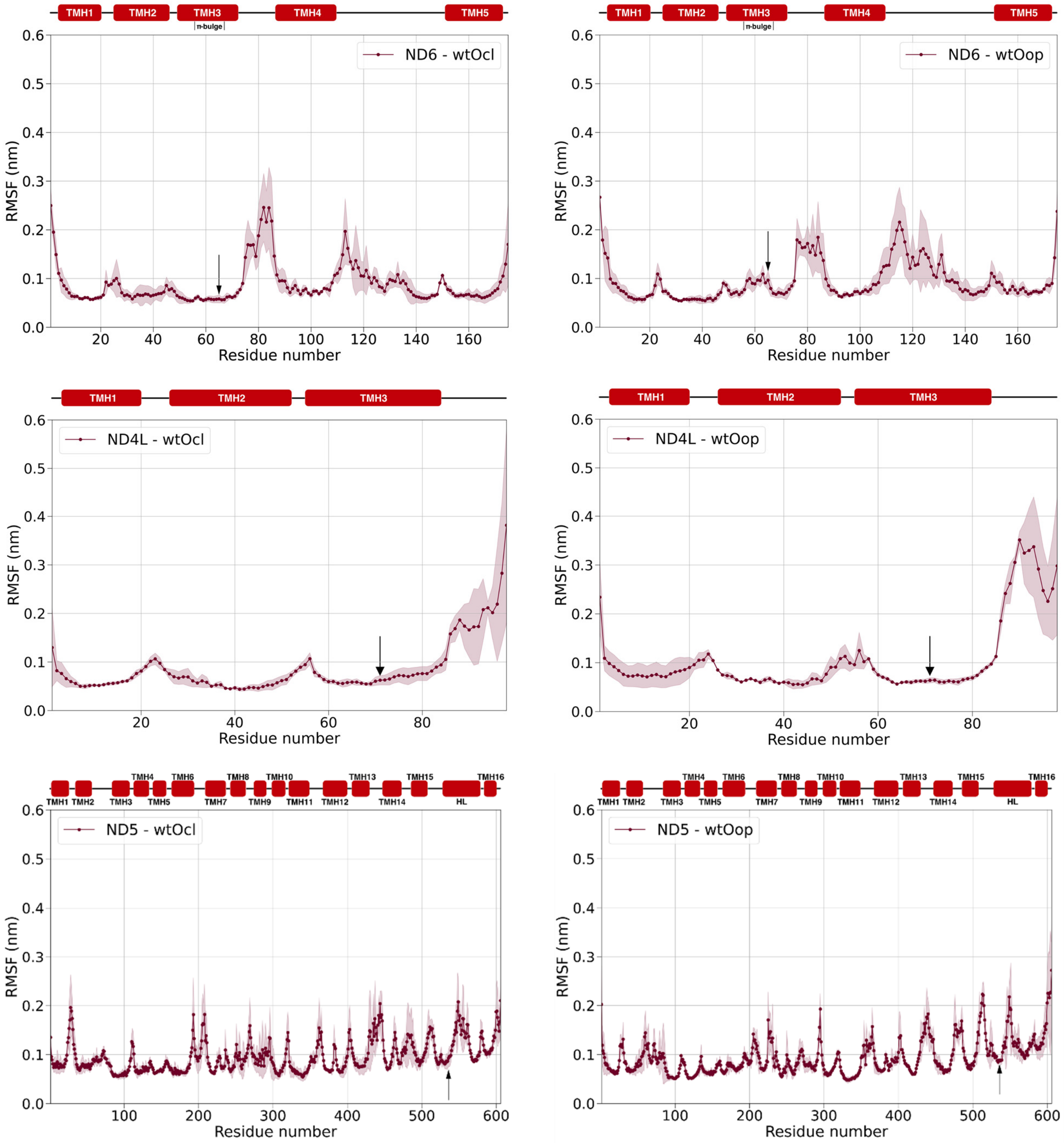

**Fig. S33. Residues within 0.6 nm of p.T536A/ND5.**
Simulations conducted from the *wild type*, single mutant and triple mutant systems are in red, blue and green bars, respectively. The height of the bars is proportional to the fraction of simulation time. The upper and bottom panels refer to the simulations conducted starting from the close and the open conformation, respectively. The six nearest residues to p.T536A/ND5 in the ND5 aminoacidic sequence were excluded from the analysis.

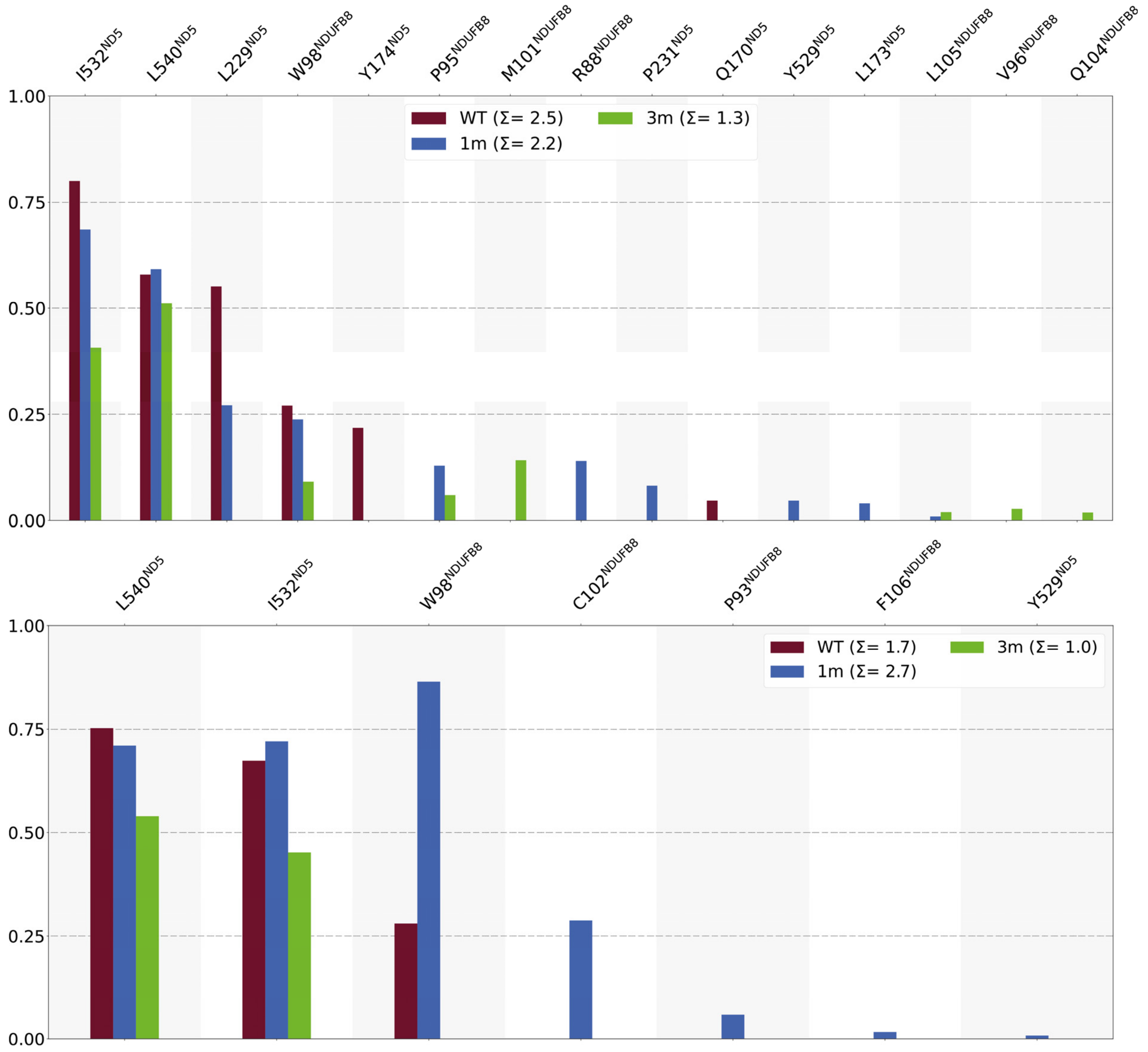

**Fig. S34. Projections along the first three principal modes obtained from the combined essential dynamics analysis performed on the time evolving BB distance matrix.**
Panels **A** and **B** refer to the open and close systems respectively. The *wild type* projections are in red, while the single and the triple mutants are in blue and green, respectively.

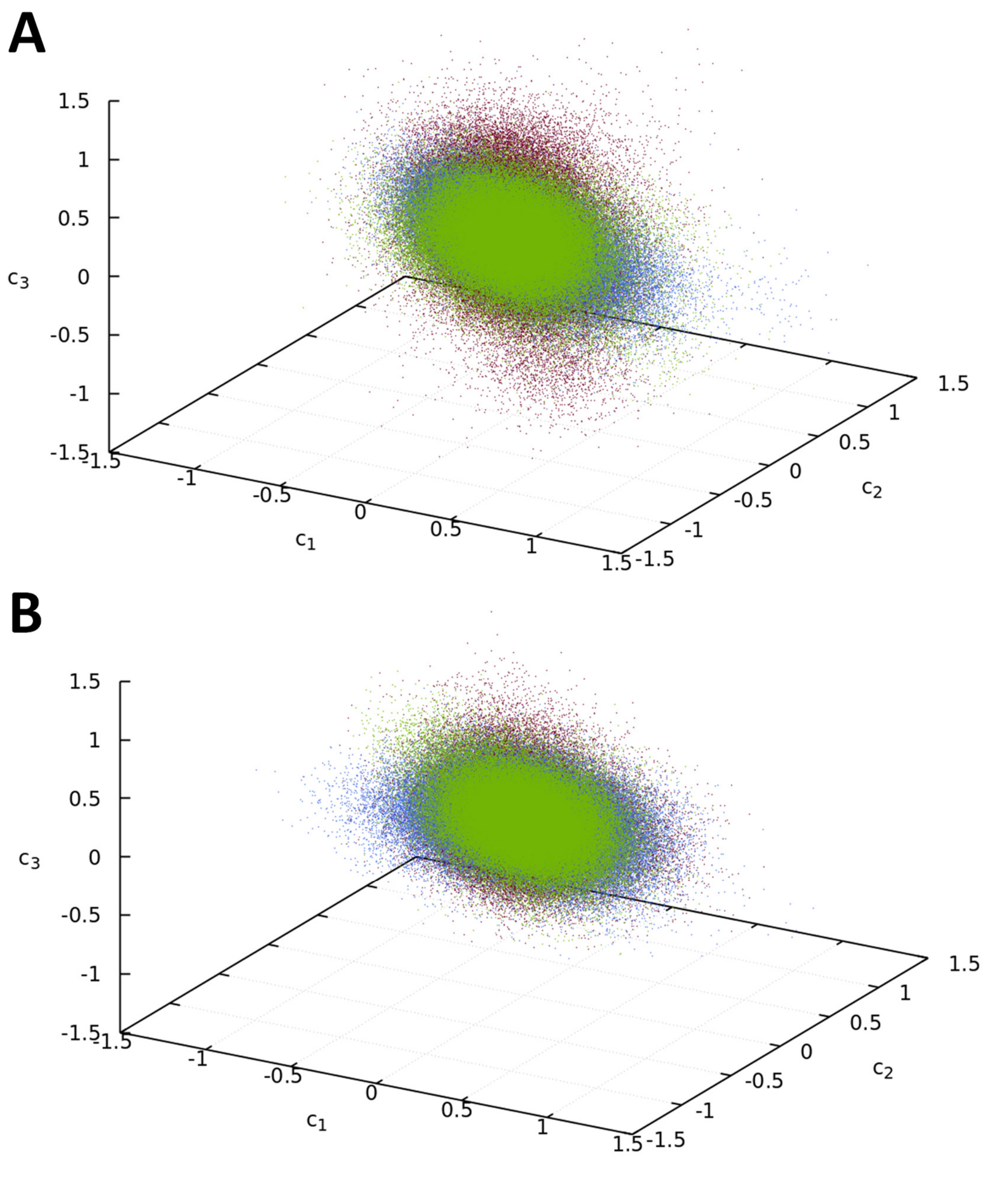

**Fig. S35. Analysis of the BB dihedrals in the TMH3/ND6 π-bulge region.**
Distribution of the backbone dihedral angles observed throughout the simulations in the close systems. The *wild type* is in red, while the single and triple mutants are in blue and green.

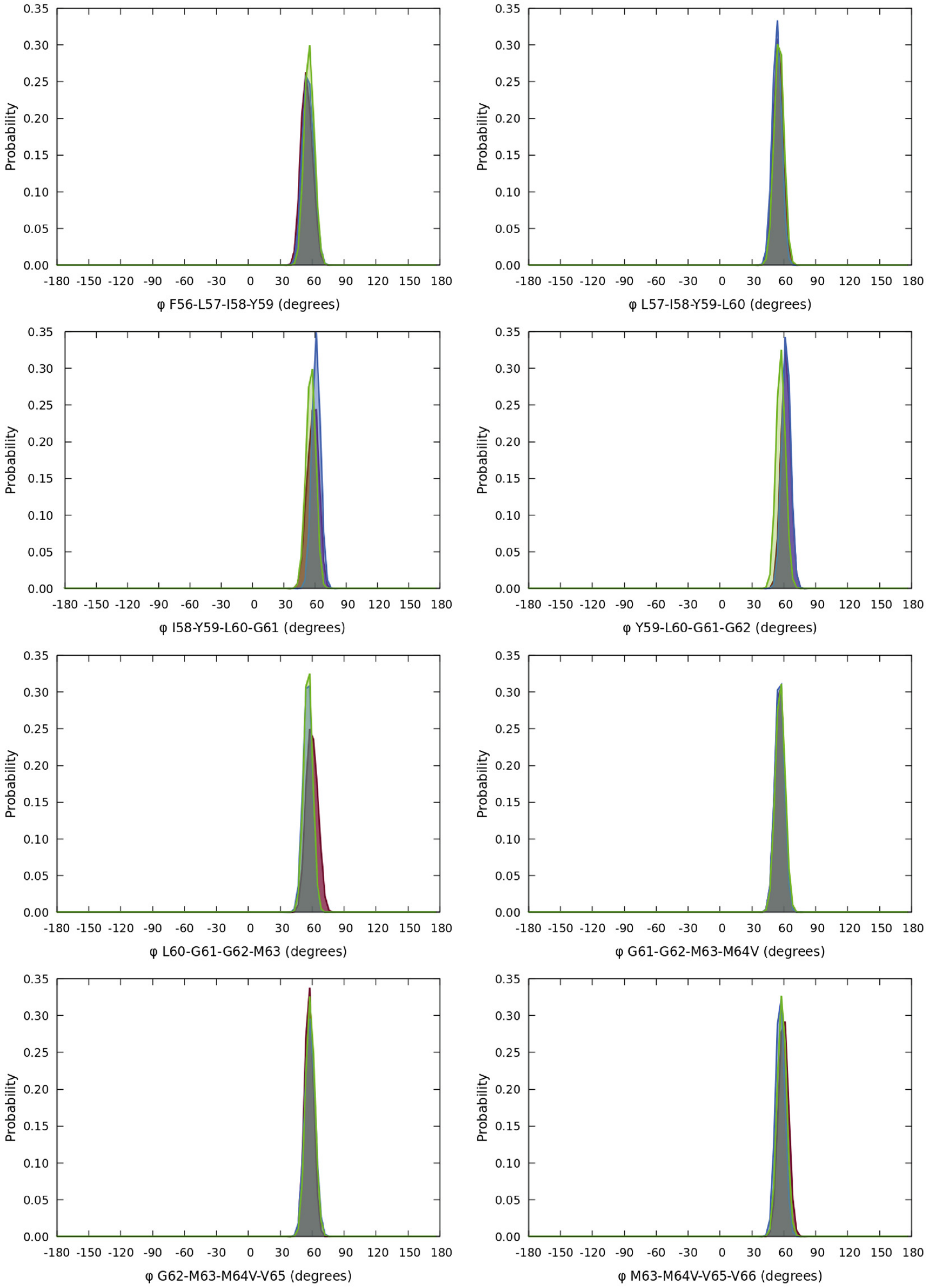

**Fig. S36. Analysis of the BB dihedrals in the TMH3/ND6 π-bulge region.**
Distribution of the backbone dihedral angles observed throughout the simulations in the first replica of wtH$^{op}$ system (brown dots). The Gaussian curves used to fit the dihedral angles distribution and the sum of the fitting Gaussians are in red.

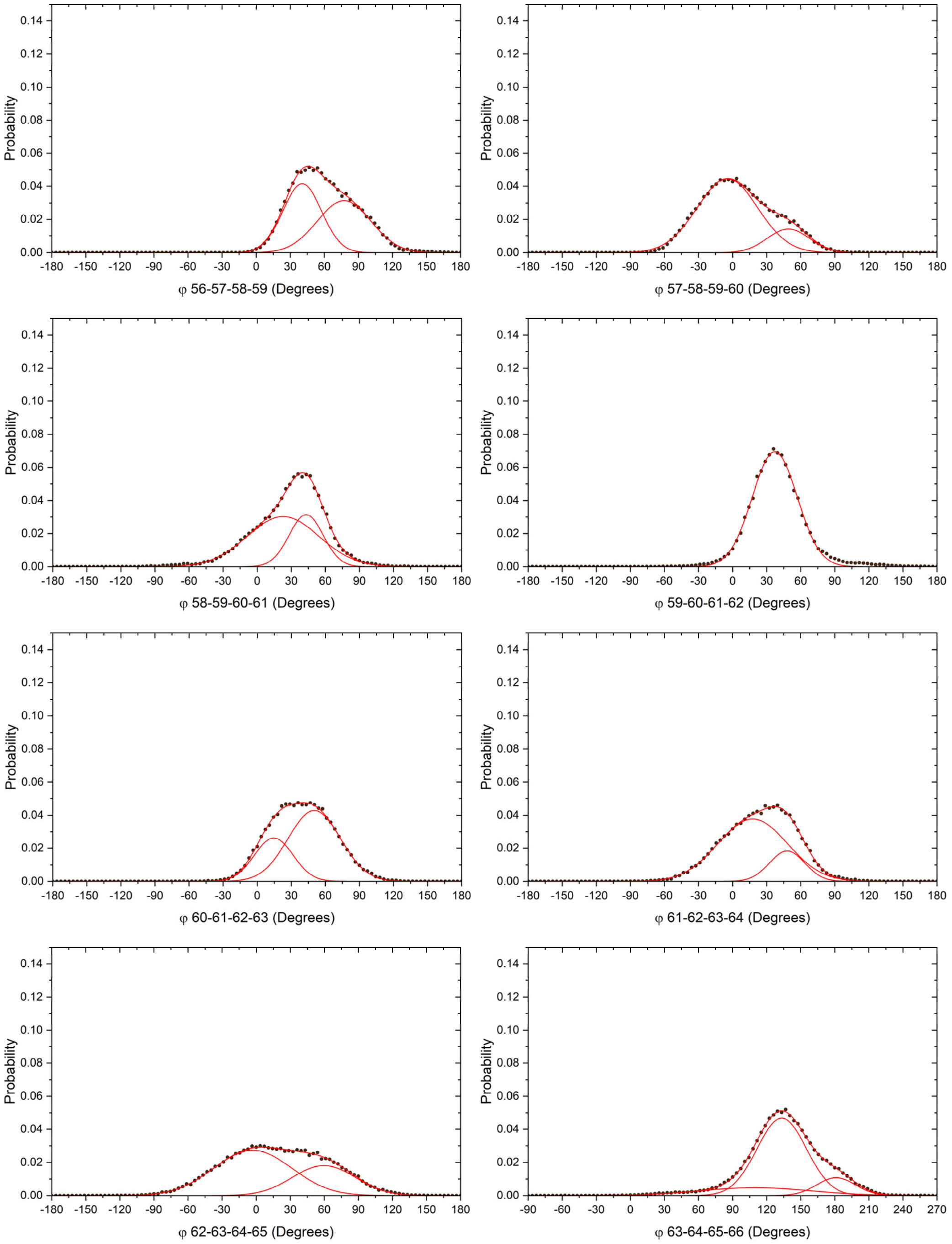

**Fig. S37. Analysis of the BB dihedrals in the TMH3/ND6 π-bulge region.**
Distribution of the backbone dihedral angles observed throughout the simulations in the second replica of wtH$^{op}$ system (maroon dots). The Gaussian curves used to fit the dihedral angles distribution and the sum of the fitting Gaussians are in red.

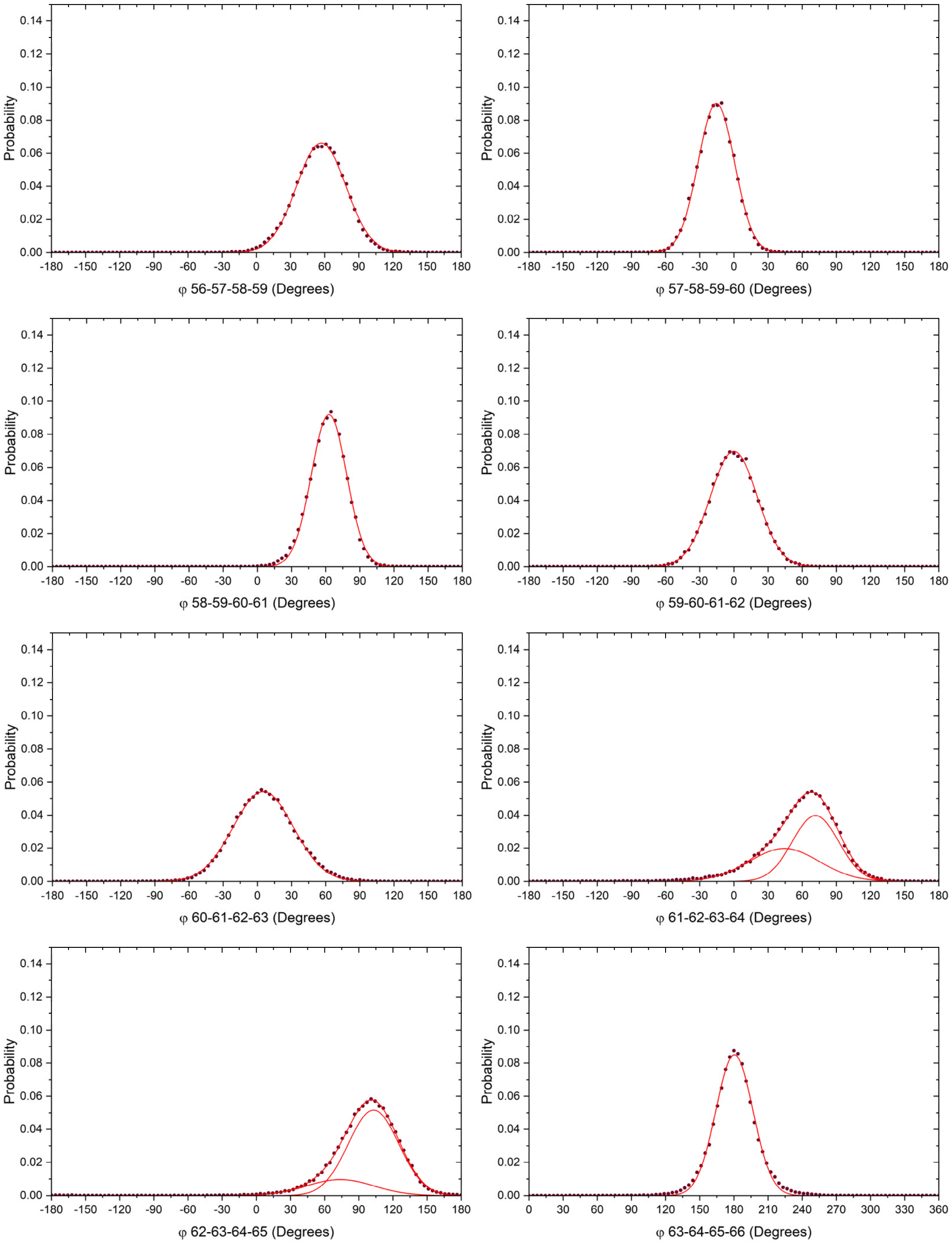

**Fig. S38. Analysis of the BB dihedrals in the TMH3/ND6 π-bulge region.**
Distribution of the backbone dihedral angles observed throughout the simulations in the third replica of wtH$^{op}$ system (crimson dots). The Gaussian curves used to fit the dihedral angles distribution and the sum of the fitting Gaussians are in red.

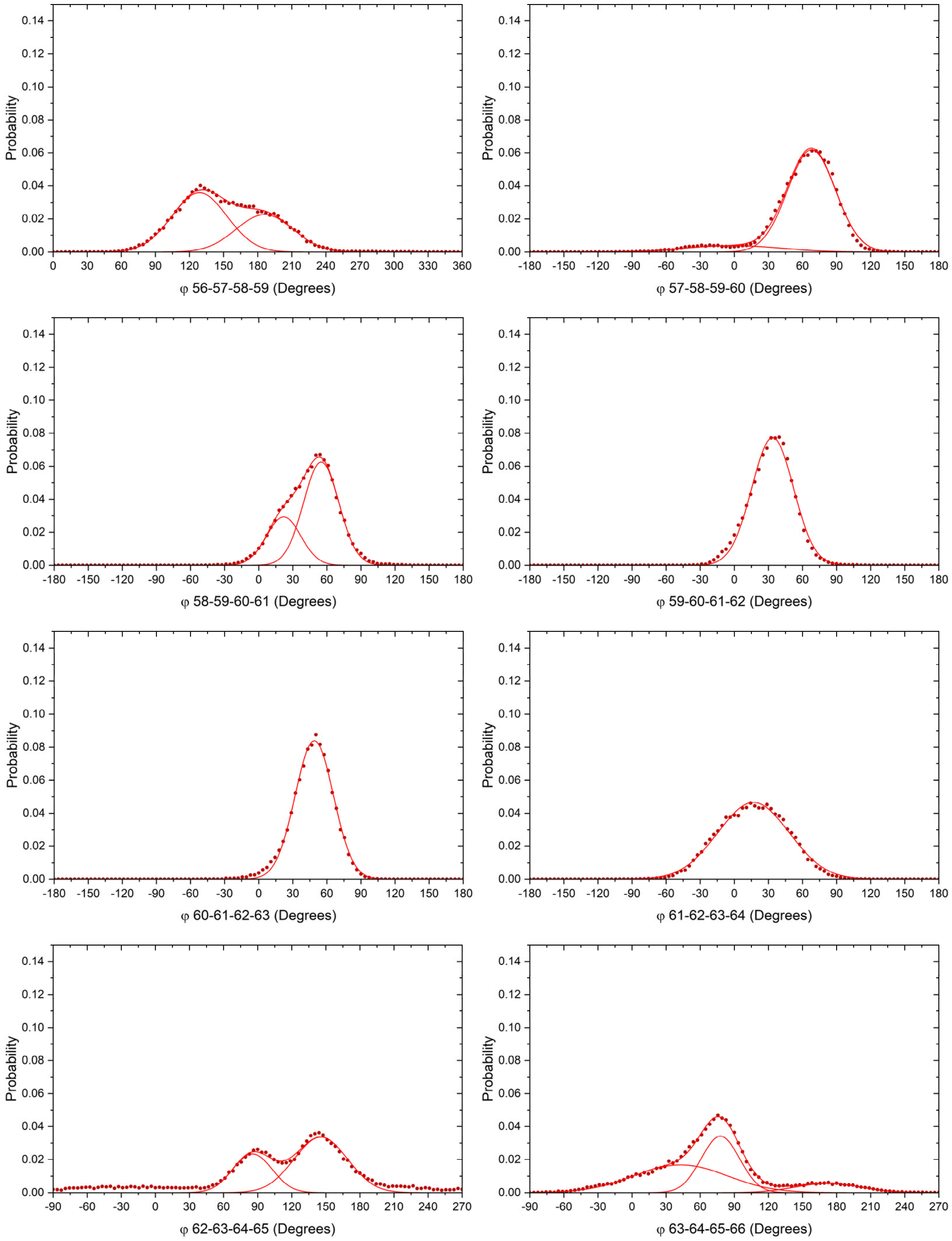

**Fig. S39. Analysis of the BB dihedrals in the TMH3/ND6 π-bulge region.**
Distribution of the backbone dihedral angles observed throughout the simulations in the first replica of m1H$^{op}$ system (navy dots). The Gaussian curves used to fit the dihedral angles distribution and the sum of the fitting Gaussians are in red.

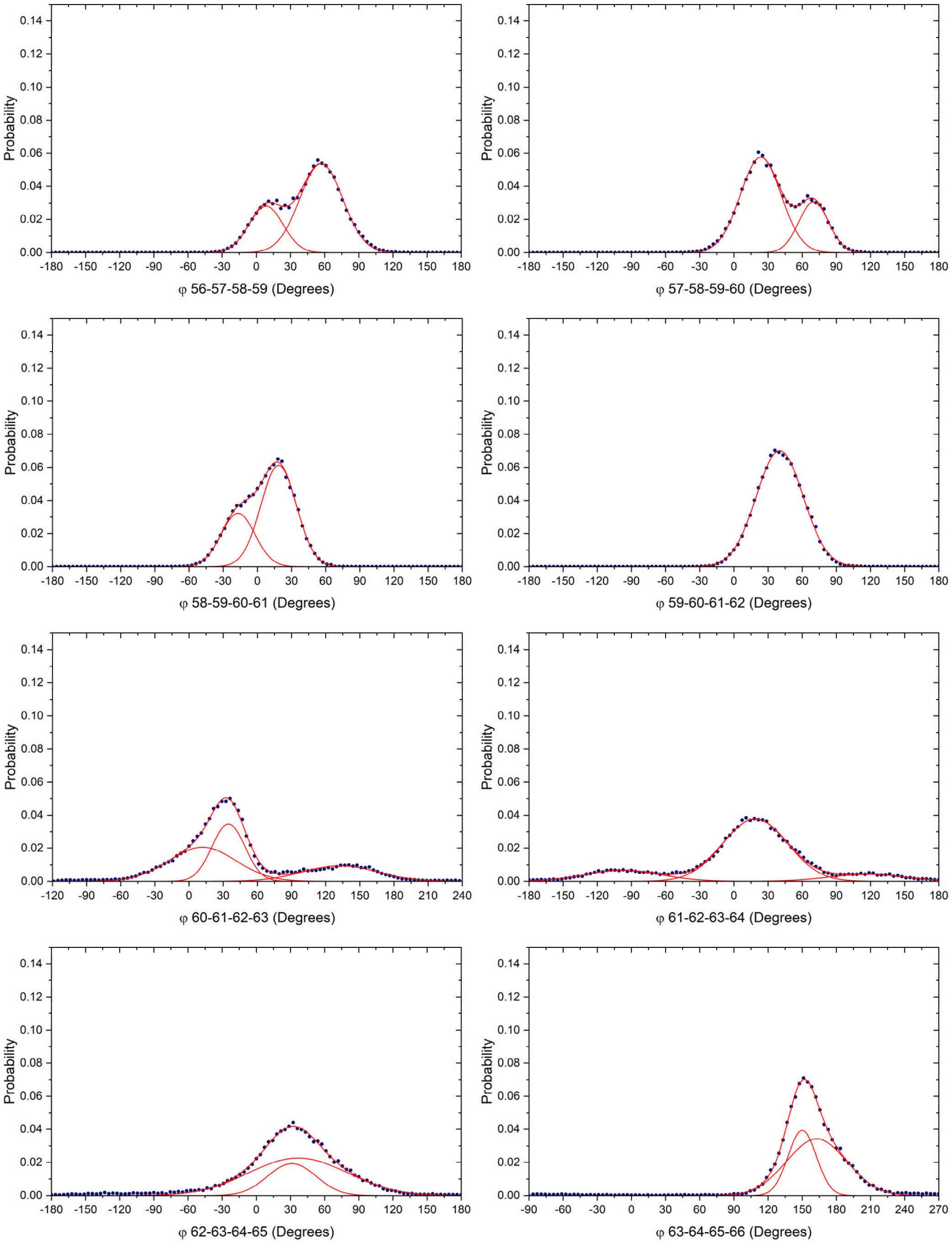

**Fig. S40. Analysis of the BB dihedrals in the TMH3/ND6 π-bulge region.**
Distribution of the backbone dihedral angles observed throughout the simulations in the second replica of m1H$^{op}$ system (royal blue dots). The Gaussian curves used to fit the dihedral angles distribution and the sum of the fitting Gaussians are in red.

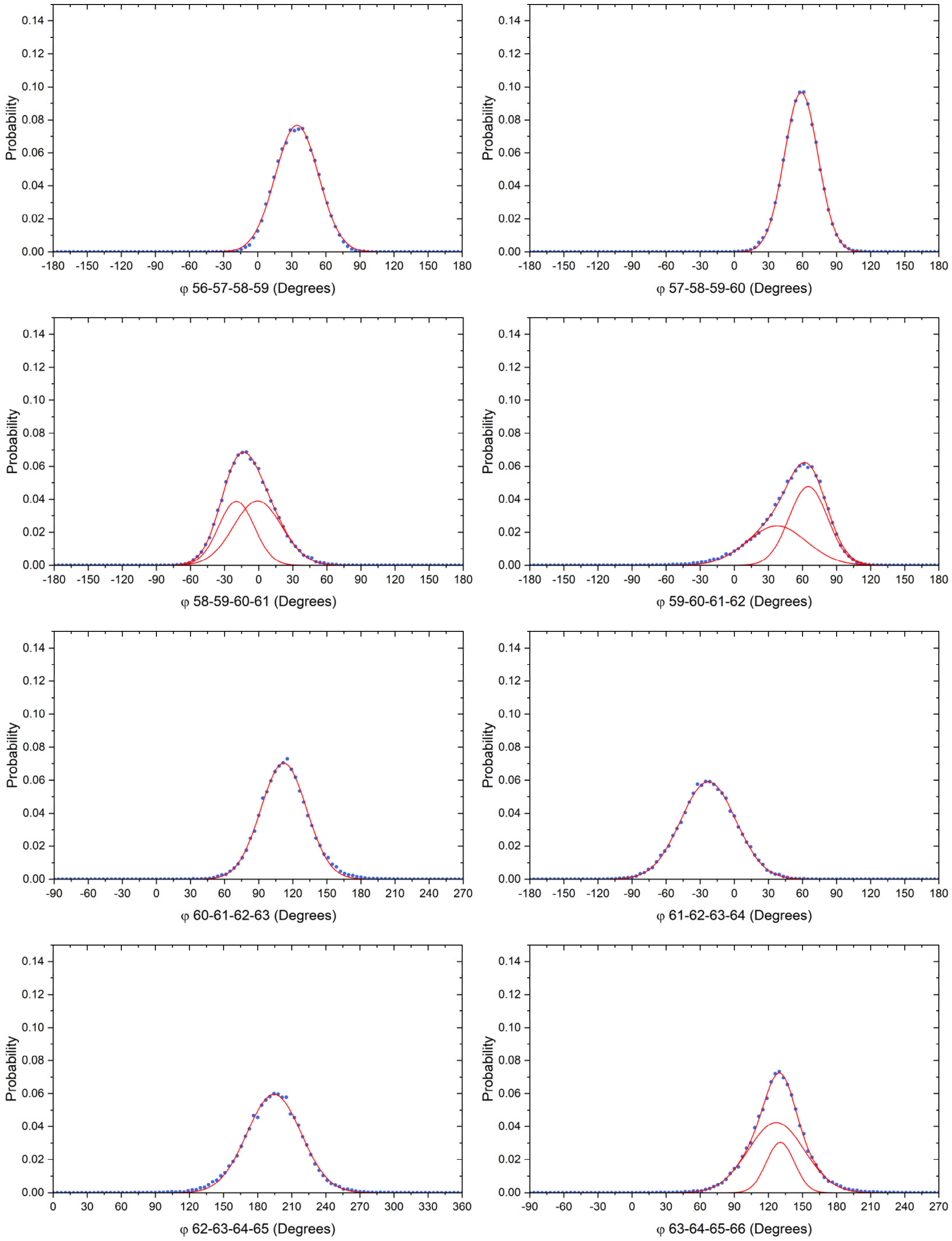

**Fig. S41. Analysis of the BB dihedrals in the TMH3/ND6 π-bulge region.**
Distribution of the backbone dihedral angles observed throughout the simulations in the third replica of m1H$^{op}$ system (fountain blue dots). The Gaussian curves used to fit the dihedral angles distribution and the sum of the fitting Gaussians are in red.

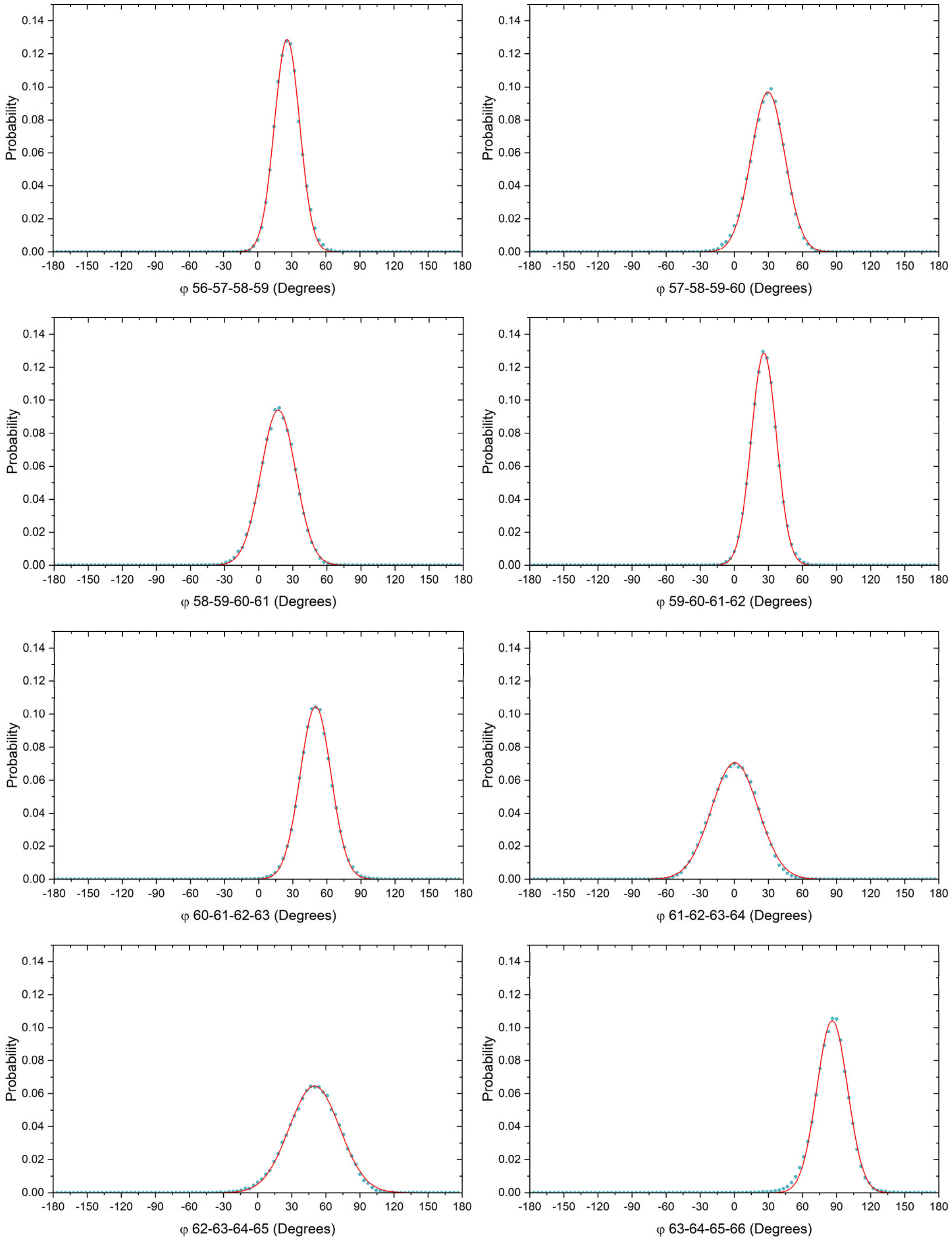

**Fig. S42. Analysis of the BB dihedrals in the TMH3/ND6 π-bulge region.**
Distribution of the backbone dihedral angles observed throughout the simulations in the first replica of m3H$^{op}$ system (pine dots). The Gaussian curves used to fit the dihedral angles distribution and the sum of the fitting Gaussians are in red.

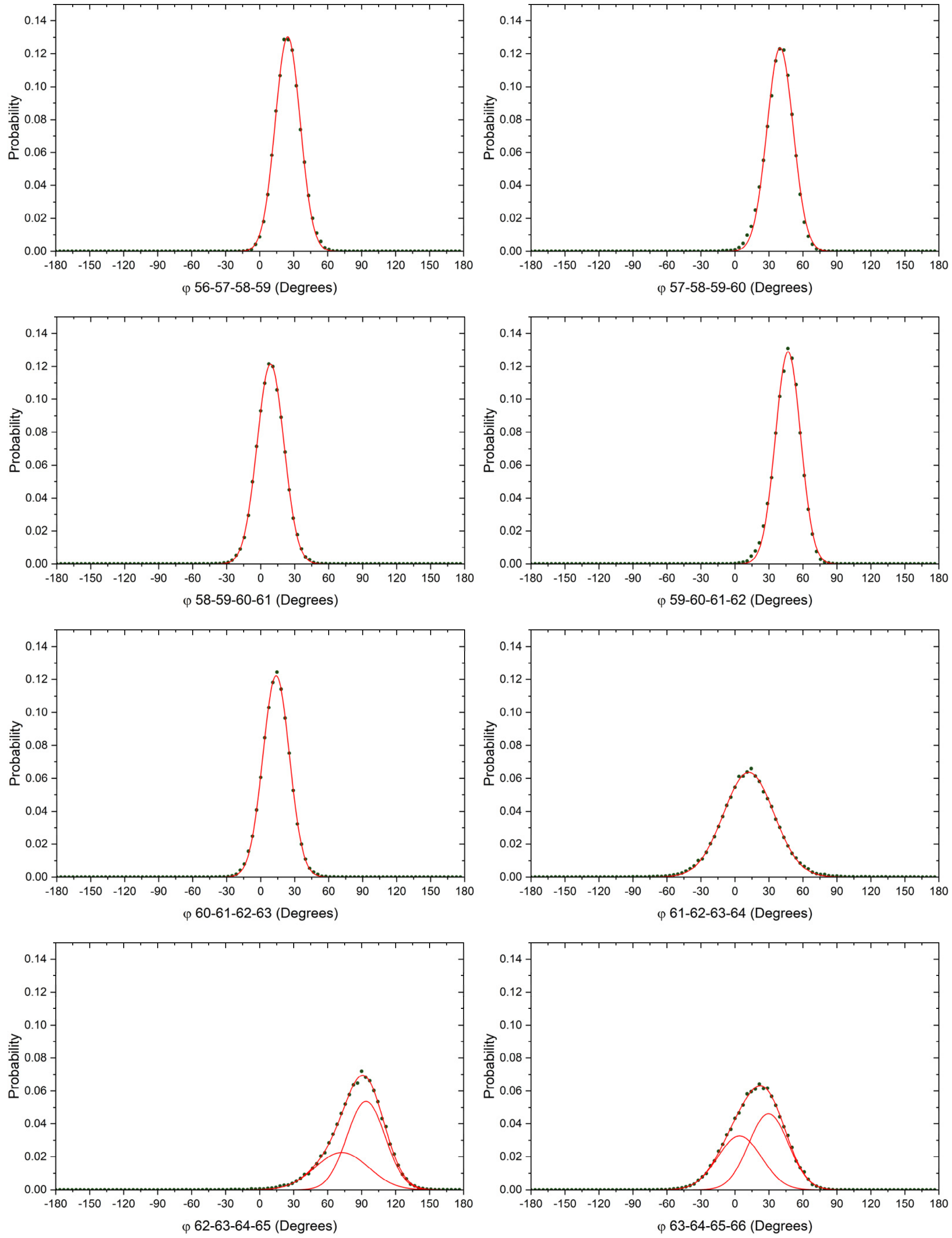

**Fig. S43. Analysis of the BB dihedrals in the TMH3/ND6 π-bulge region.**
Distribution of the backbone dihedral angles observed throughout the simulations in the second replica of m3H$^{op}$ system (dark lime dots). The Gaussian curves used to fit the dihedral angles distribution and the sum of the fitting Gaussians are in red.

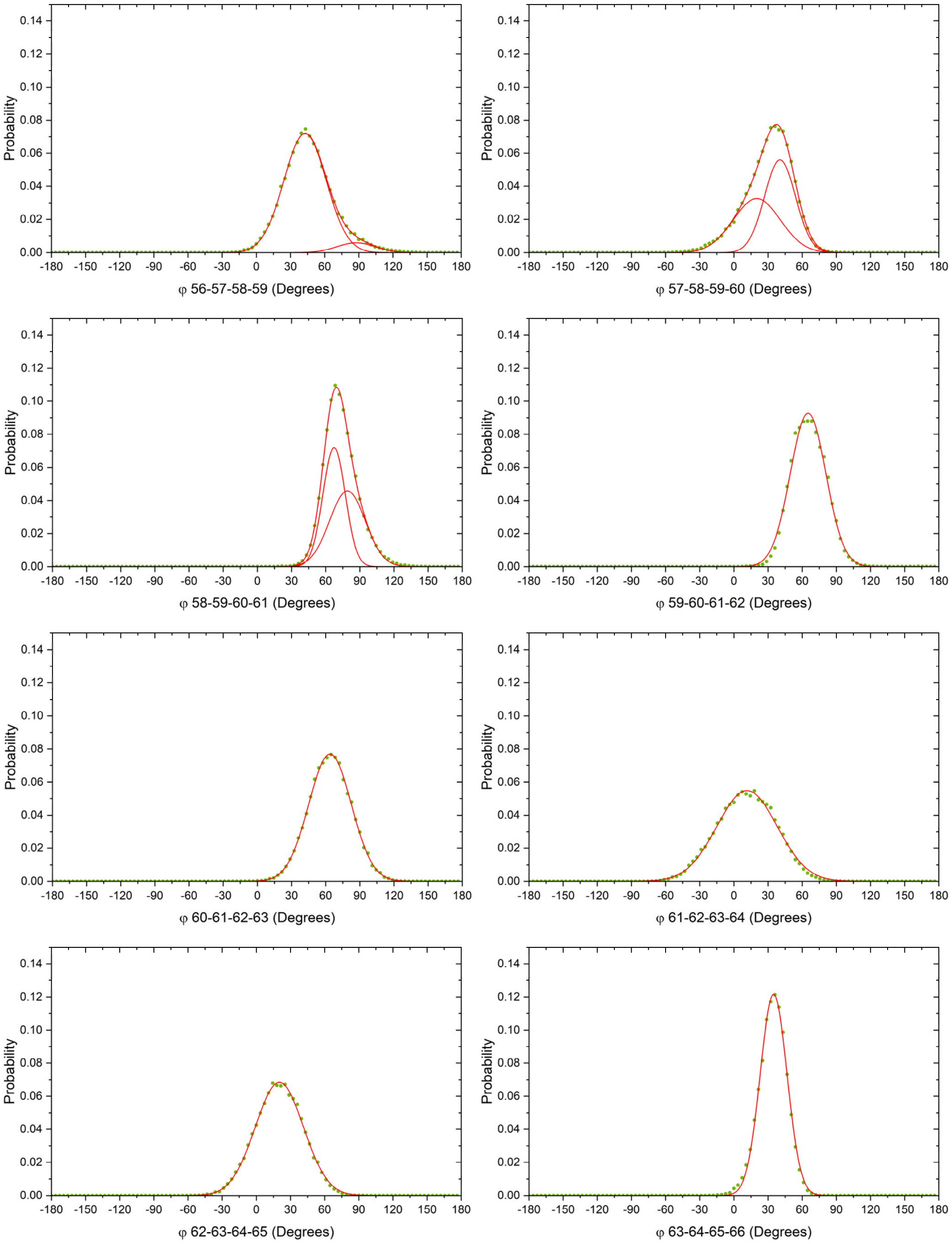

**Fig. S44. Analysis of the BB dihedrals in the TMH3/ND6 π-bulge region.** Distribution of the backbone dihedral angles observed throughout the simulations in the third replica of m3H$^{op}$ system (lime dots). The Gaussian curves used to fit the dihedral angles distribution and the sum of the fitting Gaussians are in red.

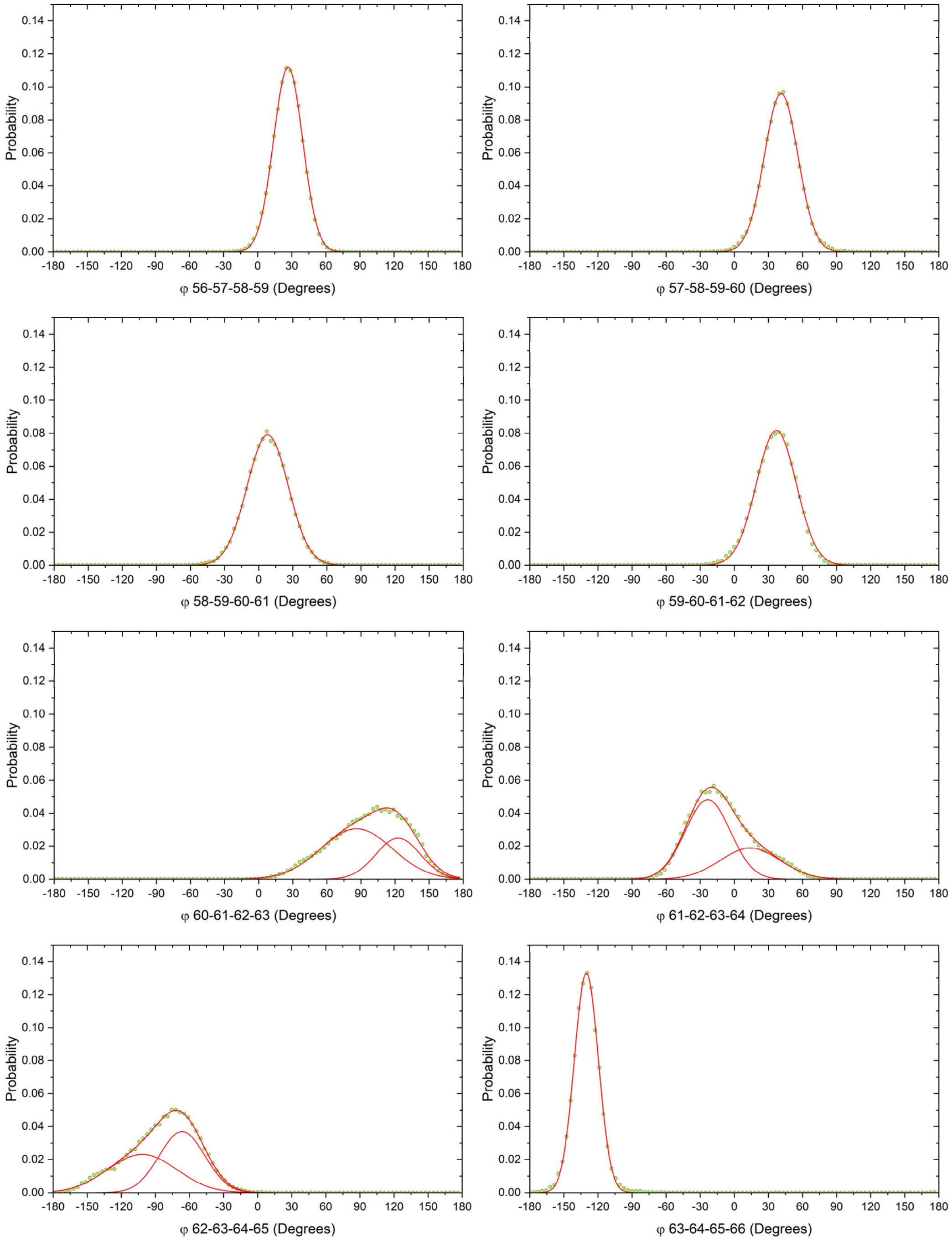

**Supplementary Methods**

Combined Essential Dynamics on the time evolving BB distance matrix.
The procedure applied here is the one proposed by Berendsen and co-worker in 1995 (*49*) and recently described by Palma and Pierdominici-Sottile (*50*), also known as "Combined Essential Dynamics".
A total trajectory is created by concatenating $n = 9$ individual simulations of distance matrix network: 3 runs for the *wild type* + 3 runs for the single mutant + 3 runs for the triple mutant, for a total of 270,000 points and 820 observables (pairwise distances between backbone beads, BB).
The covariance matrix of the concatenated trajectory is given by the average of $n=9$ individual covariance matrices plus the covariance matrix of the individual average structure, as follows:

$$\boldsymbol{C}^{(cn)} = \frac{1}{9}\sum_{k=1}^{9} \boldsymbol{C}^{(k)} + \boldsymbol{S}^{(cn)}$$

Where:

$$\boldsymbol{S}_{ij}^{(c9)} = \frac{1}{9}\sum_{k=1}^{9}\left(\langle x\rangle_i^{(k)} - \langle x\rangle_i^{(c9)}\right)\left(\langle x\rangle_j^{(k)} - \langle x\rangle_j^{(c9)}\right)$$

Here $x_i$ is the *i-th* distance between a couple of BB beads.
The total covariance matrix is then diagonalized in order to get:

$$\boldsymbol{C}^{(cn)}\boldsymbol{v}_i = \lambda_i \boldsymbol{v}_i$$

for $i = 1{,}820$, where $\boldsymbol{v}_i$ is the *i-th* eigenvector and $\lambda_i$ is the *i-th* eigenvalue.
The projections along the first three eigenvectors are calculated as follows:

$$c_i(t) = x_i(t) * \boldsymbol{v}_i, i = 1{,}3$$

The first three projections account for more of the 60% of the total variance.
The time-lagged system variance can be calculated as:

$$variance(\tau) = \langle |c_i(t+\tau) - c_i(t)| \rangle$$

Contact matrix frequency.
The side-chains distance matrix is defined as $\boldsymbol{D} = [d_{ij}]$, where $ij$ are the element of the matrix is the distance $d_{id}$ between the centers of the side-chain beads of residues $i$ and $j$.
Contact matrix $\boldsymbol{C} = [c_{ij}]$, with elements $c_{ij}$ defined as:

$$c_{ij} = 1 \text{ if } d_{ij} \leq d_{cut-off}$$
$$c_{ij} = 0 \text{ if } d_{ij} > d_{cut-off}$$

Here $d_{cut-off}$ is the cut-off distance defining residue being in contact. Here $d_{cut-off}$ was set equal to 0.6 nm, representing the distance between side-chains center of mass.

Water analysis.

For the analysis for the putative water insertion in the ND6 TMH3 π-bulge region a tool based on the python package MDAnalysis (*71*) was used. The analysis is based on approximating the space between the ND6 TMH3 and the adjacent helices in the ND3 and ND4L subunits with two cylinders of fixed diameter (0.8 and 1.0 nm, respectively). The procedure consists of four steps that are applied to each frame: *i*) the beads defining the channel and the principal axis of the channel are found; *ii*) the simulation box is rotated to align the principal axis of the channel with the z-axis of the reference system; *iii*) the "Center Of Geometry" (COG) of the regions that are approximated by the two cylinders are found. Only backbone beads are included in the definition of the COG. Finally, *iv*) the water molecules are searched in the channel region, which is defined by two cylinders built using the *cyzone* function included in MDAnalysis. The first two steps are necessary because the *cyzone* function selects a cylindrical region with the axis parallel to the z-axis of the reference system. The result of this analysis is the number of water molecules in the channel at each frame and in all the studied systems only few water molecules stick to the external side of the cylinders, while no water molecules enter the channel.

Coarse-grained modelling of the mutated residues.

To illustrate what is shown by the bar diagrams in Figs. 5 and 6, it is necessary to consider the structural differences between the *wild type* and mutated residues in the CG model (*35, 36*). Mutate a methionine into a valine does not change the number and type of beads used to represent these residues. In fact, each residue is represented by a bead for the backbone and an apolar (C) bead for the side chain (Martini's bead type C5 and C2 for methionine and valine, respectively. The number highlights the polar affinity, specifically the C5 bead has a larger polar affinity than C2). C5 and C2 beads feature identical Lennard-Jones parameters but different inter-beads interaction parameters. In most of the cases, the C2 bead can form slightly less attractive inter-molecular interactions than C5. The only exception are the interactions with other C2 beads, for which C2 beads are more attractive than C5. Moreover, the distance between the BB and the side chain bead is different between the two residues (0.31 and 0.20 nm for methionine and valine, respectively). In contrast, concerning the p.A71T/ND4L mutation, the mutation of the alanine (represented by a backbone bead alone) with a threonine implies in the addition of a second polar-type bead to simulate the side chain.

Phosphopantetheine CG mapping

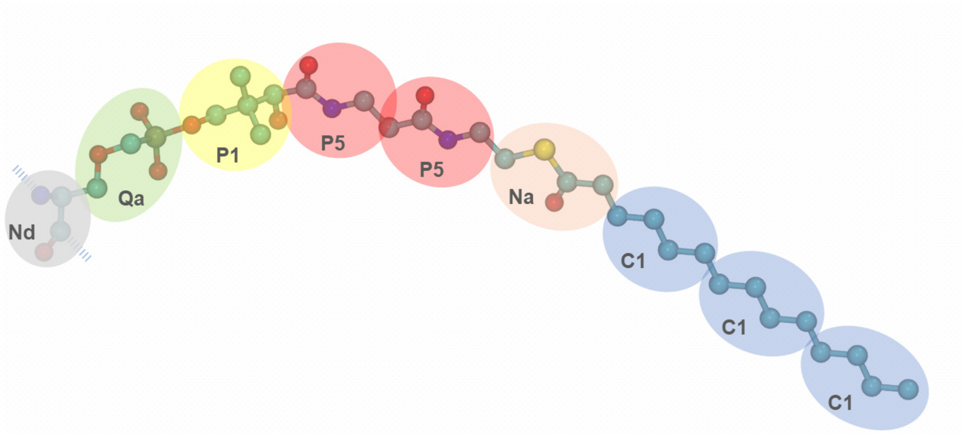


**Fig. S45. CG mappings of the phosphopantetheine (ZMP).**
CG beads are shown over the atomistic structures of the ZMP. The Martini beads are represented with different colors and type names are also reported. 'Nd' grey bead corresponds to the backbone bead of Serine 44 of the Acyl Carrier subunit.

**Table S6. Martini CG Parameters used for the phosphopantetheine prostetic group.**

| Bead type | Charge |
|---|---|
| Nd | 0 |
| Qa | -1 |
| P1 | 0 |
| P5 | 0 |
| Na | 0 |
| C1 | 0 |

The equilibrium bond length ($d$) were all set to $d$ = 0.47 nm, while the force constants ($K_d$) were set to $K_d$ = 1250 kJ nm$^{-2}$ mol$^{-1}$. Equilibrium angles ($\theta_{SSS}$) and force constants ($K_\theta$) were all set equal to $\theta_{SSS}$ = 180 degrees and $K_\theta$ = 25 kJ mol$^{-1}$, respectively.

**Table S7. Parameters used in the five equilibration stages on each system.**

| Stage | Steps | Time step (fs) | Restraints force constants (J $mol^{-1}$ $nm^{-2}$) on protein beads and on lipid heads |
|---|---|---|---|
| 1 | 1,000,000 | 2 | 1000 / 200 |
| 2 | 500,000 | 5 | 500 / 100 |
| 3 | 250,000 | 10 | 250 / 50 |
| 4 | 125,000 | 20 | 100 / 50 |
| 5 | 125,000 | 20 | 20 / 10 |
